# Converging on consistent functional connectomics

**DOI:** 10.1101/2023.06.23.546329

**Authors:** Andrea I. Luppi, Helena M. Gellersen, Zhen-Qi Liu, Alexander R. D. Peattie, Anne E. Manktelow, Ram Adapa, Adrian M. Owen, Lorina Naci, David K. Menon, Stavros I. Dimitriadis, Emmanuel A. Stamatakis

**Author notes:** Correspondence: Andrea I. Luppi. German Center for Neurodegenerative Diseases, Magdeburg, Germany (Helena M. Gellersen). These authors share senior authorship: S.I.D., E.A.S.

## Abstract

Functional interactions between brain regions can be viewed as a network, empowering neuroscientists to leverage network science to investigate distributed brain function. However, obtaining a brain network from functional neuroimaging data involves multiple steps of data manipulation, which can drastically affect the organisation and validity of the estimated brain network and its properties. Here, we provide a systematic evaluation of 576 unique data-processing pipelines for functional connectomics from resting-state functional MRI, obtained from all possible recombinations of popular choices for brain atlas type and size, connectivity definition and selection, and global signal regression. We use the portrait divergence, an information-theoretic measure of differences in network topology across scales, to quantify the influence of analytic choices on the overall organisation of the derived functional connectome. We evaluate each pipeline across an entire battery of criteria, seeking pipelines that (i) minimise spurious test-retest discrepancies of network topology, while simultaneously (ii) mitigating motion confounds, and being sensitive to both (iii) inter-subject differences and (iv) experimental effects of interest, as demonstrated by propofol-induced general anaesthesia. Our findings reveal vast and systematic variability across pipelines’ suitability for functional connectomics. Choice of the wrong data-processing pipeline can lead to results that are not only misleading, but systematically so, distorting the functional connectome more drastically than the passage of several months. We also found that the majority of pipelines failed to meet at least one of our criteria. However, we identified 8 candidates satisfying all criteria across each of four independent datasets spanning minutes, weeks, and months, ensuring the generalisability of our recommendations. Our results also generalise to alternative acquisition parameters and preprocessing and denoising choices. By providing the community with a full breakdown of each pipeline’s performance across this multi-dataset, multi-criteria, multi-scale and multi-step approach, we establish a comprehensive set of benchmarks to inform future best practices in functional connectomics.

## Introduction

The human brain is a remarkably complex system, comprising a large number of regions interacting over time. To address this challenge and obtain insights about distributed brain function and dysfunction, neuroscientists have turned to network science, whereby different parts of the brain can be viewed as nodes in a network, and the statistical relationships between them are used to represent connections between nodes^1–5^. This powerful approach leverages graph theory to quantify key aspects of brain network organisation *in vivo*, illuminating the neurobiological underpinnings of healthy and pathological cognition^6–9^. In particular, resting-state functional MRI (rs-fMRI) is a very popular imaging tool, due to its excellent spatial resolution and wide applicability^10^: being task-free, it can be easily administered even to challenging populations, from in-utero foetuses^11^ to severely injured and even unconscious patients^12–14^. Indeed, aberrant functional connectivity patterns have been observed in many neurological and psychiatric conditions^15–19^.

However, recent studies have highlighted how different analysis workflows can lead to sometimes drastically different conclusions about the same neuroimaging dataset^20^, owing to a vast pool of possible methodological choices which effectively constitute a combinatorial explosion problem^21^. Crucially, such a combinatorial explosion also plagues network analyses of the human brain: even beyond substantial differences introduced by data preprocessing and denoising procedures^22, 23^, a wide variety of approaches have been proposed to derive brain networks from preprocessed functional neuroimaging data. The very definition of nodes in brain networks is controversial: although fMRI voxels have no intrinsic biological meaning, it is well-established based on both functional involvement and lesion studies but also cellular, molecular, and fiber architecture that the brain exhibits biologically meaningful regional organisation, such that voxels can be grouped together into anatomically distinct areas ^21, 24–26^. However, there is yet no consensus on the most appropriate parcellation of the human brain, or the number and spatial extent of brain regions, or whether they should be discrete or overlapping^27^. Similar difficulties arise for the definition of functional connections (edges) between nodes: how to quantify them, which ones to retain for analysis, and whether to emphasise the presence/absence of connections (binary network) or their relative strength (weighted network)^18, 28^, highlighting the intricacies of this issue^27^. This challenge has practical consequences: even with high-quality data, a poor choice of network construction pipeline may produce misleading conclusions about neurobiology and functional organisation, and possibly misinform biomarker discovery and clinical practice. Thus, to ensure the value of graph-based estimates as clinical biomarkers, it is of paramount priority to establish what is the most appropriate way to construct a functional brain network from rs-fMRI data.

Reliability of network topology is of fundamental importance for any subsequent analysis of network properties: any pipelines that recover vastly different topologies from two scans of the same individual taken within the same hour, are liable to produce misleading results when used to associate network properties with behavioural traits^10^ or clinical outcomes^29^. Thus, identification of reliable network construction pipelines represents a fundamental prerequisite for both network-based investigation of individual differences using functional neuroimaging^30, 31^ and subsequent efforts aimed at clinical translation^32^. Existing scientific work comparing different network construction steps typically focused on specific global or local network properties (e.g., modularity, small-world character, global or local efficiency, down to individual edges) and evaluated the different alternatives by maximising the intra-class correlation (ICC) of the adopted global or local network properties^25, 33–43^.

However, these approaches both have limitations. On the one hand, focusing on local aspects (individual edges, node-level properties) runs the risk of “missing the forest for the trees”^33^, because networks are more than just collections of edges: rather, the way that edges are organised gives rise to micro-, meso- and macro-scale structure, which is precisely what makes network-based approaches so powerful. On the other hand, focusing on specific high-level properties of the network will inevitably limit the generalisability of results, because a vast and ever-growing array of network properties can be defined and used to obtain insights about brain function^44, 45^, but there is no guarantee that recommendations pertaining to one will also apply to others.

In the present study, we introduce a framework to explicitly address and tame the combinatorial explosion. First, we evaluate network construction pipelines end-to-end, rather than restricting our attention to individual steps in isolation, as most previous studies have done. Second, we base our evaluation on the network’s topology, that is, the network’s organisation as a whole. For this purpose, we leverage the advantages of the recently introduced “Portrait divergence” (PDiv) measure of dissimilarity between networks^46^. This information-theoretic measure simultaneously takes into account all scales of organisation within a network, from local structure to motifs to large-scale connectivity patterns. Therefore, it incorporates all aspects of network topology, enabling us to go beyond the use of specific and arbitrarily-chosen graph-theoretical properties.

Third, test-retest reliability is a necessary but arguably not sufficient condition for a pipeline to be suitable for functional connectomics. Therefore, we seek to identify network construction pipelines that minimise spurious (noise- or motion-induced) differences between brain networks of the same individual across repeated scan sessions, but that also satisfy additional criteria of biological relevance: sensitivity to individual differences, clinical contrasts of interest, and experimental manipulations – here operationalised by pharmacological intervention with the general anaesthetic propofol. Fourth, to ensure the generalisability of our results^29^, each pipeline is evaluated across two independent test-retest datasets, spanning short (45 minutes), medium (2-4 weeks) and long term delays (5-16 months). Our focus here is not on preprocessing/denoising approaches to fMRI data (where a vast literature exists^47–50)^, but rather on the workflow that *begins* with preprocessed fMRI data and results in a brain network. However, to ensure that our recommendations can be further generalised to datasets acquired with different scanning parameters and preprocessed with different methods, we also require that optimal pipelines should meet all the above-mentioned criteria in an additional independent dataset (test-retest dataset from the Human Connectome Project), which was acquired with higher spatial (2mm) and temporal resolution (TR=0.72s) than the other datasets; preprocessed using a surface-based rather than volume-based workflow; and denoised with a different method than the anatomical CompCor used for our main datasets (FIX-ICA, which is designed to affect artifacts specifically and avoid modifying the neural signal of interest)^47, 51, 52^.

Through this comprehensive, multi-criterion approach, we compare the topologies of functional brain networks obtained from systematic combinations of different options at each step in the network construction process. (i) First, given our interest in robustness and generalisability, we conduct all our analyses on two versions of the same data: with versus without the controversial preprocessing step of global signal regression (GSR) ^53^. This allows us to make recommendations that are specific for GSR-processed data, or for non-GSR-processed data, as well as identifying network processing pipelines that are suitable for both. (ii) Definition of network nodes: based on functional characteristics (combination of local homogeneity and global gradients of connectivity), or anatomical properties, or multimodal features from functional and structural MRI ^21, 24^. (ii) Number of nodes: approximately 100, 200, or 400, for each type of parcellation. (iv) Two different ways to define network edges from BOLD time-series: linear Pearson correlation or non-linear mutual information. (v) Eight different approaches to filter the network’s edges: by imposing a pre-specified density (retaining 5%, 10%, or 20% of total edges, or matching the density of the structural connectome), or imposing a pre-specified minimum edge weight (0.3 or 0.5), or using data-driven methods (Efficiency Cost Optimisation and Orthogonal Minimum Spanning Trees, two different strategies to define and then optimise the balance between network efficiency and wiring cost)^54–56^. (vi) Use of either binary or weighted networks. Fig.1 illustrates the set of choices across the investigated network construction steps that influence the construction of a functional brain network, yielding a total set of 576 pipelines (2*3*3*2*8*2). To the best of our knowledge, this is the first time in the literature that all of these available network construction steps are explored simultaneously end-to-end, and with a focus on topology as a whole rather than on specific network features.

Overall, a strength of our current study is our ability to make recommendations for the choice of pipelines end-to-end, not only on the basis of theoretical gold standard metrics (test-retest) but also on the basis of practical relevance: meaningful inference about changes in brain functional network topology and individual differences, and robustness and generalisability. To anticipate our main findings, we discovered large and systematic variability among pipelines’ ability to recover a reliable network topology, with the majority of pipelines failing to meet at least one criterion. Choice of an inappropriate network construction pipeline can lead to results that are not only misleading (statistically significant in the opposite direction as the true effect), but *replicably* so (being observed in two independent datasets). However, we also identified a number of pipelines that satisfy all our criteria, in all four test-retest comparisons, making them suitable candidates for functional connectomics and biomarker discovery. Through this multi-dataset, multi-criteria, multi-scale, and multi-step approach, we provide a comprehensive set of benchmarks for trustworthy functional connectomics.

**Fig. 1.**
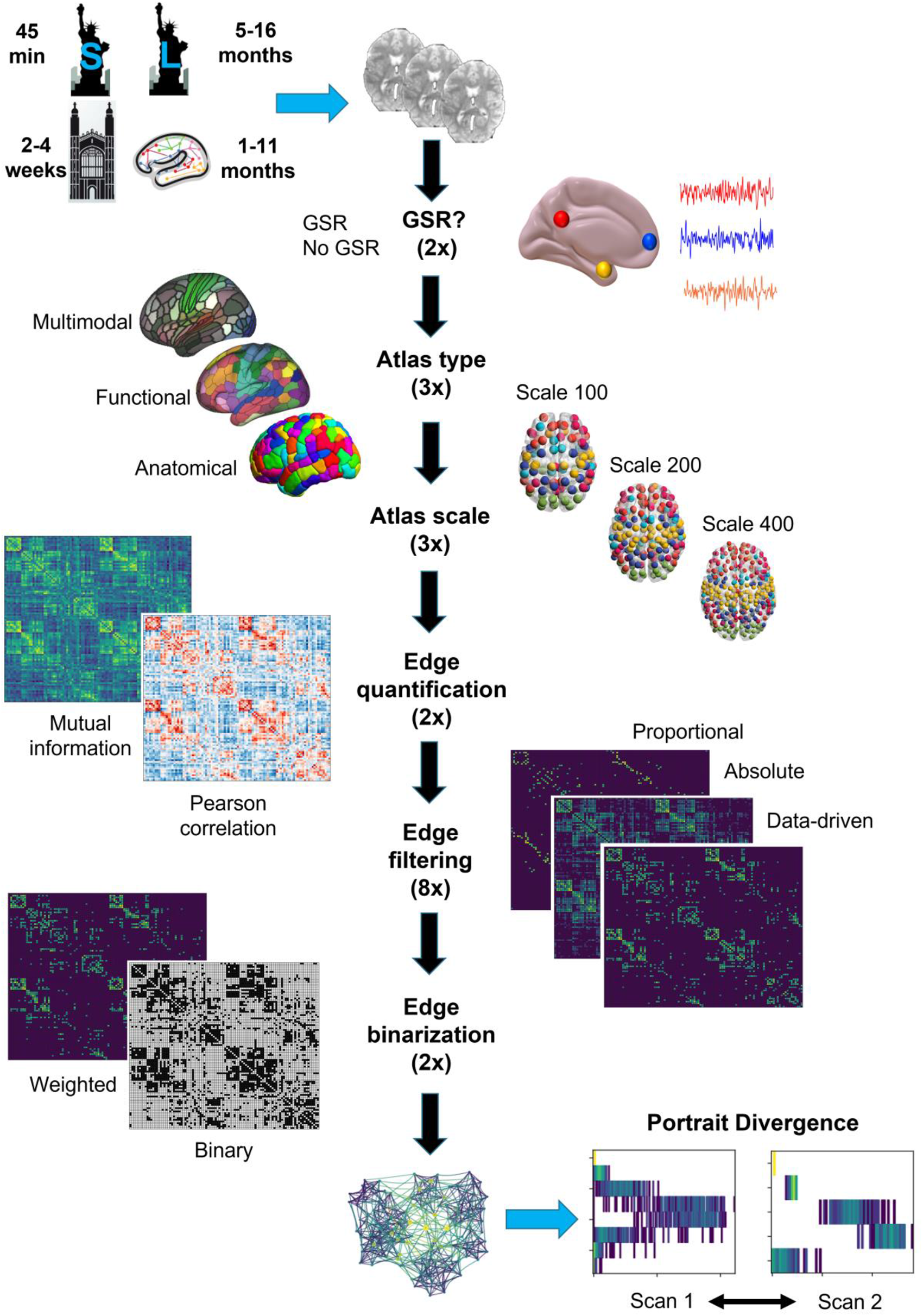
Overview of the steps to turn functional MRI data into a network. Starting from preprocessed and denoised data, the following steps are involved. (i) Use of data with vs without global signal regression (GSR), in addition to other denoising protocol (aCompCor for NYU-short, NYU-long, and Cambridge datasets; FIX-ICA for HCP); (ii) Definition of nodes (based on anatomical features, local and global functional characteristics, or multimodal features); (iii) Choice of number of nodes (approximately 100, 200, or 400); (iv) Definition of connectivity measure (from Pearson correlation or mutual information); (v) Choice of edges to retain (8 filtering schemes considered, based on *a priori* choices of network density, or minimum edge weight, or data-driven strategies to optimise the balance between network efficiency and wiring cost), (vi) Use of binary or weighted edges. In total, we consider 2*3*3*2*8*2 = 576 unique pipelines. For each pipeline, the resulting functional networks are compared for the same subject across different time-spans (minutes, weeks, or months) using the Portrait Divergence. A network portrait for a binary network is a matrix B whose rows each correspond to a histogram obtained by thresholding the matrix of shortest paths between the networks’s constituent nodes, at each path length *l* between 0 and the network’s diameter *L,* such that entry B*_l,k_* encodes the number of nodes that have *k* nodes at distance *l*. For weighted networks, the histogram is obtained by binning (see Methods). Illustration of Portrait Divergence adapted from Bagrow and Bollt (2019)^41^.

## Results

We used an information-theoretic measure of distance between network topologies across scales, termed Portrait divergence (PDiv), to systematically compare 576 alternative network construction pipelines in terms of their ability to recover similar brain network topologies from functional MRI scans of the same individual across minutes (NYU dataset, same-session scans), weeks (Cambridge dataset), or months (NYU dataset, between-sessions comparison) (see Methods, and Fig. S1-2 for examples of network portraits and their divergence). Additionally, we considered an additional dataset (HCP test-retest) that was acquired with higher spatial (2mm isotropic) and temporal resolution (0.72s TR); with longer duration (1200 volumes); denoised using FIX-ICA instead of aCompCor; and parcellated on the surface rather than in volumetric space, as for the other datasets ^57–59^. Our end-to-end approach allowed us to simultaneously assess the effects of atlas type and number of nodes; connectivity quantification, thresholding, and binarisation; and global signal regression; while ensuring robustness to aspects such as acquisition, time between test and retest, and denoising method.

Being grounded in information theory, the Portrait divergence between two networks can be interpreted as measuring how much information is lost when using one network to represent another: it ranges from 0 (no information loss) to 1 (complete information loss) ^46^.

To identify suitable pipelines, we required each of the following criteria to be met:

● **Criterion (I):** Avoiding spurious differences (“PDiv ranking”). Since the two networks that we consider are derived from different scans of the same healthy individuals under conditions in which no experimentally meaningful changes in functional network topology are expected, we aim to identify pipelines that minimise test-retest PDiv. We consider pipelines as candidates for optimal if they are in the top 20% in terms of the average PDiv rank calculated across all test-retest intervals.
● **Criterion (II):** Detecting true experimental differences (“propofol”). Suitable pipelines should detect a significant effect for propofol, in the right direction, in both propofol datasets, i.e., a pipeline is excluded if it fails to detect the expected effect (greater change between wakefulness and anaesthesia than between two awake scans) in either of the two propofol datasets.
● **Criterion (III):** Detecting inter-individual differences (“within-between”). A pipeline fails this criterion if the resulting networks are more similar between than within subjects more than 50% of the times, for any of the four test-retest datasets.
● **Criterion (IV):** Avoiding motion-induced differences (“motion”). A pipeline fails this criterion if its PDiv has a significant correlation with differences in head motion in any of the four test-retest datasets.
● **Criterion (V):** Non-empty networks. As a final sanity check, we also exclude any pipelines that remove all connections from a network, in any of the four test-retest datasets.

These criteria also incorporate the need for recommendations to be generalisable across datasets and acquisition/preprocessing choices, since we only consider a criterion to be met if it is met in *all* the relevant datasets.

A summary of all pipeline characteristics can be found in the Supplementary Interactive Tool. We provide an Excel spreadsheet with an interactive table, including filters that allow selection based on multiple criteria at once to identify pipelines which adhere to the specific criteria desired by the reader. We encourage readers to view the interactive table concurrently with the results described below, as this will allow a closer inspection of associations between a pipeline’s specific network processing choices and the desirable properties described in each subsection of the Results. A user guide for the interactive table is also included in the Supplementary Material.

### Portrait Divergence identifies drastic and systematic variability across pipelines’ capacity to avoid spurious differences

For each dataset, Fig. 2 illustrates the distributions of group-mean test-retest similarities of network topologies (portrait divergence) across the full set of 576 pipelines (See Fig.S3-30 for the distribution of PDiv across pipelines, broken down by network construction step, for each dataset). Clearly, two patterns can be observed. First, network construction pipelines differ widely in how well they are able to recover the same network topology across different scans of the same individual, on average – whether on a timescale of minutes, weeks, or months. The worst pipelines induce a greater than five-fold increase in topological dissimilarity (PDiv) between functional connectomes of the same individual, compared with the best-performing ones.

**Fig.2.**
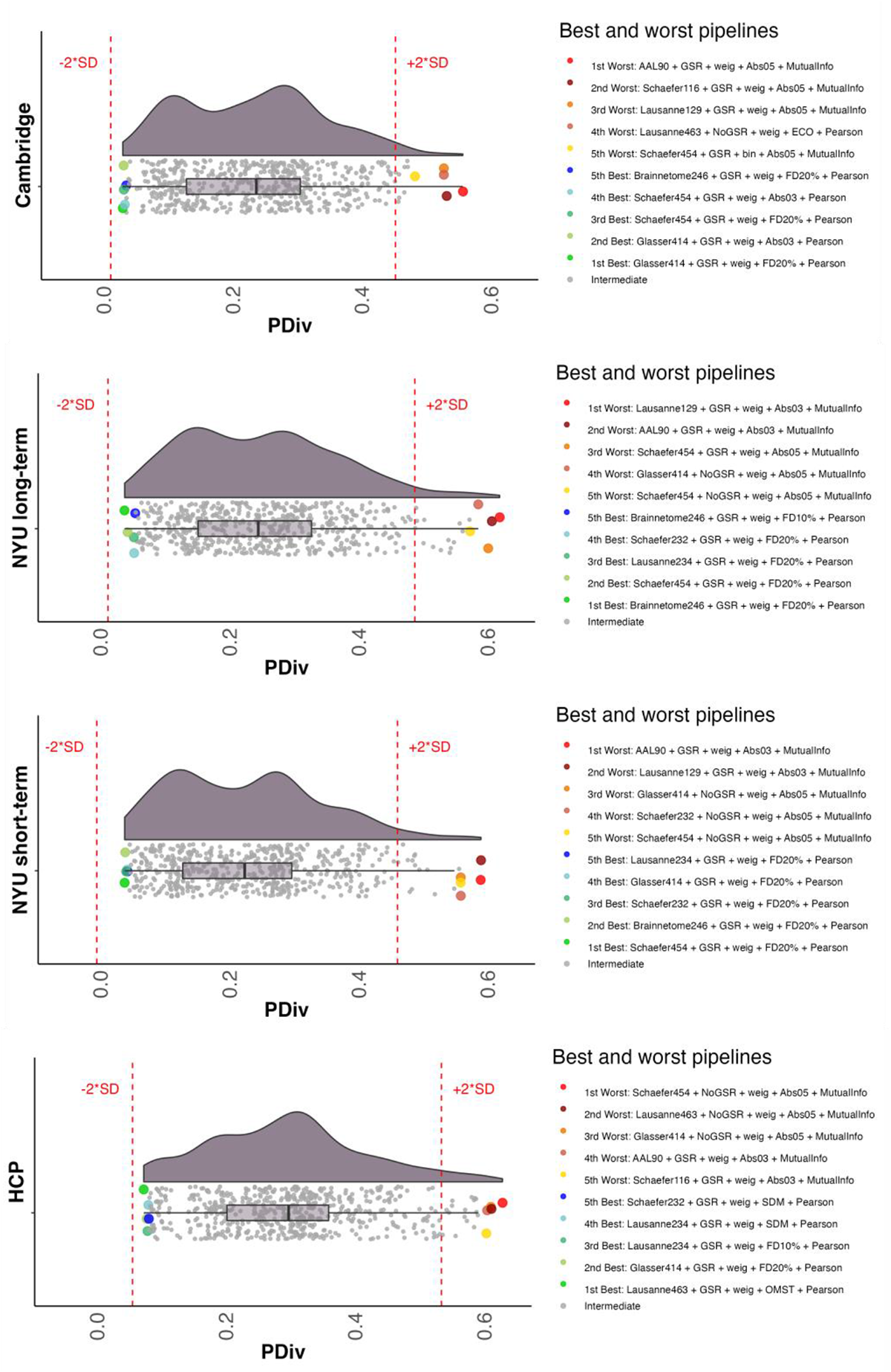
Distribution of group-average portrait divergence values for each of 576 alternative network construction pipelines, across different time intervals. From top to bottom: Cambridge dataset (rescan within 2-4 weeks).NYU short-term dataset (rescan within 45 minutes). NYU long-term dataset (rescan within 16 months; average 11.4); HCP dataset (rescan 1-11 months). Right-side: highlighting the top 5 (lowest PDiv) and bottom 5 performers (highest PDiv). Red lines mark 2 standard deviations from the mean of the distribution. center line, median; box limits, upper and lower quartiles; whiskers, 1.5x interquartile range. **Abbreviations**. GSR: Global Signal Regression. OMST: Orthogonal Minimal Spanning Trees.

Second, our results indicate high consistency across the four test-retest comparisons considered here, in terms of which data-processing steps feature prominently among the pipelines that are best (and worst) at minimising the average within-subject PDiv. Correlation between all pipelines’ ranks across time intervals revealed very high consistency between all datasets (Spearman’s rho ranging from .71 to .98, all p < 0.001) (Fig.3), indicating that pipelines’ suitability for network construction is not dataset-specific but rather can generalise to independent groups of individuals – spanning time intervals from hours to months. We view a small PDiv in these datasets as a desirable property: even though learning and plasticity could account for some amount of connectome reorganisation over weeks or months in healthy adults, such factors cannot plausibly be expected to be the cause of any network-wide reorganisation observed within the course of a single hour (in the absence of any intervention), which should instead be treated as unwanted noise.

**Fig. 3.**
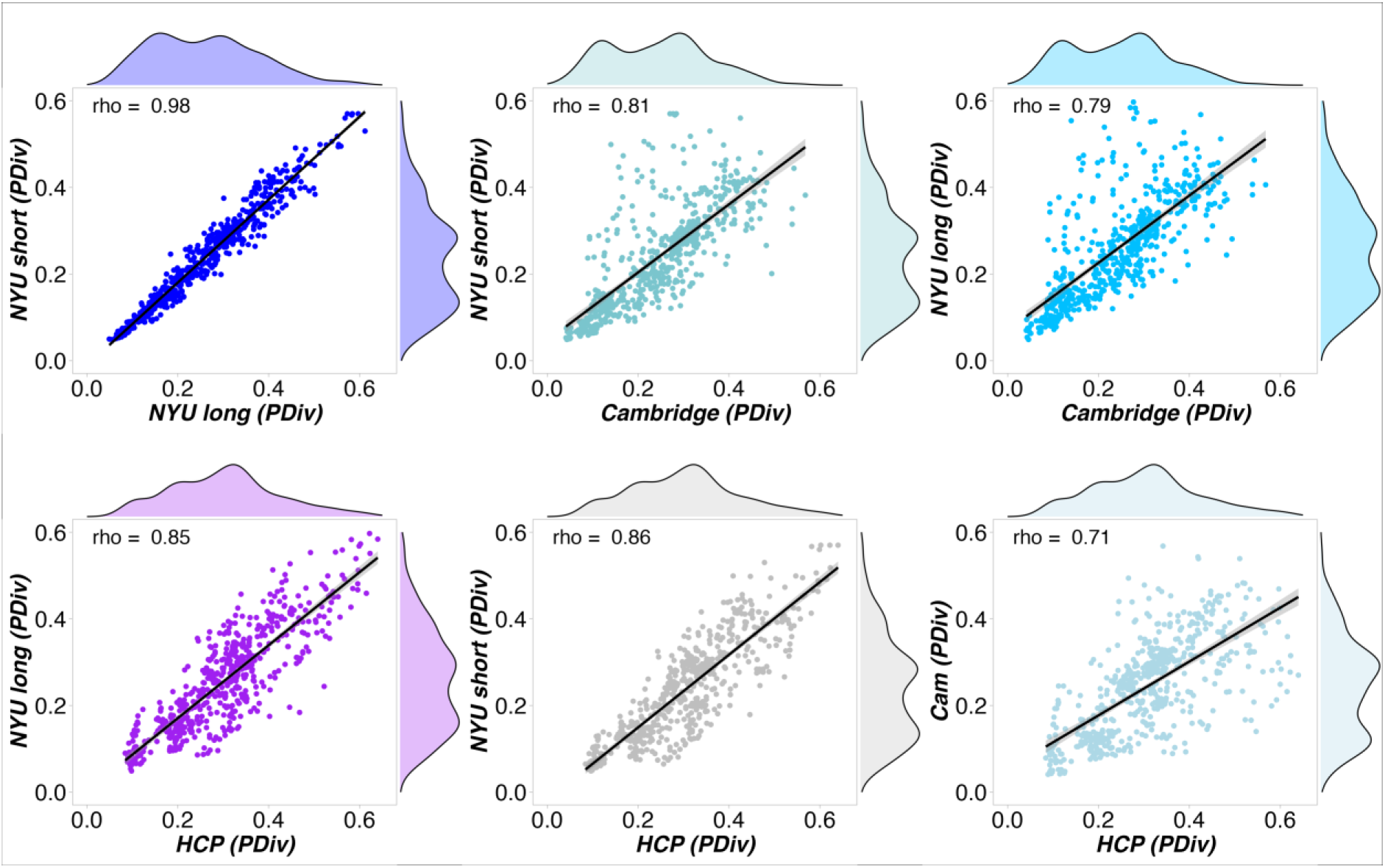
Rank-based correlations of the pipelines’ performance across datasets. PDiv, portrait divergence; HCP, Human Connectome Project data; NYU, New York University dataset. All p < 0.001.

### Sensitivity to experimental differences: Low-PDiv pipelines are more likely to detect pharmacologically-induced connectome reorganisation

We have shown that network construction pipelines vary drastically and systematically in their robustness to noise-induced changes in the functional connectomes of the same individuals scanned multiple times. However, this minimisation of noise-induced differences should not come at the expense of also minimising meaningful changes in network topology, such as control-patient contrasts (an example of this would be a pipeline that never detects any changes). Rather, a good pipeline should simultaneously minimise noise-induced differences, while remaining sensitive to true ones. In other words, test-retest reliability is not the only criterion that neuroscientists need to consider for their choice of network construction pipelines: ultimately, the resulting networks need to also demonstrate empirical usefulness by providing neurobiologically meaningful results^43, 44^. An ideal pipeline would therefore strike a balance between sensitivity to experimental manipulations or contrasts of interest on the one hand, and low portrait divergence in test-retest over relatively short periods of time in healthy individuals and under the same test conditions on the other hand. Therefore, in addition to identifying pipelines that do not detect differences when we know that there should be none or only minor ones (best exemplified by test-retest scanning within the same hour), we should find pipelines that *can* also detect a difference, when we know that a difference must be present: we need to combine a low rate of false positives (low test-retest PDiv) with a low rate of false negatives.

Perhaps the most drastic possible difference that can be induced between two scans of the same individual, is that between consciousness and unconsciousness. General anaesthetics such as the intravenous agent propofol can rapidly and reversibly induce a state of unconsciousness, whereby the subject is behaviourally unresponsive and has no subjective experiences. There is arguably no short-term, reversible alteration of the mind that is so all-encompassing, and it cannot be expected to leave the functional connectome unaltered. Therefore, if a pipeline is unable to detect anaesthetic-induced differences in the topology of the functional connectome, we can reasonably conclude that it is not sensitive enough for use in network neuroscience.

Following this rationale, we compared the PDiv from the NYU-short dataset (two scans within the same hour) against the PDiv observed between an awake rs-fMRI scan, and a second scan of the same individuals while under propofol-induced general anaesthesia (also acquired within the same visit). We seek to identify pipelines that produce significantly greater PDiv between an awake and an anaesthetised scan of the same individual, than between two awake scans acquired at a comparable distance in time. To ensure the reliability of our approach, we repeat this analysis for two independent datasets of propofol anaesthesia to further bolster the reproducibility and generalisability of our findings.

Across both datasets, our results suggest that pipelines with lower PDiv also tend to have *t*-scores reflective of the expected effect of propofol (Fig. 4), as demonstrated by significant correlations between short-term test-retest PDiv (based on the NYU dataset) and *t*-scores both for the Western (rho=.42, p<.001) and the Cambridge propofol datasets (rho=.26, p<.001). As control test-retest PDiv becomes larger, *t*-scores also seem to become more variable, Reassuringly, we identified multiple pipelines that provide the expected effect in both datasets (Fig. 4, green dots). Intriguingly, however, we also identified a number of large-PDiv pipelines that detect a statistically significant difference between test-retest and anaesthesia, but in the *opposite* direction: that is, greater connectome reorganisation between two awake scans, than between an awake and an anaesthetised scan (Fig. 4, red triangles). In other words, these pipelines produce networks that are actively misleading about what we have strong reason to believe must be the ground truth (because there is a very substantial difference introduced by anaesthesia, reflected in the suspension of the brain’s input-processing abilities and cognitive function more broadly). These pipelines can be found in the Supplementary Interactive Tool (pipelines labelled “Opposite” in the columns *Status Propofol West* and *Status Propofol Cam*). Worryingly, we find that a non-negligible number of pipelines (38) produce the *opposite* of the expected effect for *both* propofol datasets – thereby returning results that are systematically misleading, and highlighting the dangers of an inappropriate choice of network construction workflow. Of note, *all* the consistently misleading pipelines use an absolute threshold; all but two use weighted edges; and 23/38 use mutual information to quantify connectivity. Overall, 55 pipelines show the expected effect for both propofol datasets, thereby satisfying this criterion, whereas 357 pipelines are neutral (failing to detect statistically significant differences in at least one propofol dataset).

**Fig. 4.**
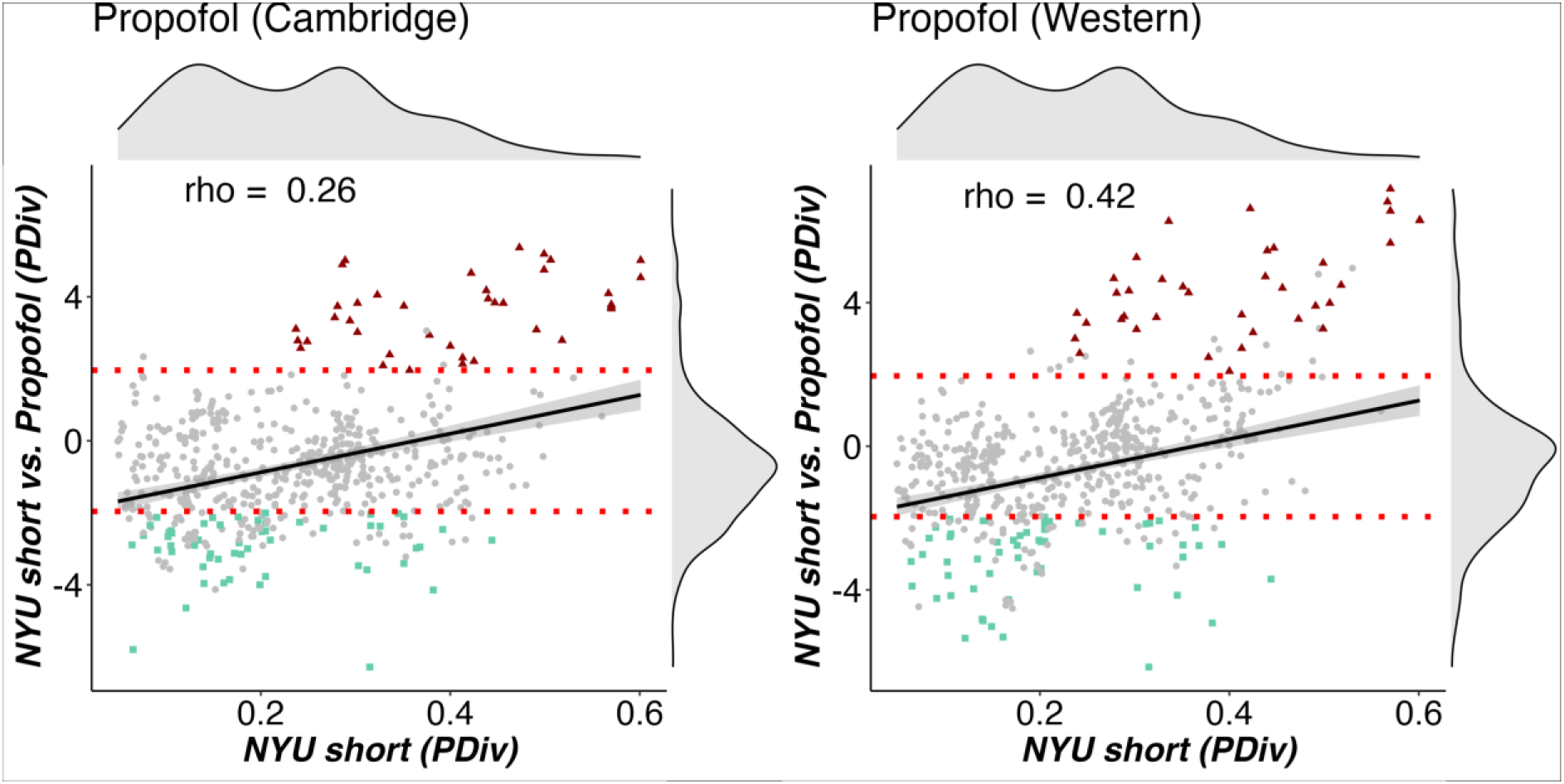
Correlation between low PDiv and ability to detect significant difference between anaesthesia and test-retest. Left: Cambridge anaesthesia dataset, Right: Western anaesthesia dataset. Both correlations are statistically significant, *p* < 0.001. The t-scores are obtained from permutation-based two-sample t-tests comparing PDiv from test-retest NYU short, against PDiv from awake vs anaesthesia. Horizontal red lines indicate t ± 1.96, corresponding to a statistically significant difference between the two groups’ mean, with negative t-scores corresponding to PDiv (anaesthesia) > PDiv (test-retest). Green dots indicate pipelines that produce the expected effect in both datasets. Red triangles indicate pipelines that produce a misleading effect in both datasets.

### Sensitivity to inter-individual differences

Another means by which the adequacy of a pipeline may be assessed is by comparing PDiv within subjects (scan 1 vs. scan 2 for subject 1, scan 1 vs. scan 2 for subject 2, etc…) and PDiv between subjects (subject 1 vs. subject 2, etc…). The proportion of participants for whom the within-subjects (WS) PDiv is smaller than between-subjects (BS) PDiv may be used as an additional criterion of pipeline quality, with the rationale that even after accounting for *bona fide* changes due to plasticity and learning, an individual’s functional connectome should not differ from itself at another point in time, more than it differs from the connectomes of other individuals.

Our results suggest that pipelines with smaller PDiv are also better at producing networks that are sensitive to individual differences, such that the same subject’s brain network diverges less from the same subject’s network than from those of other people. This was the case for the NYU short test-retest data (rho=-.39, p<.001), the medium-term test-retest time interval for the Cambridge dataset (rho=-.42, p<.001), the NYU long test-retest data (rho=- .37, p<.001) and the HCP dataset (rho=-.54, p<.001).

Passing and failing pipelines on the basis of this within-between criterion can be found in the Supplementary Interactive Tool (column *Criterion within-between all*). The interactive tool also lists the *proportion* of participants in a given pipeline for which within-subject PDiv is smaller than between-subjects PDiv in columns *Within-between Cam (%)*, *Within-between NYU short (%)*, *Within-between NYU long (%), and Within-between HCP (%)*. In the Cambridge dataset, 59 pipelines were excluded based on this criterion. This was the case for 43 in the NYU short-term test-retest data and for 32 for the NYU long-term data as well as for 21 pipelines in the HCP data. In total, on the basis of the overall within-between criterion across datasets, 112 pipelines were excluded and 464were retained.

**Fig. 5.**
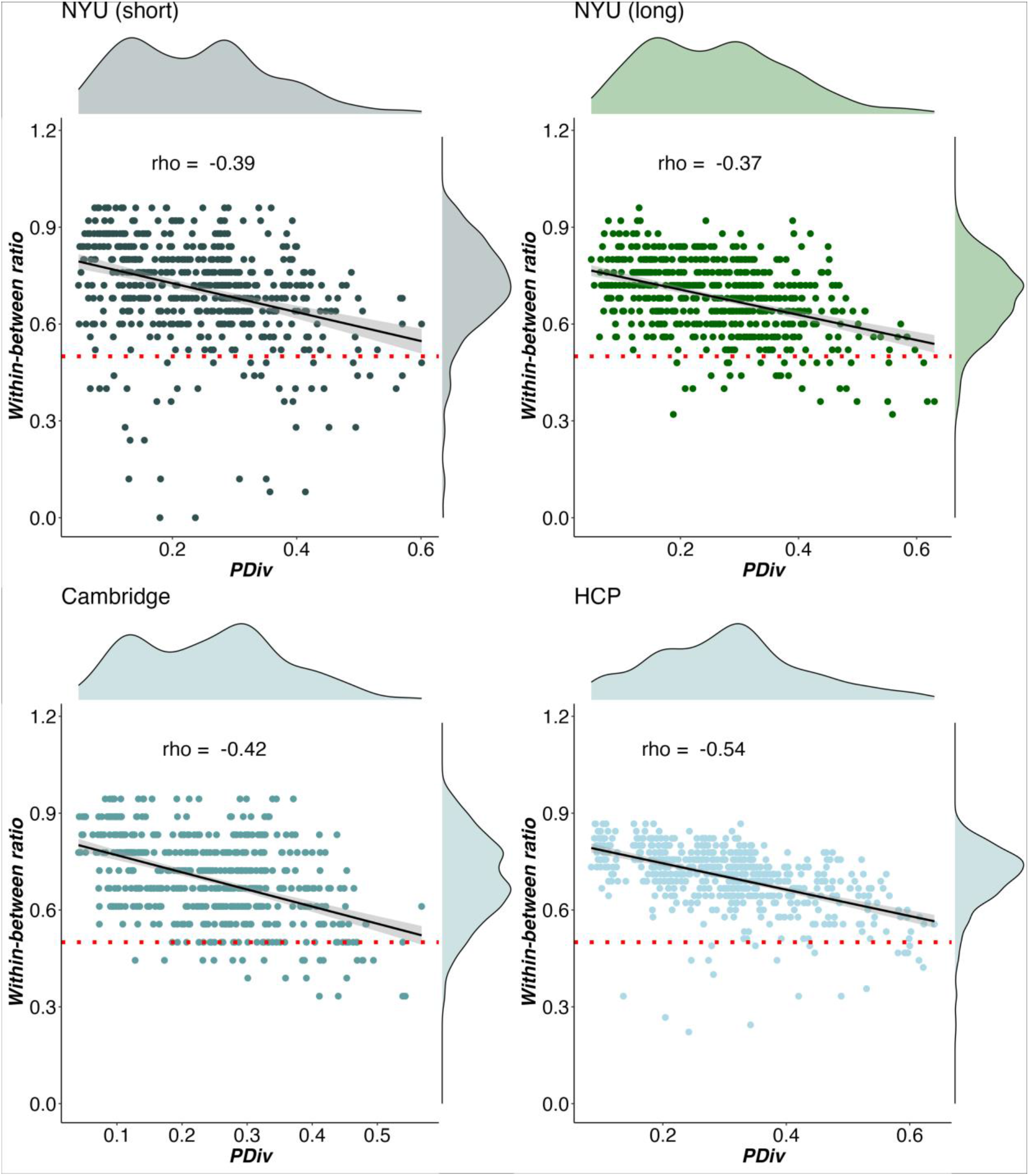
PDiv within versus between individuals. Pipeline PDiv in a given dataset is plotted against the proportion of participants in the same dataset for whom the within-subject PDiv (baseline vs follow up) is smaller than between-subject PDiv. Pipelines above the red line meet the within-between criterion such that portrait divergence is smaller for within-subject test-retest compared to between subject comparisons.

### Avoiding motion confound

As a further criterion, we sought to identify and exclude pipelines whose PDiv is significantly correlated with differences in subject motion (mean framewise displacement). For the Cambridge dataset, 34 pipelines showed a significant correlation between PDiv and motion (magnitude of the correlation coefficient rho ranging between 0.60 and -0.66). For the NYU short-term dataset, 16 pipelines exhibited a significant correlation between PDiv and motion (magnitude of the correlation ranging between 0.45 and -0.57). For the NYU long-term dataset, 13 pipelines exhibited a significant correlation between PDiv and motion (magnitude of the correlation ranging between -0.52 and 0.56). Finally, for the HCP dataset we found that PDiv and motion were correlated significantly in 29 pipelines (rho between -.38 and .41).

It is argued in the literature that GSR can help to mitigate the noise induced by subject motion^42, 45^. When contrasting all pipelines with GSR against those without GSR, no significant difference in the strength of the correlation (absolute r-statistic) between PDiv and motion based on this option was found in the Cambridge (t(568)=.54, p=.592, d=.05), the NYU short test-retest (t(524)=1.44, p=.151, d=.12) or the NYU long-term test-retest (t(524)=1.32, p=.186, d=.11). That is, whether GSR was or was not applied, this decision had no bearing on the degree to which test-retest portrait divergence was associated with motion, on average across all pipelines. However, in the HCP data, there was a significant, moderate effect of GSR on the magnitude of the correlation between motion and PDiv (t(530)=-8.72. p<.001, d=-.76), showing a stronger association between PDiv and motion in pipelines without GSR than with GSR.

### Sanity check: Avoiding empty networks

Pipelines employing an a priori threshold on the strength of edges, rather than on their density (i.e., removing all edges with weight below a pre-specified value, also known as an “absolute” threshold) run the risk of removing *all* edges in the network, if none surpass the threshold value. This would be unquestionably incorrect, but it is conceivable that such an occurrence might never materialise in practice. Indeed, we found that this never occurred when edge weights were defined in terms of Pearson correlation. However, empty networks were returned for at least one subject by a total of 46 unique pipelines employing mutual information for edge weight definition (37 occurrences in the NYU long test-retest, 32 occurrences in the NYU short test-retest, 44 in the HCP dataset, none in the Cambridge dataset). As expected, all of these pipelines used absolute threshold values: mostly with the 0.5 threshold, but for ten pipelines this was also the case for the more lenient 0.3 threshold (reported in the Supplementary Interactive Tool under the column *Criterion edge failure*). Therefore, any pipeline which removes all edges in any one dataset is excluded from further consideration as a suitable candidate. However, note that pipelines that fail this sanity check would also be eliminated from consideration based on the other four criteria: only one of those that failed the sanity check satisfied both the within-between and propofol criteria (Lausanne129 + No GSR + binarisation + Abs 0.3 + Mutual Info).

### Overall recommendations for network construction pipelines

As a final step, we combined all the criteria identified above:

I. Avoiding spurious differences: we operationalise this as having low PDiv (pipelines with the average global rank in the top 20%, as calculated from the average of independent rankings within each dataset; 115 pipelines fulfilled this criterion);
II. Detecting true experimental differences: ability to correctly identify statistically greater PDiv in anaesthesia than test-retest, across *both* propofol datasets (55 pipelines passed);
III. Sensitivity to inter-individual differences: ability to detect smaller within-than between-subjects PDiv in at least 50% of subjects, in each of the four test-retest datasets (464 pipelines passed);
IV. Avoiding motion confounds: no significant correlation between PDiv and subject motion, in any of the four test-retest datasets (444 passed);
V. Non-empty networks: we rejected pipelines that produce empty networks for any subject in any of the four test-retest datasets (530 pipelines pass).

Out of the full set of 576 pipelines considered here, we found that only 8 (around 1%) jointly satisfied all of our criteria in each of the test-retest datasets that we considered – meaning that the vast majority of pipelines (568 out of 576) may be less than optimal (Figure 6 and Table 1). However, 84 pipelines were excluded from the optimal ones because they each failed one single criterion in one single dataset, such that their failures were neither systematic nor pervasive. In particular, the set of optimal pipelines would expand to 26 (∼5% of the total) if a less stringent criterion for the PDiv were adopted, such that all pipelines in the upper 50% were admissible (while still having to satisfy all other criteria in each of the relevant datasets).

**Fig. 6.**
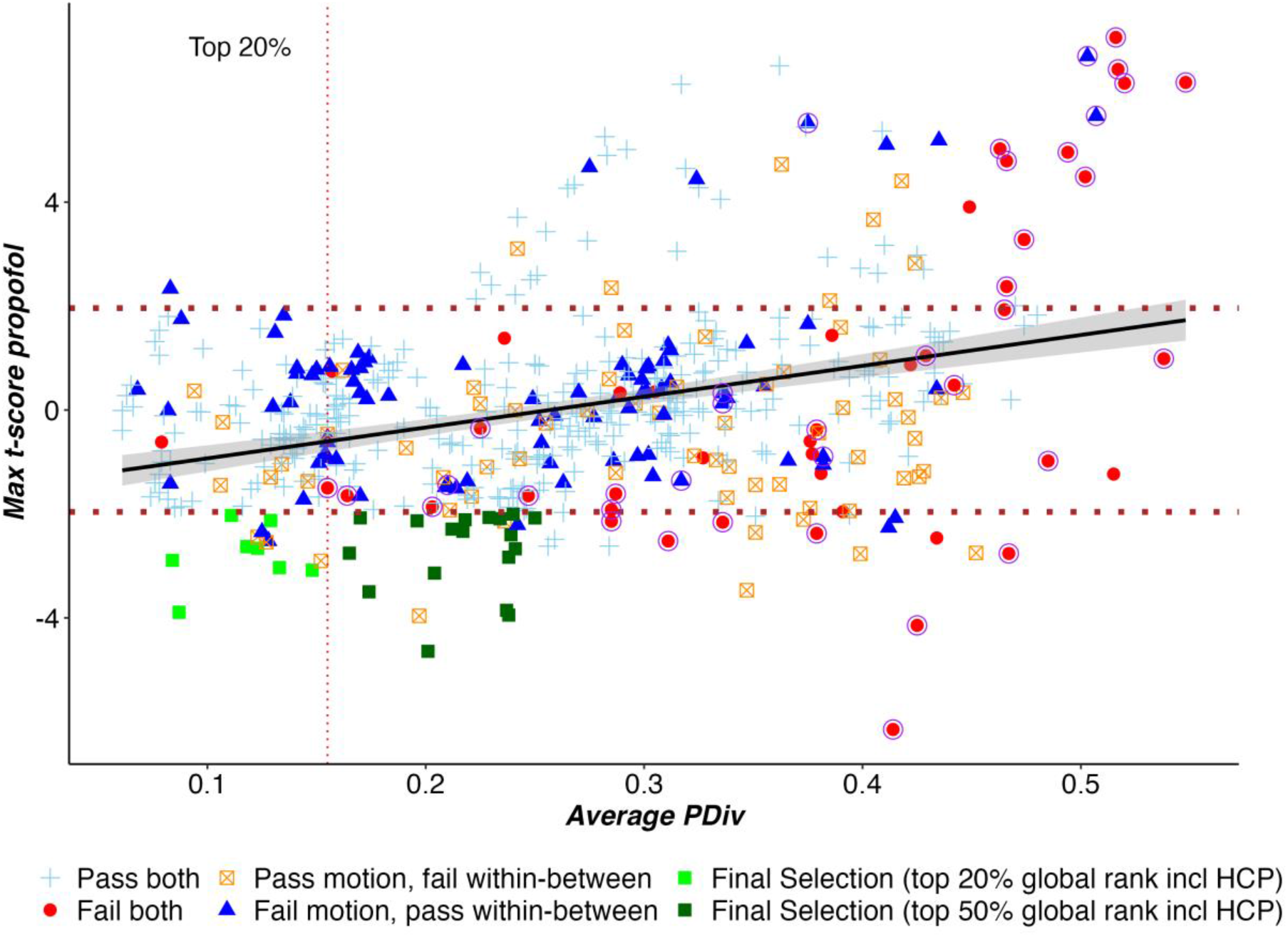
Evaluating pipelines across all criteria. Each data-point represents one pipeline, with colour and shape reflecting which criteria are meet. Criterion (I): Avoiding spurious differences (“PDiv ranking”). We consider pipelines as optimal if they are in the top 20% in terms of the global rank based on PDiv calculated as the average rank achieved in each dataset . We further show pipelines which fulfil all other criteria while being among the top 50% in terms of the average global rank. The maximum average PDiv among the top 50% pipelines is .266. Multiple pipelines were tied for 50^th^ place meaning that a total of 289 pipelines were selected as belonging to the top 50% based on global rank alone. Criterion (II): Detecting true experimental differences (“propofol”). Suitable pipelines should detect a significant effect for propofol, in the right direction, in both propofol datasets, i.e. a pipeline is excluded if it fails to detect the expected effect in either of the two propofol datasets. The Y axis reports the maximum between the two t-statistics obtained for the two propofol datasets, so pipelines satisfy the sensitivity criterion if they score < 1.96 on this axis (i.e., find a significant effect for propofol, in the *right direction*, in *both* propofol datasets). Criterion (III): Detecting inter-individual differences (“within-between”). A pipeline fails this criterion if the resulting networks are more similar between than within subjects more than 50% of the times, for any of the four test-retest datasets. Criterion (IV): Avoiding motion-induced differences (“motion”). A pipeline fails this criterion if its PDiv has a significant correlation with differences in head motion in any of the four test-retest datasets. Criterion (V): Non-empty networks. As a final sanity check, we also exclude any pipelines that remove all connections from a network, in any of the four test-retest datasets. *Fail both* refers to pipelines failing in terms of motion and within-between criteria, while *Pass both* refers to pipelines which satisfy both of these criteria. Points circled in purple represent pipelines that produced empty networks. Overall, 8 pipelines satisfy all criteria in all datasets; this number grows to 26 if a more liberal PDiv criterion is adopted (top 50% global rank).

When considering the distribution of individual pipeline steps among the 8 optimal ones, three clear patterns emerge: all pipelines use weighted (rather than binary) edges, and all quantified connectivity in terms of Pearson correlation (rather than mutual information) (Fig.7). Moreover, the preferred filtering method among the optimal pipelines is the OMST, a method to optimise the balance between network efficiency and wiring cost in a data-driven manner (selected in 5/8 cases). In other words, the single combination of Pearson correlation, weighted edges, and OMST accounted for 5 out of 8 optimal pipelines, despite being only one out of 2*8*2=32 equally likely combinations of edge quantification, thresholding, and binarisation. This is highly unlikely to occur just by chance: the probability of randomly choosing 8 pipelines out of 576 and having 5 or more of them belong to the same group (out of 32 possible groups) is less than 3×10^-4^ (confirmed with permutation testing: *p* < 0.001). In contrast to the clear importance of edge definition, atlas choice seems to have less bearing on a pipeline’s performance, but we do observe greater prevalence of pipelines using GSR than not (six out of eight). However, this last pattern disappears when using a less stringent criterion in terms of PDiv (average rank across datasets in the top 50%, rather than the top 20% – while still satisfying all other criteria). We also find that while edges based on Pearson correlation still dominate under this less stringent criterion, there is now also a number of well-performing pipelines using proportional thresholds (either fixed or SDM) with binarised edges. Node type and number remains less clearly decisive, and the occurrence of GSR versus no-GSR pipelines is nearly equal.

**Fig. 7.**
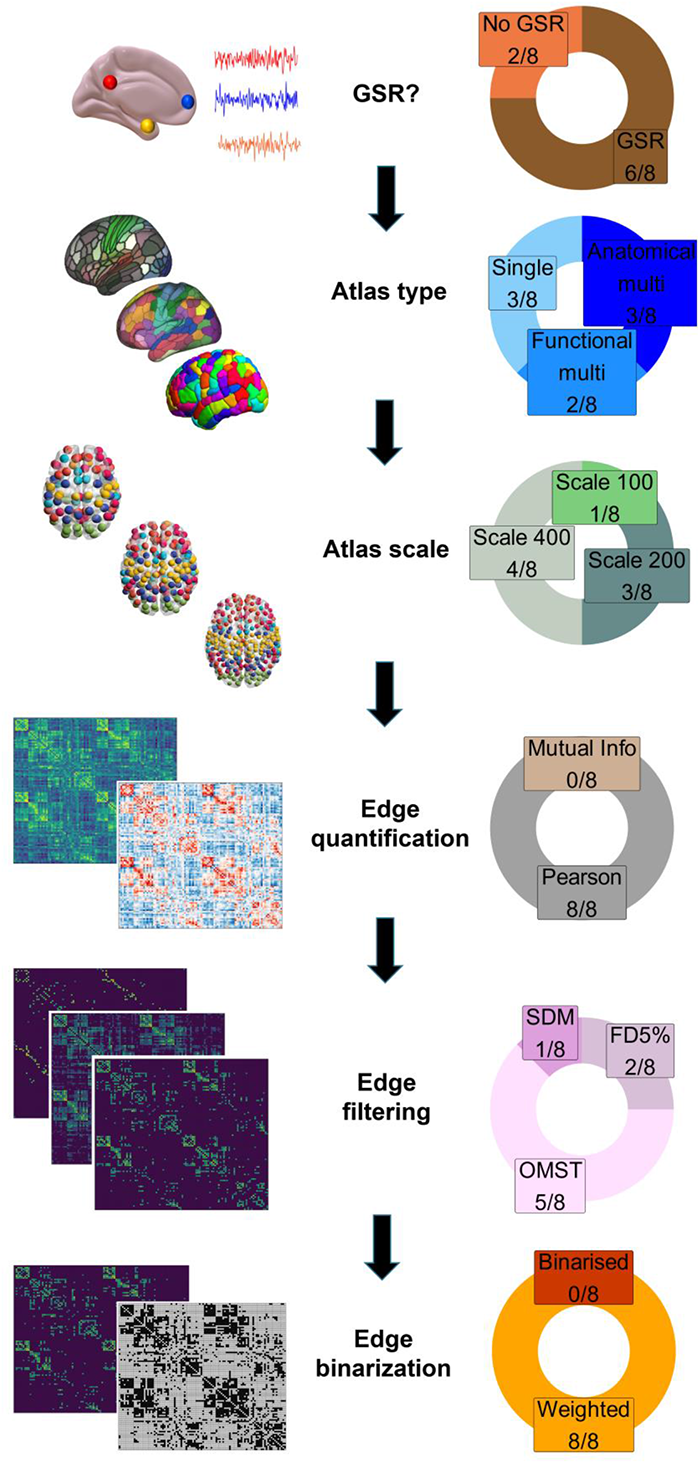
Prevalence of specific network construction steps among the 8 optimal pipelines. Pie charts demonstrate, for each network construction step, the proportion and absolute number of each option that is found among the optimal pipelines. **Abbreviations**. FD: fixed density. GSR: global signal regression. OMST: orthogonal minimal spanning tree. SDM: structural density. See Fig. S31 for a version of this figure with a breakdown of the pipelines under the more liberal PDiv criterion.

**Table 1.** Final selection of pipelines which meet all selection criteria.

| Pipeline | Atlas type | Atlas size | Binarisation | GSR options | Threshold | Edge type | PDiv (Cam) | Rank (Cam) | PDiv (NYUs) | Rank (NYUs) | PDiv (NYUI) | Rank (NYUI) | Global Rank | Average PDiv | NYUs vs. Propofol (Western) | NYUs vs. Propofol (Cam) | Within-between (Cam) | Within-between (NYUs) | Within-between (NYUI) | Within-between (HCP) | Motion $\rho$ (Cam) | Motion $\rho$ (NYUs) | Motion $\rho$ (NYUI) | Motion $\rho$ (HCP) |
| --- | --- | --- | --- | --- | --- | --- | --- | --- | --- | --- | --- | --- | --- | --- | --- | --- | --- | --- | --- | --- | --- | --- | --- | --- |
| Lausanne234 + GSR + weig + OMST + Pearson | Anat | Scale 200 | Weighted | GSR | OMST | Pearson | .11 | 76 | .07 | 18 | .08 | 17 | 32 | .09 | -3.89 | -5.80 | .61 | .68 | .72 | .84 | -.31 | -.08 | -.07 | -.24 |
| Lausanne463 + GSR + weig + SDM + Pearson | Anat | Scale 400 | Weighted | GSR | SDM | Pearson | .21 | 235 | .11 | 74 | .16 | 142 | 121 | .15 | -4.16 | -3.08 | .56 | .88 | .60 | .73 | -.14 | -.12 | -.15 | .04 |
| Lausanne463 + GSR + weig + FD5% + Pearson | Anat | Scale 400 | Weighted | GSR | FD5% | Pearson | .19 | 205 | .09 | 51 | .14 | 104 | 96 | .13 | -4.24 | -3.03 | .56 | .88 | .56 | .78 | -.15 | -.23 | -.16 | .09 |
| Brainnetome246 + GSR + weig + OMST + Pearson | Single | Scale 200 | Weighted | GSR | OMST | Pearson | .10 | 60 | .06 | 13 | .07 | 11 | 25 | .08 | -3.21 | -2.89 | .67 | .72 | .72 | .73 | .10 | .39 | -.23 | -.09 |
| Glasser414 + GSR + weig + FD5% + Pearson | Single | Scale 400 | Weighted | GSR | FD5% | Pearson | .16 | 163 | .10 | 65 | .13 | 82 | 82 | .12 | -3.22 | -2.66 | .56 | .80 | .60 | .82 | -.14 | -.30 | -.04 | .04 |
| Schaefer454 + NoGSR + weig + OMST + Pearson | Func | Scale 400 | Weighted | No GSR | OMST | Pearson | .10 | 66 | .08 | 34 | .10 | 41 | 63 | .12 | -3.01 | -2.63 | .78 | .92 | .72 | .76 | -.35 | -.09 | -.05 | -.03 |
| Schaefer116 + GSR + weig + OMST + Pearson | Func | Scale 100 | Weighted | GSR | OMST | Pearson | .13 | 128 | .11 | 73 | .10 | 50 | 69 | .11 | -2.03 | -2.50 | .56 | .56 | .64 | .82 | -.27 | .29 | -.26 | .02 |
| Brainnetome246 + NoGSR + weig + OMST + Pearson | Single | Scale 200 | Weighted | No GSR | OMST | Pearson | .14 | 144 | .09 | 49 | .10 | 39 | 83 | .13 | -2.25 | -2.13 | .78 | .84 | .80 | .69 | -.20 | .05 | -.04 | -.03 |

Overall, inspecting the whole list of optimal pipelines (Table 1) clearly reveals that considering each pipeline step in isolation from the others does not provide the full picture. Specifically, we found that a few combinations of options account for most of the optimal pipelines (Figure 8), with 5 out of 8 pipelines which meet all inclusion criteria using the combination of weighted edges, Pearson correlation and OMST filtering for edge definition and thresholding.

These results suggest that a pipeline’s performance is not solely attributable to any specific step: rather, some combinations of steps seem to be especially favourable.

**Fig. 8.**
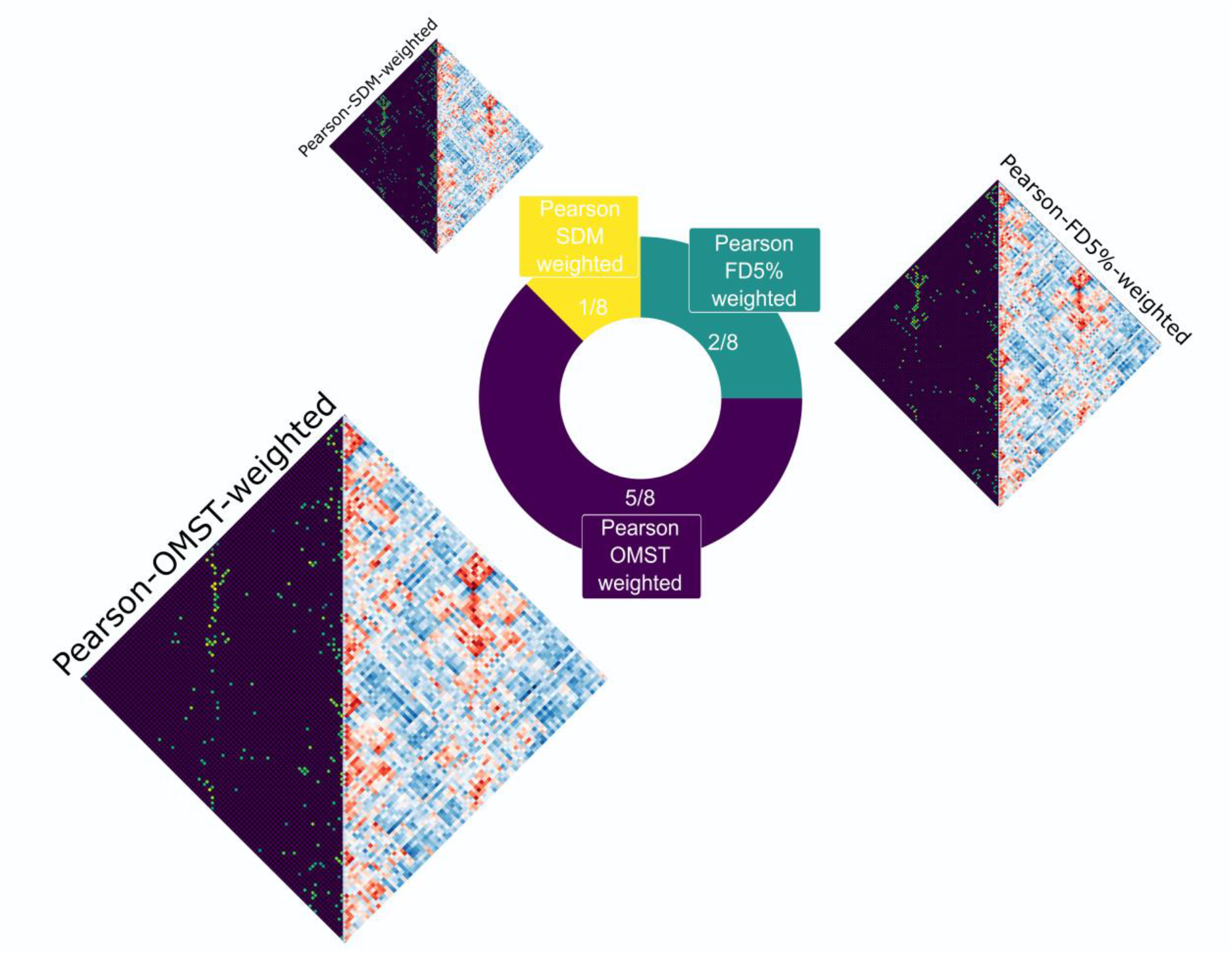
Optimal edge processing combinations. Pie chart displays the frequency of each combination of edge type definition, filtering, and binarisation among the 8 pipelines which fulfil all criteria for a suitable network construction pipeline. See Fig. S32 for a version of this figure with a breakdown of the pipelines under the more liberal global rank criterion, and Fig. S33-35 for a breakdown of the relationship between PDiv and commonly studied graph properties, in terms of edge quantification, binarisation, and filtering method.

## Discussion

A tremendous amount of neuroimaging research with functional MRI is devoted to finding reliable functional connectomic biomarkers for brain function and its disorders – but this process involves a combinatorial explosion of arbitrary choices^18, 19, 28^.

Here, we tackled this challenge by systematically investigating 576 unique pipelines that a neuroscientist could adopt to obtain brain networks from resting-state fMRI data, arising from the combination of several key data-processing steps. To do so, we departed from most previous studies in a number of key respects. First, we explicitly addressed the combinatorial explosion, by considering pipelines end-to-end, rather than restricting our attention to specific steps. Second, rather than choosing any arbitrary local or global graph-theoretical property for our comparisons, we focused on the pipelines’ ability to recover the networks’ overall topology across all scales. Third, we did not focus exclusively on test-retest reliability, but rather we adopted an entire battery of criteria that any appropriate pipeline for functional connectomics should meet, in order to provide practically useful results: these include minimising both random (noise-induced) and systematic (motion-induced) topological distortions, while also being sensitive to differences between individuals and between experimental conditions. Finally, we required all criteria to be consistently met in each of several independent datasets, encompassing short (minutes), medium (weeks) and long timespans (up to 16 months), and using different spatial and temporal resolution, and different preprocessing/denoising approaches, to ensure the generalisability of our recommendations. Through this multi-dataset, multi-criteria, multi-scale and multi-step approach, our goal was to provide a comprehensive set of benchmarks for trustworthy functional connectomics.

### Inappropriate network pipelines are ubiquitous and can produce systematically misleading results

Our first finding is that the substantial majority of the pipelines that we considered failed to meet at least one of our criteria for consistent functional connectomics. We also observed drastic and systematic variability among pipelines’ performance: an inappropriate choice of pipeline can greatly impair one’s ability to recover a reliable network topology. Even for scans obtained less than 45 minutes apart, we observed up to a 5-fold increase in topological dissimilarity (PDiv) compared with the best-performing pipelines (Fig. 2), even across several months. Put differently, adoption of an inappropriate pipeline can distort the functional connectome more drastically than the passage of nearly a year – which may have far-reaching repercussions for longitudinal studies of brain network properties.

A recent review of statistical power in network neuroscience suggested that “many real effects may be missed by current studies”^60^. Our results are in line with this observation: we found that the vast majority (approximately 90%) of pipelines were unable to reliably detect the effect of general anaesthesia on the functional connectome. Thus, one potential implication of our work is that some true effects may have been missed due to a suboptimal choice of network construction pipelines for functional connectomics.

Even more worryingly, choice of the wrong pipeline can lead to results that are not only misleading (statistically significant in the opposite direction as the true effect), but *replicably so* (being observed in two independent propofol datasets): we found this to be the case for 38 pipelines. This means that adopting an inappropriate pipeline for network analysis can turn the replicability of results *against* researchers, boosting their confidence in results that are actively the opposite of the truth. Being consistently wrong rather than randomly so, these results would not be “washed out” by approaches such as meta-analytic aggregation: on the contrary, they would propagate to the meta-analysis itself. Clearly, such a scenario would have devastating consequences for the use of functional connectomics for biomarker identification; in the worst-case scenario, a treatment that actually makes the disease worse may be systematically mis-identified as making it better.

Finally, our results show that the above-mentioned concerns cannot be easily dismissed, because suboptimal pipelines are not a rare exception, but rather the rule: the vast majority of pipelines among those considered (568 out of 576) failed to meet at least one of our criteria (or 550 if the criterion of having low PDiv on average is relaxed). In other words, our results clearly demonstrate that even when combining steps for network construction that are individually sensible, it is overwhelmingly likely (over 98%) that the resulting overall pipeline will *not* be appropriate for functional connectomics – at least not optimal. Indeed, we find that no single step uniquely determines a pipeline’s ability (or inability) to accurately recover the network’s topology: pipelines differing by only one step are largely overlapping in terms of their portrait divergence distributions (Fig. S7-S30). These observations highlight the importance of focusing on entire pipelines as we do here, in contrast to most published approaches that typically consider only one or two steps in isolation.

### Identification of optimal network construction pipelines

Fortunately, we were able to identify a number of pipelines (8 out of 576) that consistently recover effects in the correct direction, and that additionally satisfy all our other criteria for trustworthy functional connectomics: low PDiv for test-retest scans, indicating that the pipeline minimises spurious differences; greater PDiv across subjects than within the same subject on average, indicating that the pipeline reflects the ground-truth difference between networks; no empty networks; and no correlation between PDiv and motion. We emphasise that each of these criteria had to be met in all our datasets, which included both differences in time-span, and also differences in data resolution and preprocessing/denoising.

Additionally, we found that pipelines’ performance on our criteria is far from random, nor does it vary idiosyncratically with each dataset, instead being highly correlated across different independent datasets spanning short, medium, and long timespans (with Spearman’s *rho* ranging between .71 and .98; Fig.3). Ability to minimise test-retest differences is also correlated with a pipeline’s ability to detect true differences, when they do exist – both between different individuals (Fig. 5), and within the same individual (induced by potent pharmacological intervention; Fig. 4). In other words, there are systematic factors at play. Indeed, patterns of similarity clearly emerge among the pipelines that satisfy all our criteria. Specifically, 5 out of 8 optimal pipelines employ the same procedure for edge definition (out of 32 possible ones), consisting of Pearson correlation, weighted edges, and the OMST method of optimising the balance between network efficiency and wiring cost. This is a statistically unlikely occurrence, suggesting that there may be something about this combination that makes it especially appropriate. In fact, all 8 (or 22/26 under the less stringent PDiv criterion) employ Pearson correlation for edge definition. More combinations for edge construction become available if pipelines with PDiv rank in the top 50% are included, with fixed-density thresholds at 5% and 20% density also performing well in combination with weighted and binary edges, respectively (6/26 pipelines each). The edge construction part of the pipeline therefore appears as the most crucial choice: once it is fixed, both GSR and NoGSR options are available among the optimal pipelines, and many combinations of atlas type and size.

It is especially reassuring that our results about pipeline performance are shared across multiple independent datasets. Likewise, our results generalise across different popular methods for functional MRI denoising (aCompCor and FIX-ICA). The Cambridge and NYU datasets were acquired with parameters for spatial and temporal resolution that are widely used in functional neuroimaging studies. Therefore, we expect our results to generalise to other datasets with similar specifications, such as the publicly available and intensely studied Cam-CAN^61^, Philadelphia Neurodevelopmental Cohort^62^, CENTER-TBI^63^, Harvard Aging Brain Study^64^, Autism Brain Imaging Data Exchange (ABIDE)^65^, and UCLA Neurophenomics^66^ datasets, enabling the functional connectomics community to make the most of these valuable resources to study development, aging, and disease. Importantly though, our results about pipeline performance and choice of optimal pipelines also replicated in the high-quality HCP data, which have higher temporal and spatial resolution (suitable for surface-based analysis). Therefore, we expect that our recommendations should also be applicable to more recent datasets acquired with HCP-like specifications, such as UK Biobank^67^. However, our recommendations are intended to complement investigators’ domain-expertise, *not* replace it: each study has its own driving hypotheses and unique challenges. For this reason, we have made available our Supplementary Interactive Tool, which provides a full breakdown of each pipeline’s performance across each criterion and each dataset: to enable readers to engage with our results, and identify pipelines that fit their specific requirements.

Our optimal pipelines are those that pass *all* our tests across *all* datasets: they minimise noise-driven differences, but correctly detect genuine ones, in a way that is consistent across datasets. While this stringency undoubtedly contributed to the exclusion of many pipelines – 84 of which only due to a single failure in a single dataset – it should equally bolster our confidence about the recommended pipelines’ suitability to provide sensible results, including across different time-spans and different data acquisition and preprocessing choices. By recommending a select number of network construction pipelines that provide the most replicable and generalisable results, we hope that the present work will facilitate future meta-analyses of functional connectomics studies.

### Shared characteristics among optimal network construction pipelines

It is reassuring that our recommended pipelines overwhelmingly favour Pearson correlation to quantify functional connectivity. Owing to its ease of application and interpretation, Pearson correlation is a cornerstone of functional connectomics, and remains the most widely used method to quantify connectivity between regions across thousands of published studies (accounting for over 75% of the studies reviewed by Hallquist and Hillary). Nevertheless, here we did not assume a priori that fMRI BOLD signals are linear, and instead also considered a nonlinear method (MI).

At the microscopic level of neurons and circuits, the brain is unquestionably a nonlinear system. However, the superior performance of (linear) Pearson correlation that we observed in our results dovetails with multiple lines of evidence that the macroscale level observed by functional MRI signals may be suitably accounted for as linear ^68, 69^, such that limited or no additional benefit is obtained when using more complex nonlinear methods to relate structural and functional connectivity ^70, 71^, or to predict demographic variables from functional MRI ^72^, or when comparing the ability of linear versus nonlinear models to fit high-resolution BOLD timeseries ^73^. Crucially, the observed predominance of linear dynamics in macroscale brain signals cannot be dismissed as a mere artifact of functional MRI ^73^. Although fMRI’s low temporal resolution does contribute to linearising the signal due to both temporal averaging and the limited number of samples, linear models were also recently shown to outperform nonlinear ones in terms of their ability to fit intracranial EEG (iEEG) time-series ^73^, which are electrodynamic rather than haemodynamic in origin, and have much higher temporal resolution. Thus, empirical results from diverse neuroimaging modalities converge with both simulations ^73^ and theoretical analysis ^74^, showing that the dynamics of nonlinear stochastic populations converge to linear dynamics at the macroscale, as a result of spatial averaging. In other words, observing superior performance of linear methods at the macroscale should not be viewed as un-physiological, or a mere artifact of a specific imaging modality, or a denial of the brain’s microscale nonlinearity. Rather, linearisation is an inherent consequence of observing brain activity at the macroscale, and this phenomenon contributes to explaining why Pearson correlation is suitable for quantifying functional connectivity.

Pertaining to edge filtering, the OMST – our main recommended approach – is a data-driven method that optimises the balance between efficiency and wiring cost of the network. OMST is unique among the filtering schemes considered here, for multiple reasons. First, because it guarantees that the resulting network is not fragmented into disconnected components (Fig. S35). This feature makes OMST analogous to percolation-based filtering schemes, whereby the weakest edges are iteratively removed from the network, up to the point where further removal would make the network disconnected, which corresponds to the percolation threshold^75–77^. Thus, OMST and percolation thresholding both ensure that global connectivity is not impacted by removal of a few weak but topologically important edges. Unlike percolation, however, OMST is not restricted to preserving only the strongest edges. Rather, weaker edges can be preferred to stronger ones and be included in the OMST-filtered network, if they contribute to an optimal balance of efficiency and cost. Because of this ability to include weaker edges over stronger ones based on their role in the overall topology, OMST avoids a pitfall of percolation thresholding, whereby the presence of a single node whose edges are all relatively weak, can result in a network that is potentially very dense (because the percolation threshold is determined by the weakest edge whose removal would make the network disconnected, and if this edge is very weak, many other edges may survive the threshold).

In other words, the second feature that makes OMST unique among the filtering schemes considered here is that OMST takes into account not only the strength of connections, but also their more general topological role in the network. Therefore, connectomes obtained through OMST can include edges that both absolute and proportional thresholding methods would simply disregard as too weak, regardless of any further role they may play in network organisation. The key role of weak connections acting as shortcuts between segregated modules, often referred to as the “strength of weak ties”^77–79^, has been increasingly recognised across artificial and biological networks, including the human brain – a clear argument in favour of OMST’s ability to reconstruct biologically plausible networks, especially in combination with weighted (rather than binary) edges, which is consistent with our optimal pipelines.

It is key to note that despite the similar name, OMST is very different in practice from simple Minimum Spanning Tree filtering. Reducing the network to its minimum spanning tree will enforce every individual’s network to have the same number of edges, which is the minimum number possible. In contrast, while the OMST does ensure the desirable property of network connectedness, it determines the final number of edges in a data-driven manner by optimising the network’s balance of efficiency and wiring cost. This approach therefore produces plausibly sparse networks, but without imposing the same a-priori level across all individuals (arguably a biologically implausible feature of fixed-density methods).

The good performance of OMST is arguably due to this method being data-driven based on each individual connectome, rather than a one-size-fits-all. Indeed, although OMST is a relatively recent method, its use has already been recommended by several studies on multiple grounds. OMST filtering was shown to minimise topological differences between pipelines^54^; it has outperformed alternative thresholding schemes for functional networks in terms of recognition accuracy and reliability^30, 55, 80^; and it has also been recommended for use with alternative neuroimaging modalities such as electro- and magneto-encephalography^30, 55, 80^, suggesting that its applicability may generalise beyond rs-fMRI. Finally, the use of OMST (as well as 20% fixed-density thresholding) was also recommended by another recent study^43^ that evaluated a large number of individual options (though without combining them, and using as criterion the ICC of specific network properties instead of our topological approach). Therefore, our results suggest a convergence of recommendations for brain network construction across different criteria and different studies – possibly heralding the emergence of consistent analytic practices in the field. This convergence may in part be helped by our choice to use the Portrait divergence, which enabled us to take into account both local and global aspects of network organisation across scales^46^: by considering the network’s topology as a whole, our results are inherently more general than results based on any specific graph-theoretical metric.

### Limitations and future directions

In this study, we endeavoured to systematically sample and combine many of the most common options across each step in the process of constructing a functional brain network from rs-fMRI data – resulting in 576 unique pipelines. However, due to combinatorial explosion, it would be unfeasible to consider every single option that has been proposed in the literature, and this inevitable limitation should be borne in mind when interpreting our results. In particular, although we considered some of the most widely used parcellation schemes for defining nodes in the brain, encompassing the most common range of network sizes used in the field, we inevitably could not include all the possible atlases in existence^21, 25, 26, 81, 82^, and we chose to focus on some of the most widely adopted. More broadly, thanks to the interpretability of dividing the brain into discrete, spatially circumscribed regions, atlas-based methods have enjoyed enduring popularity for defining nodes in brain networks ^21, 25^, which motivated the focus of the present work. However, they also come with implicit assumptions about spatial localisation (e.g., by imposing the constraints that parcels should be spatially contiguous and non-overlapping) and about what should be regarded as the functional units of the brain^26^. Indeed, recent gradient-based approaches provide alternative representations of the brain that are spatially extended and continuous rather than discrete, offering a complementary perspective on the constituent elements of the brain’s functional organisation ^27, 83–86^.

Here, we made the pragmatic choice to consider well-established and widely used methods for node definition that vary along some of the most relevant dimensions for network construction. Combinatorial explosion prevented us from extending our investigation to alternative, parcellation-free methods for node definition, such as voxelwise/vertexwise networks with thousands of nodes^37, 87^ (which sacrifice biological interpretability for maximal spatial resolution), and methods based on PCA or (spatial or temporal) ICA that can provide non-contiguous, spatially overlapping parcels, possibly better able to reflect the complexity of brain organisation^88–90^. Simulations previously suggested that defining nodes based on ICA may outperform the use of regions-of-interest (e.g., based on atlases)^90^: future work in this direction may reveal whether some of these alternative approaches to node definition perform consistently better – or consistently worse – across our criteria, than the atlas-based node definitions adopted in the present work. However, based on the pattern of shared features among our recommended pipelines, we note that our results point towards a more prominent role of edge definition than node definition, for determining the success of a pipeline.

Pertaining to edge definition, many alternative thresholding methods also exist, whether based on statistical significance^91^, percolation^75–77^, or shrinkage methods^89, 92^ – or avoiding thresholding entirely, by using analytic methods that can deal with fully connected and signed networks^45^. More broadly, future work may adopt more advanced methods of quantifying connectivity: for instance, by adopting multivariate connectivity estimators^93^ or methods from information decomposition capable of recovering different kinds of information sharing between regions^94–96^ or the directionality of connections (transfer entropy, Granger causality, Dynamic Causal Modelling^97^), or disambiguating between direct and indirect connections (e.g. partial correlation^89^). Additionally, it remains to be determined how our results will generalise to the case of time-varying (“dynamic”) networks, an increasingly popular approach in fMRI functional connectivity^98–100^, and to frequency-specific networks obtained from EEG or MEG^101^ (although see Jiang et al. ^43^ and Dimitriadis et al. ^102, 103^, for recent investigations of frequency bands for fMRI network construction).

It is also known that different motion correction strategies can influence the validity of BOLD signals and subsequent network characteristics; however, no correction strategy offered perfect motion correction^23^. Here, we adopted a widely used denoising strategy (anatomical CompCor), and required our results to also replicate in a dataset denoised with FIX-ICA instead, which unlike aCompCor is designed to affect artifacts specifically and avoid modifying the neural signal of interest^51, 52^. Additionally, we also considered two versions of each dataset, preprocessed with versus without the additional step of global signal regression, due to ongoing controversy about the effect of GSR on functional connectivity^53, 104^. Finally, to further mitigate the potential impact of motion on our recommendations, we also explicitly included as one of our criteria that pipelines should not produce a PDiv distribution that is significantly correlated with the distribution of differences in subject motion, across any of the four test-retest datasets. We note that when GSR is included, PDiv tends to be smaller across all datasets – possibly reflecting the elimination of residual noise. However, our final recommendations include pipelines both with and without GSR – although the latter is somewhat more prevalent among the very best-performing ones. In particular, we even found that the set of optimal pipelines includes versions of the same pipeline both with and without GSR: Brainnetome-246 for Pearson-OMST-weighted (with GSR and no-GSR versions both featuring among the 8 optimal pipelines); and in the expanded set, Schaefer454 Top20%-binary-Pearson, Lausanne-463 Top20%-binary-

Pearson, Lausanne-463 Top20%-binary-MutualInfo. Therefore, our results suggest that investigators may have some discretion in the choice of using GSR, depending on their specific datasets and hypotheses. As an example, GSR may remove physiological and motion-induced noise^48, 50^, and it may strengthen brain-behaviour associations^104^, but it can also remove signal of interest pertaining to some pharmacological and pathological conditions^32, 105^, or distort group^33^ and individual differences^106^. Likewise, a recent study observed reduced generalisability of graph-theoretical properties across sites, sessions, and paradigms when GSR was used^36^, although Tozzi et al (2020)^42^ delineated a more intricate picture, whereby GSR decreases reliability for networks and most edges, but increases it for some others. A comprehensive evaluation of the relative advantages and drawbacks of GSR is beyond the scope of this paper, and the reader is referred to Fox & Murphy (2017)^53^ and Liu et al. (2017)^104^ for extensive discussions. Finally, we did not explore potential differences between resting-state conditions (eyes-open vs eyes-closed vs naturalistic viewing)^40, 107^, or the impact of scan duration and spontaneous fluctuations in arousal state – although we did include datasets with different scan duration, up to 1200 volumes ^18, 108^.

In addition to increasing the number of options and pipelines considered, future work may further expand on the present results in several ways: it remains to be determined to what extent our results apply to task-based rather than resting-state fMRI ^109, 110^. The generalizability of the proposed framework beyond healthy individuals is also worthy of future exploration. Compared to healthy controls, some clinical populations have demonstrated lower test-retest reliability^111, 112^. Reliability across the lifespan should be also considered by comparing age-groups, as early evidence untangled age-related differences in test-retest reliability of rs-fMRI^113^. The choice of the optimal pipeline for functional connectomics may therefore vary by clinical characteristics, which still remains to be ascertained and may benefit from topology-based approaches such as the one adopted here. This is an important next step following the present work. It is also possible that a different proportion of optimal pipelines would be found when alternative reconstruction methods are included, or different neuroimaging modalities.

The time-spans that we considered here range from less than an hour to nearly a year between scans. Certainly, in addition to measurement noise, some degree of change in the topology of the functional connectome over the course of weeks or months is to be expected, due to learning and plasticity. However, such physiological phenomena cannot be expected to appreciably reorganise the entire functional connectome within the span of less than an hour (in the absence of experimental interventions). Therefore, any test-retest PDiv observed within the same hour is most plausibly attributable to noise, and an appropriate pipeline should simply minimise it, as per our test-retest criterion. Additionally, plasticity and learning should not make the functional connectome so different that it becomes indistinguishable from the connectomes of other individuals: rather, such an occurrence should be minimised, as it indicates measurement noise. Our results clearly show a convergence of these criteria: pipelines that produce small test-retest PDiv over weeks or months are also those that minimise within-hour PDiv, and that minimise the mis-identification of individuals. Thus, across datasets and time-spans we observed an encouraging convergence of criteria for reliable functional connectomics.

As a final note, our approach has been to identify which pipelines produce sensible functional connectomes, so that researchers may have a guide to orient their choice among the “forking paths” of analytical possibilities. However, an alternative approach exists: performing a “multiverse” analysis, adopting not one but many pipelines and then finding suitable ways to aggregate the results – or using machine learning tools to characterise a low-dimensional space of pipelines^114^. The two approaches are not mutually exclusive, but rather complementary: our criteria and our final recommendations could be used to prune the number of branching options to a manageable number of optimal pipelines, and a multiverse analysis could then be carried out in parallel across them, with the confidence that the overall picture will not be contaminated by inappropriate choices.

## Conclusion

In conclusion, our study provides a principled framework to search for the best network construction pipelines across hundreds of candidates, with the aim of recovering brain networks that satisfy multiple criteria for scientific accuracy and practical utility. We revealed drastic differences across pipelines in terms of their ability to recover similar network topologies across different scans of the same individual – even within the same hour – and to recover the true directionality of experimental effects of interest. The existence and prevalence of systematically misleading pipelines further enhances the importance of identifying suitable network construction pipelines. Thus, our results indicate that researchers should pay careful consideration to their choice of network processing pipeline: pipelines vary widely in their ability to detect true effects while mitigating spurious ones, and the vast majority of pipelines are not optimal. Our findings further indicate that no single step in the network construction workflow can single-handedly guarantee that all criteria will be met. Fortunately, however, we also show that by carefully combining different steps in the network construction workflow, neuroscientists can obtain functional brain networks that satisfy all our criteria, across datasets covering different time-spans and different acquisition and preprocessing procedures, and may be used with confidence. These recommendations can inform future studies, to help investigators make principled choices and minimise the chance that an inappropriate choice of network construction will lead to unreliable or false negatives results. Overall, by enabling systematic evaluation of network processing steps in a way that does not require the arbitrary selection of specific network properties of interest, we hope that the topology-based, multi-criteria framework proposed here will lead towards an objective consensus and more consistent practices in functional connectomics.

## Materials and Methods

### NYU Test-Retest dataset

This is an open dataset from the International Neuroimaging Data-Sharing Initiative (INDI) (http://www.nitrc.org/projects/nyu_trt), originally described in Shehzad et al., (2009)^115^. Briefly, this dataset includes 25 participants (mean age 30.7 ± 8.8 years, 16 females) with no history of psychiatric or neurological illness. The study was approved by the institutional review boards of the New York University School of Medicine and New York University, and participants provided written informed consent and were compensated for their participation.

For each participant, 3 resting-state scans were acquired. Scans 2 and 3 were conducted in a single scan session, 45 min apart, which took place on average 11 months (range 5–16 months) after scan 1. Each scan was acquired using a 3T Siemens (Allegra) scanner, and consisted of 197 contiguous EPI functional volumes (TR = 2000 ms; TE = 25 ms; flip angle = 90°; 39 axial slices; field of view (FOV) = 192 × 192 mm2; matrix = 64 × 64; acquisition voxel size = 3 × 3 × 3 mm3). Participants were instructed to relax and remain still with their eyes open during the scan. For spatial normalization and localization, a high-resolution T1-weighted magnetization prepared gradient echo sequence was also obtained (MPRAGE, TR = 2500 ms; TE = 4.35 ms; TI = 900 ms; flip angle = 8°; 176 slices, FOV = 256 mm).

### Cambridge test-retest dataset

Right-handed healthy participants (N=22, age range, 19–57 years; mean age, 35.0 years; SD 11.2; female-to-male ratio, 9/13) were recruited via advertisements in the Cambridge area and were paid for their participation. Cambridgeshire 2 Research Ethics Committee approved the study (LREC 08/H0308/246) and all volunteers gave written informed consent before participating. Exclusion criteria included National Adult Reading Test (NART) <70, Mini Mental State Examination (MMSE) <23, left-handedness, history of drug/alcohol abuse, history of psychiatric or neurological disorders, contraindications for MRI scanning, medication that may affect cognitive performance or prescribed for depression, and any physical handicap that could prevent the completion of testing.

The study consisted of two visits (separated by 2–4 weeks). For each visit, resting-state fMRI was acquired for 5:20 minutes using a Siemens Trio 3T scanner (Erlangen, Germany). Functional imaging data were acquired using an echo-planar imaging (EPI) sequence with parameters TR 2,000 ms, TE 30 ms, Flip Angle 78◦, FOV 192 × 192mm2, in-plane resolution 3.0 × 3.0mm, 32 slices 3.0mm thick with a gap of 0.75mm between slices. A 3D high resolution MPRAGE structural image was also acquired, with the following parameters: TR 2,300 ms, TE 2.98 ms, Flip Angle 9◦, FOV 256 × 256 mm2. Task-based data were also collected, and have been analysed before to investigate separate experimental questions^116, 117^ . A final set of 18 participants had usable data for both resting-state fMRI scans and were included in the present analysis.

### Human Connectome Project test-retest data

This dataset is a subset of the 1,200 Human Connectome Project (HCP) subjects^57, 58^. It includes resting-state functional MRI (and accompanying structural MRI) scans for 46 healthy individuals (13 male, age 22–35 years), who were each scanned twice at 3T, at intervals ranging between 1 month and 11 months). All HCP scanning protocols were approved by the local Institutional Review Board at Washington University in St. Louis. Detailed information about the acquisition and imaging is provided in the dedicated HCP publications. Briefly: anatomical (T1-weighted) images were acquired in axial orientation, with FOV = 224 × 224 mm, voxel size 0.7 mm3 (isotropic), TR 2,400ms, TE 2.14ms, flip angle 8°. Functional MRI data (1200 volumes) were acquired with EPI sequence, 2 mm isotropic voxel size, TR 720ms, TE 33.1ms, flip angle 52°, 72 slices.

### Cambridge propofol dataset

The Cambridge University (“Cambridge”) propofol dataset has been published before^118–120;^ we refer the reader to the original study for a detailed description^118^. As previously reported, 16 healthy volunteer subjects were initially recruited for scanning. In addition to the original 16 volunteers, data were acquired for nine participants using the same procedures, bringing the total number of participants in this dataset to 25 (11 males, 14 females; mean age 34.7 years, SD = 9.0 years). Ethical approval for these studies was obtained from the Cambridgeshire 2 Regional Ethics Committee, and all subjects gave informed consent to participate in the study. Volunteers were informed of the risks of propofol administration, such as loss of consciousness, respiratory and cardiovascular depression. They were also informed about more minor effects of propofol such as pain on injection, sedation and amnesia. In addition, standard information about intravenous cannulation, blood sampling and MRI scanning was provided.

Three target plasma levels of propofol were used: no drug (Awake), 0.6 mg/ml (Mild sedation) and 1.2 mg/ml (Moderate sedation). Scanning (rs-fMRI) was acquired at each stage, and also at Recovery; anatomical images were also acquired. The level of sedation was assessed verbally immediately before and after each of the scanning runs. Propofol was administered intravenously as a “target controlled infusion” (plasma concentration mode), using an Alaris PK infusion pump (Carefusion, Basingstoke, UK). A period of 10 min was allowed for equilibration of plasma and effect-site propofol concentrations. Blood samples were drawn towards the end of each titration period and before the plasma target was altered, to assess plasma propofol levels. In total, 6 blood samples were drawn during the study. The mean (SD) measured plasma propofol concentration was 304.8 (141.1) ng/ml during mild sedation, 723.3 (320.5) ng/ml during moderate sedation and 275.8 (75.42) ng/ml during recovery. Mean (SD) total mass of propofol administered was 210.15 (33.17) mg, equivalent to 3.0 (0.47) mg/kg. Two senior anaesthetists were present during scanning sessions and observed the subjects throughout the study from the MRI control room and on a video link that showed the subject in the scanner. Electrocardiography and pulse oximetry were performed continuously, and measurements of heart rate, non-invasive blood pressure, and oxygen saturation were recorded at regular intervals.

The acquisition procedures are described in detail in the original study^118^. As previously reported, MRI data were acquired on a Siemens Trio 3T scanner (WBIC, Cambridge). For each level of sedation, 150 rs-fMRI volumes (5 min scanning) were acquired. Each functional BOLD volume consisted of 32 interleaved, descending, oblique axial slices, 3 mm thick with interslice gap of 0.75 mm and in-plane resolution of 3 mm, field of view = 192×192 mm, TR = 2000 ms, acquisition time = 2000 ms, time echo = 30 ms, and flip angle 78. T1-weighted structural images at 1 mm isotropic resolution were also acquired in the sagittal plane, using an MPRAGE sequence with TR = 2250 ms, TI = 900 ms, TE = 2.99 ms and flip angle = 9 degrees, for localization purposes. During scanning, volunteers were instructed to close their eyes and think about nothing in particular throughout the acquisition of the resting state BOLD data. Of the 25 healthy subjects, 15 were ultimately retained (7 males, 8 females): 10 were excluded, either because of missing scans (n=2), or due of excessive motion in the scanner (n=8, 5mm maximum motion threshold). Here, we only use data from the Awake and Moderate anaesthesia resting-state scanning.

### Western propofol dataset

The Western University (“Western”) propofol data have been published before^14, 120, 121^ and we refer the reader to the original study for a detailed description. Briefly, data were collected between May and November 2014 at the Robarts Research Institute, Western University, London, Ontario (Canada). The study received ethical approval from the Health Sciences Research Ethics Board and Psychology Research Ethics Board of Western University (Ontario, Canada). Healthy volunteers (n=19) were recruited (18–40 years; 13 males). Volunteers were right-handed, native English speakers, and had no history of neurological disorders. In accordance with relevant ethical guidelines, each volunteer provided written informed consent, and received monetary compensation for their time. Due to equipment malfunction or physiological impediments to anaesthesia in the scanner, data from n=3 participants (1 male) were excluded from analyses, leaving a total n=16 for analysis^14^.

Resting-state fMRI data were acquired at different propofol levels: no sedation (Awake), Deep anaesthesia (corresponding to Ramsay score of 5) and also during post-anaesthetic recovery. As previously reported^14^, for each condition fMRI acquisition began after two anaesthesiologists and one anaesthesia nurse independently assessed Ramsay level in the scanning room. The anaesthesiologists and the anaesthesia nurse could not be blinded to experimental condition, since part of their role involved determining the participants’ level of anaesthesia. Note that the Ramsay score is designed for critical care patients, and therefore participants did not receive a score during the Awake condition before propofol administration: rather, they were required to be fully awake, alert and communicating appropriately. To provide a further, independent evaluation of participants’ level of responsiveness, they were asked to perform two tasks: a test of verbal memory recall, and a computer-based auditory target-detection task. Wakefulness was also monitored using an infrared camera placed inside the scanner.

Propofol was administered intravenously using an AS50 auto syringe infusion pump (Baxter Healthcare, Singapore); an effect-site/plasma steering algorithm combined with the computer-controlled infusion pump was used to achieve step-wise sedation increments, followed by manual adjustments as required to reach the desired target concentrations of propofol according to the TIVA Trainer (European Society for Intravenous Aneaesthesia, eurosiva.eu) pharmacokinetic simulation program. This software also specified the blood concentrations of propofol, following the Marsh 3-compartment model, which were used as targets for the pharmacokinetic model providing target-controlled infusion. After an initial propofol target effect-site concentration of 0.6 *µ*g mL^-1^, concentration was gradually increased by increments of 0.3 *µ*g mL^1^, and Ramsay score was assessed after each increment: a further increment occurred if the Ramsay score was lower than 5. The mean estimated effect-site and plasma propofol concentrations were kept stable by the pharmacokinetic model delivered via the TIVA Trainer infusion pump. Ramsay level 5 was achieved when participants stopped responding to verbal commands, were unable to engage in conversation, and were rousable only to physical stimulation. Once both anaesthesiologists and the anaesthesia nurse all agreed that Ramsay sedation level 5 had been reached, and participants stopped responding to both tasks, data acquisition was initiated. The mean estimated effect-site propofol concentration was 2.48 (1.82-3.14) *µ*g mL^-1^, and the mean estimated plasma propofol concentration was 2.68 (1.92-3.44) *µ*g mL^-1^. Mean total mass of propofol administered was 486.58 (373.30-599.86) mg. These values of variability are typical for the pharmacokinetics and pharmacodynamics of propofol. Oxygen was titrated to maintain SpO2 above 96%.

At Ramsay 5 level, participants remained capable of spontaneous cardiovascular function and ventilation. However, the sedation procedure did not take place in a hospital setting; therefore, intubation during scanning could not be used to ensure airway security during scanning. Consequently, although two anaesthesiologists closely monitored each participant, scanner time was minimised to ensure return to normal breathing following deep sedation. No state changes or movement were noted during the deep sedation scanning for any of the participants included in the study^14^. Propofol was discontinued following the deep anaesthesia scan, and participants reached level 2 of the Ramsey scale approximately eleven minutes afterwards, as indicated by clear and rapid responses to verbal commands.

As previously reported^14^, once in the scanner participants were instructed to relax with closed eyes, without falling asleep. Resting-state functional MRI in the absence of any tasks was acquired for 8 minutes for each participant. A further scan was also acquired during auditory presentation of a plot-driven story through headphones (5-minute long). Participants were instructed to listen while keeping their eyes closed. The present analysis focuses on the resting-state data only, from the Awake and Deep scanning; the story scan data have been published separately^122^ and will not be discussed further here.

As previously reported^14^, MRI scanning was performed using a 3-Tesla Siemens Tim Trio scanner (32-channel coil), and 256 functional volumes (echo-planar images, EPI) were collected from each participant, with the following parameters: slices = 33, with 25% inter-slice gap; resolution = 3mm isotropic; TR = 2000ms; TE = 30ms; flip angle = 75 degrees; matrix size = 64x64. The order of acquisition was interleaved, bottom-up. Anatomical scanning was also performed, acquiring a high-resolution T1-weighted volume (32-channel coil, 1mm isotropic voxel size) with a 3D MPRAGE sequence, using the following parameters: TA = 5min, TE = 4.25ms, 240x256 matrix size, 9 degrees flip angle^14^.

### Functional MRI preprocessing and denoising

Preprocessing of the functional MRI data for all datasets except HCP followed the same standard workflow as in our previous studies^54^, and was implemented in the CONN toolbox (http://www.nitrc.org/projects/conn), version 17f^123^. The following steps were performed: removal of the first 5 volumes to allow for steady-state magnetisation; functional realignment, motion correction, and spatial normalisation to Montreal Neurological Institute (MNI-152) standard space with 2×2×2mm isotropic resolution. Denoising followed the anatomical CompCor (aCompCor) method of removing cardiac and motion artifacts, by regressing out of each individual’s functional data the first 5 principal components corresponding to white matter signal, and the first 5 components corresponding to cerebrospinal fluid signal, as well as six subject-specific realignment parameters (three translations and three rotations) and their first-order temporal derivatives, and nuisance regressors identified by the artifact detection software *art*^124^. The subject-specific denoised BOLD signal time-series were linearly detrended and band-pass filtered between 0.008 and 0.09 Hz to eliminate both low-frequency drift effects and high-frequency noise. No spatial smoothing was applied, since all analyses were performed on parcellated data, whereby the signal was averaged across voxels belonging to the same ROI (see below, section *Node definition*).

For the HCP test-retest dataset, we instead used the minimally preprocessed functional data made available by HCP, which were further denoised with FIX-ICA^51, 52^. This popular approach is intended to remove non-BOLD noise arising from multiple known sources, including spatially specific noise from head motion, cardiac pulsation, breathing, and scanner artifacts. Using different denoising methods enables us to ensure that our final results are not specific to a particular way of denoising rs-fMRI data, thereby ensuring their robustness and generalisability.

A further, particularly controversial denoising step is global signal regression (GSR): although some authors suggest that GSR may improve subsequent construction of functional brain networks^35, 41^, others did not find such an effect^34, 37^ or even reported GSR as deleterious^36, 42^. Here, we therefore evaluated the performance of different network construction pipelines on two versions of each dataset: with the application of GSR, and without the application of GSR.

### Node definition

When deciding on how to turn preprocessed and denoised fMRI data into a brain network, the first decision that needs to be made is: what are the elements of the network? Different approaches exist in the literature, from the use of each voxel as a node to maximise spatial resolution, to the use of Independent Components Analysis and similar data-driven techniques to obtain study- or even subject-specific clusterings of brain signals, which may be spatially extended or even nested within each other, coalescing and splitting over time. Although each of these approaches has unquestionable merits, perhaps the most common approach for defining nodes in human network neuroscience is the use of parcellations: pre-defined assignments of spatially contiguous voxels into regions-of-interest (ROIs) – typically on the ground of neuroanatomical/cytoarchitectonic considerations, or shared function, or some combination thereof. A wide variety of parcellations exist^21^, and recent work reported how the choice of parcellation scheme can affect aspects such as structure-function similarity estimation^125^ but also the intra-subject and inter-subject variability of the functional connectome and whole-brain resting-state modeling^42, 126^. Parcellation schemes vary on two main dimensions: the criterion based on which clusters are identified (e.g., based on neuroanatomy, or functional considerations, or a combination thereof from multiple modalities) and the number of ROIs – ranging from a few tens to thousands. The number of ROIs involves a trade-off between the superior spatial resolution of finer-grained parcellations, and the robustness and increased signal-to-noise ratio that derive from spatial averaging of many neighboring voxels.

Here, we considered both of these dimensions: we employed parcellations spanning three scales (approximately 100, 200 and 400 nodes) and obtained based on anatomical, functional, or multimodal considerations, across one or multiple scales (summarised in Table 2). We consider the multi-scale anatomical Lausanne atlas with 129, 234 and 463 cortical and subcortical nodes obtained by subdividing the sulcus-based Desikan-Killiany atlas^127^. We also consider the functional multi-scale parcellation developed by Schaefer and colleagues^128^ which combines local gradients and global similarity across task-based and resting-state functional connectivity. Following our previous work, we included versions with 100, 200 and 400 cortical regions, respectively supplemented with 16, 32 or 54 subcortical regions from the recent subcortical functional atlas developed by Tian and colleagues^129^.

Finally, we include three widely used single-scale parcellations: (i) the Automated Anatomical Labelling (AAL) atlas, an anatomical parcellation with 90 cortical and subcortical regions^130^; (ii) the Brainnetome atlas, which comprises 210 cortical and 36 subcortical regions, identified by combining anatomical, functional and meta-analytic information^131^; (iii) and the Glasser parcellation comprising 360 cortical regions identified by combining multi-modal information about cortical architecture, function, connectivity, and topography^132^. The volumetric Glasser parcellation in MNI-152 space made available by Preti and Van de Ville^133^ was used. Since the Glasser parcellation is cortical-only, it was also supplemented with the 54-region version of the Melbourne atlas, in order to include a comparable number of subcortical regions, resulting in 414 ROIs.

For all but the HCP dataset, we used parcellations in volumetric MNI-152 space; for each parcellation, the average denoised BOLD timeseries across all voxels belonging to a given ROI were extracted. For the HCP test-retest dataset, given the higher spatial resolution, we opted to use a surface-based parcellation approach instead – thereby enabling us to verify that our final results are not specific to a given parcellation approach.

**Table 2.**
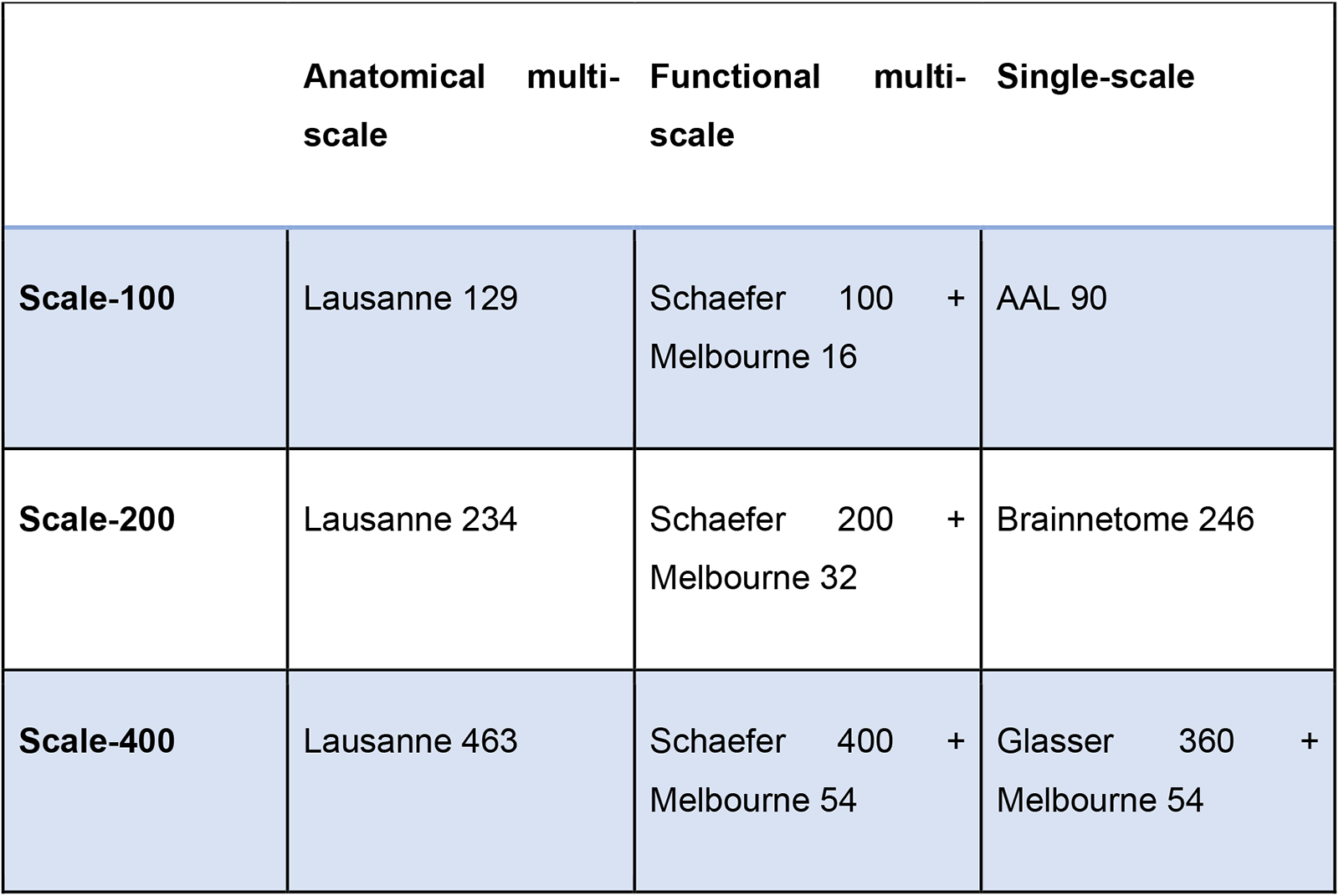
Atlases adopted in the present study, by scale (rows) and method (columns).

### Functional connectivity

We considered two alternative ways of quantifying the interactions between regional BOLD signal timeseries. First, we used Pearson correlation, whereby for each pair of nodes *i* and *j,* their functional connectivity *F_ij_* was given by the Pearson correlation coefficient between the timecourses of *i* and *j*, over the full scanning length. Second, we also used the mutual information *I,* which quantifies the interdependence between two random variables X and Y, and is defined as the average reduction in uncertainty about X when Y is given (or vice versa, since this quantity is symmetric):

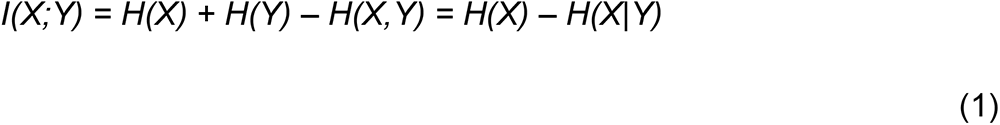

With *H(X)* being the Shannon entropy of a variable X. Unlike Pearson correlation, mutual information considers both linear and nonlinear relationships, and it does not provide negative values. For consistency with previous work^54^, the values in each individual matrix of mutual information were divided by the maximum value in the matrix, thereby rescaling them to lie between zero and unity.

### Filtering Schemes

Both Pearson correlation and MI provide continuous values for the statistical association between pairs of nodes, resulting in a dense matrix of functional connections. Therefore, some form of filtering is typically employed to remove spurious connections that are likely to be driven by noise, and obtain a sparse network of functional connectivity. However, there is no gold standard approach to decide which connections to retain, and different filtering schemes have emerged in the literature. Here, we considered 8 different edge filtering schemes (Table 3), described below. The Brain Connectivity Toolbox^44, 45^ was used to implement absolute and proportional thresholds and quantify network density, as well as the networks’ mean clustering coefficient and characteristic path length (Fig. S33-S35).

**Table 3.** Edge filtering schemes adopted in the present study.

| Filtering Scheme | Description |
| --- | --- |
| Fixed Density 5% (FD5%) | Top 5% of strongest edges |
| Fixed Density 10% (FD10%) | Top 10% of strongest edges |
| Fixed Density 20% (FD20%) | Top 20% of strongest edges |
| Absolute Threshold 0.3<br>(Abs0.3) | Edges with value > 0.3 |
| Absolute Threshold 0.5<br>(Abs0.5) | Edges with value > 0.5 |
| Efficiency Cost Optimisation<br>(ECO) | Average node degree = 3, to maximise trade-off between overall efficiency and wiring cost |
| Structural Density Matching<br>(SDM) | Proportional thresholding, with same density as the HCP group-average DTI data parcellated using the same atlas |
| Orthogonal Minimum Spanning Trees (OMST) | Optimisation of global efficiency minus wiring cost, by combining independent minimum spanning trees of the network. |

### Absolute thresholding

The simplest approach to decide which edges to retain is to accept or reject edges based on a pre-determined minimum acceptable weight. However, there is no consensus in the literature about which threshold one should adopt. Here, we considered absolute threshold values of 0.3 or 0.5 (for Pearson correlation, only positively-value edges were considered).

### Proportional thresholding

Absolute thresholding can produce networks with very different densities, which can introduce confounds in subsequent network analyses. Therefore, a popular approach simply retains a fixed proportion of the strongest edges. However, there is once again no consensus in the literature on the correct proportion of edges to retain. We therefore employed three different density levels, in the range commonly reported in the literature: fixed density (FD) of 5%, 10%, and 20% of the strongest edges.

### Structural Density Matching

The main problem with proportional thresholding is the selection of an appropriate target density – especially since this may vary depending on the number of nodes in the network. To address this issue in a principled manner, we recently introduced a method termed Structural Density Matching (SDM)^54^, whereby the proportion of functional edges to retain corresponds to the density *s* of the corresponding structural connectome (the network of anatomical connectivity obtained from the group-averaged diffusion-weighted MRI data from the Human Connectome Project^134^. In other words, SDM ensures that functional and structural networks obtained using the same parcellation have the same density, instead of using an arbitrary target density.

### Efficiency Cost Optimisation

The Efficiency Cost Optimisation (ECO) is designed to optimise the trade-off between the network’s overall efficiency (sum of global and average local efficiency) and its wiring cost (number of edges)^56^, by ensuring that the network maximises the following target function *J*:

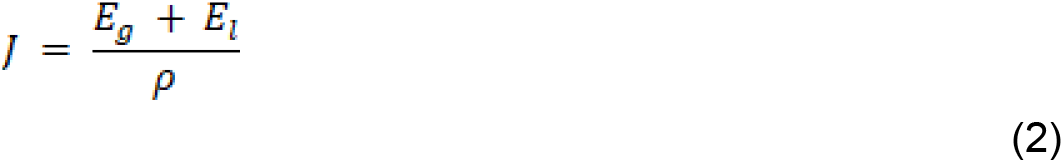

With *E_g_* and *E_i_* being the global and mean local efficiency of the network, respectively. This filtering scheme produces sparse graphs while still preserving their structure, as demonstrated by its empirical success at discriminating between different graph topologies^56^. Here, we obtained ECO-thresholded graphs by setting a proportional threshold such that the average node degree would be 3, since previous analytic and empirical results indicate that the optimal density corresponds to enforcing an average node degree approximately equal to 3^56^.

### Orthogonal Minimum Spanning Trees

OMST^55, 80^ is another data-driven approach intended to optimise the balance between efficiency and density of the network, while also ensuring that the network is fully connected. Specifically, the method involves three steps: (1) identifying the minimum set of edges such that each node can be reached from each other node – known as the minimum spanning tree (MST); (2) identifying an alternative (orthogonal) MST, and combining it with the previous one; (3) repeating steps (1) and (2) until the network formed by the progressive addition of orthogonal MSTs optimises a global cost function defined as *E_g_* – Cost (with Cost corresponding to the ratio of the total weight of the selected edges, divided by the total strength of the original fully weighted graph). This approach produces plausibly sparse networks without imposing an a-priori level across all subjects, and it has been shown that the resulting networks provide higher recognition accuracy and reliability than many alternative filtering schemes^55, 80^.

### Binarisation

For all filtering schemes considered here, edges that were not selected were set to zero. However, edges that were included in the network could be weighted or unweighted. In the case of unweighted (binary) networks, we set all non-zero edges to unity. Otherwise, their original weight was retained.

### Topological distance as Portrait Divergence

To quantify the difference between network topologies, we used the recently developed Portrait Divergence. The Portrait Divergence between two graphs G_1_ and G_2_ is the Jensen-Shannon divergence between their “network portraits”, which encode the distribution of shortest paths of the two networks^46^. Specifically, the network portrait is a matrix *B* whose entry *B_lk_, l* = 0, 1, …, *d* (with *d* being the graph diameter), *k* = 0, 1, …, *N* – 1, is the number of nodes having *k* nodes at shortest-path distance *l*.

Thus, to compute the Portrait Divergence one needs to compute the probability *P(k, l)* (and similarly *Q(k, l)* for the second graph) of randomly choosing two nodes at distance *l* and, for one of the two nodes, to have *k* nodes at distance *l*:

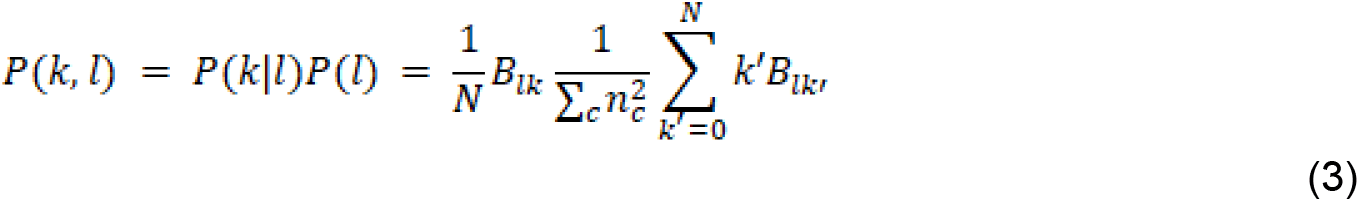

where *n_c_* is the number of nodes in the connected component *c*. Then, the Portrait Divergence distance is defined using the Jensen-Shannon divergence (an information-theoretic notion of distance):

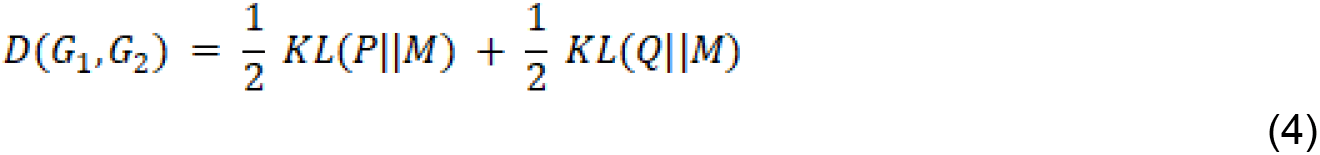

where *M = (P + Q)/2* is the mixture distribution of *P* and *Q*, and *KL*(⋅||⋅) is the Kullback-Leibler divergence.

The Portrait Divergence offers three key advantages that make it well suited for the present investigation. First, it is based on network portraits, which do not change depending on how a graph is represented. Comparing network topologies based on such “graph invariants” is highly desirable, because it removes the potential confound of encoding format. Second, the Portrait Divergence does not require the networks in question to have the same number of nodes or edges, and it can be applied to both binary and weighted networks – making it ideally suited for the applications of the present study. And finally, the Portrait Divergence is not predicated on a single specific network property, but rather it takes into account all scales of structure within networks, from local structure to motifs to large scale connectivity patterns: that is, it considers the topology of the network as a whole^46^.

For each subject, at each timepoint, we obtained one brain network following each of the possible combinations of steps above (576 distinct pipelines in total).

For each pipeline, we then computed the Portrait Divergence between networks obtained from the same subject at different points in time, and subsequently obtained a group-average value of Portrait Divergence for each pipeline.

## Supporting information

Supplementary Interactive Tool

## Acknowledgements

This work was supported by the Gates Cambridge Trust (OPP 1144) [AIL]; the Canadian Institute for Advanced Research (CIFAR; grant RCZB/072 RG93193) [to DKM and EAS]; The National Institute for Health Research (NIHR, UK), Cambridge Biomedical Research Centre and NIHR Senior Investigator Awards [DKM]; The British Oxygen Professorship of the Royal College of Anaesthetists [DKM]; The Stephen Erskine Fellowship, Queens’ College, University of Cambridge [EAS]; the Wellcome Trust Research Training Fellowship (grant no. 083660/Z/07/Z), Raymond and Beverly Sackler Studentship, and the Cambridge Commonwealth Trust [to RA]; the Medical Research Council Doctoral Training Grant (#RG86932) [HMG]; Pinsent Darwin Award [HMG]; MRC grant MR/K004360/1 [SID], a Marie Sklodowska-Curie COFUND EU-UK Research Fellowship [SID], a Beatriu de Pinós fellowship (2020 BP 00116) [SID]; AMO is supported by the Canada Excellence Research Chairs program (215063); LN acknowledges support by the L’Oreal-Unesco for Women in Science Excellence Research Fellowship; ZQL is supported by the Fonds de Recherche du Quebec – Nature et Technologies (FRQNT). Acquisition of the NYU Test-Retest dataset was funded by Stavros S. Niarchos Foundation, the Leon Lowenstein Foundation, NARSAD (The Mental Health Research Association) grants to F.Xavier Castellanos; and Linda and Richard Schaps, Jill and Bob Smith, and the Taubman Foundation gifts to F. Xavier Castellanos. This work was performed using resources provided by the Cambridge Service for Data Driven Discovery (CSD3) operated by the University of Cambridge Research Computing Service (www.csd3.cam.ac.uk), provided by Dell EMC and Intel using Tier-2 funding from the Engineering and Physical Sciences Research Council (capital grant EP/T022159/1), and DiRAC funding from the Science and Technology Facilities Council (www.dirac.ac.uk). For the purpose of open access, the authors have applied a Creative Commons Attribution (CC BY) licence to any Author Accepted Manuscript version arising from this submission.

## Author contributions

AIL, EAS, SID conceived the study. AIL, HMG, SID and EAS designed the methodology and the analysis. ZQL and ARDP contributed to data analysis. AIL and HMG analysed the data. EAS, AEM, RA, AMO, LN participated in data collection. AIL, SID, HMG wrote the manuscript with feedback from all co-authors.

## Conflicts of interest

The authors declare that no conflicts of interest exist.

## Data and code availability

The NYU dataset is freely available from the International Neuroimaging Data-Sharing Initiative (INDI) (http://www.nitrc.org/projects/nyu_trt). The Cambridge datasets are available upon request from author EAS. The HCP data are available from https://www.humanconnectome.org/. The CONN toolbox is freely available online (http://www.nitrc.org/projects/conn). Python code for the portrait divergence is freely available online (https://github.com/bagrow/network-portrait-divergence). MATLAB code for the Orthogonal Minimum Spanning Tree thresholding is freely available online (https://github.com/stdimitr/topological_filtering_networks). The Brain Connectivity Toolbox code used for graph-theoretical analyses is freely available online (https://sites.google.com/site/bctnet/).

## Notes

### Competing Interest Statement

The authors have declared no competing interest.

## References

1. Bullmore, E. & Sporns, O. Complex brain networks: graph theoretical analysis of structural and functional systems. Nature Reviews Neuroscience 10, 312–312 (2009).

2. Fox, M. D., Snyder, A. Z., Vincent, J. L., Corbetta, M., Van Essen, D. C. & Raichle, M. E. The human brain is intrinsically organized into dynamic, anticorrelated functional networks. Proceedings of the National Academy of Sciences 102, 9673–9678 (2005).

3. Park, H. J. & Friston, K. Structural and functional brain networks: From connections to cognition. Science 342, (2013).

4. Sporns, Olaf. Networks of the brain. (MIT Press, 2011).

5. Petersen, S. E. & Sporns, O. Brain Networks and Cognitive Architectures. Neuron 88, 207–219 (2015).

6. Bassett, D. S. & Sporns, O. Network neuroscience. Nature Neuroscience 20, 353–364 (2017).

7. Fornito, A., Zalesky, A. & Breakspear, M. The connectomics of brain disorders. Nature Reviews Neuroscience 16, 159–172 (2015).

8. Smith, S. M. et al. A positive-negative mode of population covariation links brain connectivity, demographics and behavior. Nature Neuroscience 18, 1–7 (2015).

9. Perovnik, M., Rus, T., Schindlbeck, K. A. & Eidelberg, D. Functional brain networks in the evaluation of patients with neurodegenerative disorders. Nat Rev Neurol 19, 73–90 (2023).

10. Smith, S. M. et al. Functional connectomics from resting-state fMRI. Trends in cognitive sciences 17, 666–82 (2013).

11. Turk, E. et al. Functional Connectome of the Fetal Brain. The Journal of neuroscience : the official journal of the Society for Neuroscience 39, 9716–9724 (2019).

12. Demertzi, A. et al. Human consciousness is supported by dynamic complex patterns of brain signal coordination. Science Advances 5, 1–12 (2019).

13. Huang, Z., Zhang, J., Wu, J., Mashour, G. A. & Hudetz, A. G. Temporal circuit of macroscale dynamic brain activity supports human consciousness. Science Advances 6, 87–98 (2020).

14. Luppi, A. I. et al. Consciousness-specific dynamic interactions of brain integration and functional diversity. Nature Communications 10, (2019).

15. Burianová, H., Faizo, N. L., Gray, M., Hocking, J., Galloway, G. & Reutens, D. Altered functional connectivity in mesial temporal lobe epilepsy. Epilepsy Res 137, 45–52 (2017).

16. Holiga, Š., et al. Patients with autism spectrum disorders display reproducible functional connectivity alterations. Sci Transl Med 11, eaat9223 (2019).

17. Karbasforoushan, H. & Woodward, N. D. Resting-state networks in schizophrenia. Curr Top Med Chem 12, 2404–2414 (2012).

18. Hallquist, M. N. & Hillary, F. G. Graph theory approaches to functional network organization in brain disorders: A critique for a brave new small-world. Network Neuroscience 3, 1–26 (2019).

19. Petrella, J. R. Use of graph theory to evaluate brain networks: a clinical tool for a small world? Radiology 259, 317–320 (2011).

20. Botvinik-Nezer, R. et al. Variability in the analysis of a single neuroimaging dataset by many teams. Nature 582, 84–88 (2020).

21. Eickhoff, S. B., Yeo, B. T. T. & Genon, S. Imaging-based parcellations of the human brain. Nature Reviews Neuroscience 19, 672–686 (2018).

22. Carp, J. On the plurality of (methodological) worlds: Estimating the analytic flexibility of fmri experiments. Frontiers in Neuroscience (2012) doi:10.3389/fnins.2012.00149.

23. Parkes, L., Fulcher, B., Yücel, M. & Fornito, A. An evaluation of the efficacy, reliability, and sensitivity of motion correction strategies for resting-state functional MRI. NeuroImage 171, 415–436 (2018).

24. Toga, A. W., Thompson, P. M., Mori, S., Amunts, K. & Zilles, K. Towards multimodal atlases of the human brain. Nat Rev Neurosci 7, 952–966 (2006).

25. Arslan, S., Ktena, S. I., Makropoulos, A., Robinson, E. C., Rueckert, D. & Parisot, S. Human brain mapping: A systematic comparison of parcellation methods for the human cerebral cortex. NeuroImage 170, 5–30 (2018).

26. Genon, S., Reid, A., Langner, R., Amunts, K. & Eickhoff, S. B. How to Characterize the Function of a Brain Region. Trends in Cognitive Sciences 22, 350–364 (2018).

27. Bijsterbosch, J., Harrison, S. J., Jbabdi, S., Woolrich, M., Beckmann, C., Smith, S. & Duff, E. P. Challenges and future directions for representations of functional brain organization. Nature Neuroscience 23, 1484–1495 (2020).

28. Korhonen, O., Zanin, M. & Papo, D. Principles and open questions in functional brain network reconstruction. Human Brain Mapping (2021) doi:10.1002/hbm.25462.

29. Woo, C.-W., Chang, L. J., Lindquist, M. A. & Wager, T. D. Building better biomarkers: brain models in translational neuroimaging. Nat Neurosci 20, 365–377 (2017).

30. Messaritaki, E., Dimitriadis, S. I. & Jones, D. K. Optimization of graph construction can significantly increase the power of structural brain network studies. NeuroImage 199, 495–511 (2019).

31. Nichols, T. E. et al. Best practices in data analysis and sharing in neuroimaging using MRI. Nat Neurosci 20, 299–303 (2017).

32. Chen, X., Liao, X., Dai, Z., Lin, Q., Wang, Z., Li, K. & He, Y. Topological analyses of functional connectomics: A crucial role of global signal removal, brain parcellation, and null models. Hum Brain Mapp 39, 4545–4564 (2018).

33. Saad, Z. S., Gotts, S. J., Murphy, K., Chen, G., Jo, H. J., Martin, A. & Cox, R. W. Trouble at Rest: How Correlation Patterns and Group Differences Become Distorted After Global Signal Regression. Brain Connectivity 2, 25–32 (2012).

34. Andellini, M., Cannatà, V., Gazzellini, S., Bernardi, B. & Napolitano, A. Test-retest reliability of graph metrics of resting state MRI functional brain networks: A review. Journal of Neuroscience Methods 253, 183–192 (2015).

35. Braun, U. et al. Test-retest reliability of resting-state connectivity network characteristics using fMRI and graph theoretical measures. NeuroImage 59, 1404–1412 (2012).

36. Cao, H. et al. Test-retest reliability of fMRI-based graph theoretical properties during working memory, emotion processing, and resting state. NeuroImage 84, 888–900 (2014).

37. Du, H.-X., Liao, X.-H., Lin, Q.-X., Li, G.-S., Chi, Y.-Z., Liu, X., Yang, H.-Z., Wang, Y. & Xia, M.-R. Test-retest reliability of graph metrics in high-resolution functional connectomics: a resting-state functional MRI study. CNS Neurosci Ther 21, 802–816 (2015).

38. Romero-Garcia, R., Atienza, M., Clemmensen, L. H. & Cantero, J. L. Effects of network resolution on topological properties of human neocortex. NeuroImage 59, 3522–3532 (2012).

39. Termenon, M., Jaillard, A., Delon-Martin, C. & Achard, S. Reliability of graph analysis of resting state fMRI using test-retest dataset from the Human Connectome Project. NeuroImage 142, 172–187 (2016).

40. Wang, J.-H., Zuo, X.-N. X., Gohel, S., Milham, M. P., Biswal, B. B. & He, Y. Graph theoretical analysis of functional brain networks: test-retest evaluation on short- and long-term resting-state functional MRI data. PloS one 6, e21976 (2011).

41. Welton, T., Kent, D. A., Auer, D. P. & Dineen, R. A. Reproducibility of Graph-Theoretic Brain Network Metrics: A Systematic Review. Brain Connectivity 5, 193–202 (2015).

42. Tozzi, L., Fleming, S. L., Taylor, Z. D., Raterink, C. D. & Williams, L. M. Test-retest reliability of the human functional connectome over consecutive days: identifying highly reliable portions and assessing the impact of methodological choices. Network Neuroscience 4, 925–945 (2020).

43. Jiang, C., Betzel, R., He, Y., Wang, Y.-S., Xing, X.-X. & Zuo, X.-N. Toward Reliable Network Neuroscience for Mapping Individual Differences. 2021.05.06.442886 Preprint at https://doi.org/10.1101/2021.05.06.442886 (2021).

44. Rubinov, M. & Sporns, O. Complex network measures of brain connectivity: Uses and interpretations. NeuroImage 52, 1059–1069 (2010).

45. Rubinov, M. & Sporns, O. Weight-conserving characterization of complex functional brain networks. NeuroImage 56, 2068–2079 (2011).

46. Bagrow, J. P. & Bollt, E. M. An information-theoretic, all-scales approach to comparing networks. Applied Network Science 4, 45 (2019).

47. Burgess, G. C., Kandala, S., Nolan, D., Laumann, T. O., Power, J. D., Adeyemo, B., Harms, M. P., Petersen, S. E. & Barch, D. M. Evaluation of Denoising Strategies to Address Motion-Correlated Artifacts in Resting-State Functional Magnetic Resonance Imaging Data from the Human Connectome Project. Brain Connectivity 6, 669–680 (2016).

48. Power, J. D., Schlaggar, B. L. & Petersen, S. E. Recent progress and outstanding issues in motion correction in resting state fMRI. (2015) doi:10.1016/j.neuroimage.2014.10.044.

49. Power, J. D., Mitra, A., Laumann, T. O., Snyder, A. Z., Schlaggar, B. L. & Petersen, S. E. Methods to detect, characterize, and remove motion artifact in resting state fMRI. NeuroImage 84, 320–341 (2014).

50. Ciric, R. et al. Benchmarking of participant-level confound regression strategies for the control of motion artifact in studies of functional connectivity. NeuroImage 154, 174–187 (2017).

51. Salimi-Khorshidi, G., Douaud, G., Beckmann, C. F., Glasser, M. F., Griffanti, L. & Smith, S. M. Automatic denoising of functional MRI data: Combining independent component analysis and hierarchical fusion of classifiers. NeuroImage 90, 449–468 (2014).

52. Griffanti, L. et al. ICA-based artefact removal and accelerated fMRI acquisition for improved resting state network imaging. NeuroImage 95, 232–247 (2014).

53. Murphy, K. & Fox, M. D. Towards a consensus regarding global signal regression for resting state functional connectivity MRI. NeuroImage 154, 169–173 (2017).

54. Luppi, A. I. & Stamatakis, E. A. Combining network topology and information theory to construct representative brain networks. Network Neuroscience 5, 96–124 (2021).

55. Dimitriadis, S. I., Antonakakis, M., Simos, P., Fletcher, J. M. & Papanicolaou, A. C. Data-Driven Topological Filtering Based on Orthogonal Minimal Spanning Trees: Application to Multigroup Magnetoencephalography Resting-State Connectivity. Brain Connectivity 7, 661–670 (2017).

56. De Vico Fallani, F., Latora, V. & Chavez, M. A Topological Criterion for Filtering Information in Complex Brain Networks. PLoS Computational Biology 13, (2017).

57. Glasser, M. F. et al. The Minimal Preprocessing Pipelines for the Human Connectome Project. Neuroimage 80, 105–124 (2013).

58. Van Essen, D. C., Smith, S. M., Barch, D. M., Behrens, T. E. J., Yacoub, E. & Ugurbil, K. The WU-Minn Human Connectome Project: An overview. NeuroImage 80, 62–79 (2013).

59. Smith, S. M. et al. Resting-state fMRI in the Human Connectome Project. NeuroImage 80, 144–168 (2013).

60. Helwegen, K., Libedinsky, I. & Heuvel, M. P. van den. Statistical power in network neuroscience. Trends in Cognitive Sciences 27, 282–301 (2023).

61. Shafto, M. A. et al. The Cambridge Centre for Ageing and Neuroscience (Cam-CAN) study protocol: A cross-sectional, lifespan, multidisciplinary examination of healthy cognitive ageing. BMC Neurology 14, 1–25 (2014).

62. Satterthwaite, T. D. et al. The Philadelphia Neurodevelopmental Cohort: A publicly available resource for the study of normal and abnormal brain development in youth. NeuroImage 124, 1115–1119 (2016).

63. Maas, A. I. R., Menon, D. K., Steyerberg, E. W., Citerio, G., Lecky, F., Manley, G., Hill, S., Legrand, V. & Sorgner, A. Collaborative European NeuroTrauma Effectiveness Research in Traumatic Brain Injury (CENTER-TBI): A Prospective Longitudinal Observational Study. Neurosurgery 76, 67–80 (2014).

64. 64. Dagley, A. et al. Harvard Aging Brain Study: dataset and accessibility. Neuroimage 144, 255–258 (2017).

65. Di Martino, A. et al. The autism brain imaging data exchange: towards a large-scale evaluation of the intrinsic brain architecture in autism. Mol Psychiatry 19, 659–667 (2014).

66. Poldrack, R. A. et al. A phenome-wide examination of neural and cognitive function. Scientific Data 2016 3:1 3, 1–12 (2016).

67. Littlejohns, T. J. et al. The UK Biobank imaging enhancement of 100,000 participants: rationale, data collection, management and future directions. Nat Commun 11, 2624 (2020).

68. Friston, K. J., Harrison, L. & Penny, W. Dynamic causal modelling. NeuroImage 19, 1273–1302 (2003).

69. Deco, G., Ponce-Alvarez, A., Hagmann, P., Romani, G. L., Mantini, D. & Corbetta, M. How local excitation-inhibition ratio impacts the whole brain dynamics. Journal of Neuroscience 34, 7886–7898 (2014).

70. Honey, C. J., Sporns, O., Cammoun, L., Gigandet, X., Thiran, J. P., Meuli, R. & Hagmann, P. Predicting human resting-state functional connectivity from structural connectivity. Proceedings of the National Academy of Sciences 106, 2035–2040 (2009).

71. Saggio, M. L., Ritter, P. & Jirsa, V. K. Analytical Operations Relate Structural and Functional Connectivity in the Brain. PLOS ONE 11, e0157292 (2016).

72. 72. Schulz, M. A., Yeo, B. T. T., Vogelstein, J. T., Mourao-Miranada, J., Kather, J. N., Kording, K., Richards, B. & Bzdok, D. Different scaling of linear models and deep learning in UK Biobank brain images vs. machine-learning datasets. Nature Communications 11, (2020).

73. Nozari, E., Stiso, J., Caciagli, L., Cornblath, E. J., He, X., Bertolero, M. A., Mahadevan, A. S., Pappas, G. J. & Bassett, D. S. Is the brain macroscopically linear? A system identification of resting state dynamics. bioRxiv (2020) doi: 10.1101/2020.12.21.423856.

74. Ahmed, S. & Nozari, E. On the Linearizing Effect of Spatial Averaging in Large-Scale Populations of Homogeneous Nonlinear Systems. in 2022 IEEE 61st Conference on Decision and Control (CDC) 641–648 (2022). doi:10.1109/CDC51059.2022.9993260.

75. Esfahlani, F. Z. & Sayama, H. A Percolation-Based Thresholding Method with Applications in Functional Connectivity Analysis. in Complex Networks IX (eds. Cornelius, S., Coronges, K., Gonçalves, B., Sinatra, R. & Vespignani, A.) 221–231 (Springer International Publishing, 2018). doi:10.1007/978-3-319-73198-8_19.

76. Bordier, C., Nicolini, C. & Bifone, A. Graph analysis and modularity of brain functional connectivity networks: Searching for the optimal threshold. Frontiers in Neuroscience 11, 1–9 (2017).

77. Gallos, L. K., Makse, H. A. & Sigman, M. A small world of weak ties provides optimal global integration of self-similar modules in functional brain networks. Proc. Nati. Acad. Sci. USA 109, 2825–2830 (2012).

78. Gallos, L. K., Sigman, M. & Makse, H. A. The conundrum of functional brain networks: Small-world efficiency or fractal modularity. Frontiers in Physiology 3, (2012).

79. Granovetter, M. S. The Strength of Weak Ties. American Journal of Sociology 78, 1360– 1380 (1973).

80. Dimitriadis, S. I., Salis, C., Tarnanas, I. & Linden, D. E. Topological filtering of dynamic functional brain networks unfolds informative chronnectomics: A novel data-driven thresholding scheme based on orthogonal minimal spanning trees (OMSTs). Frontiers in Neuroinformatics 11, 28 (2017).

81. Craddock, R. C., James, G. A., Holtzheimer, P. E., Hu, X. P. & Mayberg, H. S. A whole brain fMRI atlas generated via spatially constrained spectral clustering. Human Brain Mapping 33, 1914–1928 (2012).

82. Craddock, R. C. et al. Imaging human connectomes at the macroscale. Nature Methods 10, 524–539 (2013).

83. Margulies, D. S. et al. Situating the default-mode network along a principal gradient of macroscale cortical organization. Proceedings of the National Academy of Sciences of the United States of America 113, 12574–12579 (2016).

84. Atasoy, S., Donnelly, I. & Pearson, J. Human brain networks function in connectome-specific harmonic waves. Nature Communications 7, 1–10 (2016).

85. Glomb, K., Kringelbach, M. L., Deco, G., Hagmann, P., Pearson, J. & Atasoy, S. Functional harmonics reveal multi-dimensional basis functions underlying cortical organization. Cell Reports 36, 109554 (2021).

86. Lioi, G., Gripon, V., Brahim, A., Rousseau, F. & Farrugia, N. Gradients of connectivity as graph Fourier bases of brain activity. Network Neuroscience 5, 322–336 (2021).

87. Thirion, B., Varoquaux, G., Dohmatob, E. & Poline, J.-B. Which fMRI clustering gives good brain parcellations? Front Neurosci 8, 167 (2014).

88. Kiviniemi, V. et al. Functional segmentation of the brain cortex using high model order group PICA. Human Brain Mapping 30, 3865–3886 (2009).

89. Smith, S. M., Miller, K. L., Salimi-Khorshidi, G., Webster, M., Beckmann, C. F., Nichols, T. E., Ramsey, J. D. & Woolrich, M. W. Network modelling methods for FMRI. NeuroImage 54, 875–891 (2011).

90. Yu, Q., Du, Y., Chen, J., He, H., Sui, J., Pearlson, G. & Calhoun, V. D. Comparing brain graphs in which nodes are regions of interest or independent components: A simulation study. J Neurosci Methods 291, 61–68 (2017).

91. Váša, F., Bullmore, E. T. & Patel, A. X. Probabilistic thresholding of functional connectomes: Application to schizophrenia. NeuroImage 172, 326–340 (2018).

92. Wang, M. B., Owen, J. P., Mukherjee, P. & Raj, A. Brain network eigenmodes provide a robust and compact representation of the structural connectome in health and disease. PLoS Computational Biology 13, (2017).

93. Yoo, K., Rosenberg, M. D., Noble, S., Scheinost, D., Constable, R. T. & Chun, M. M. Multivariate approaches improve the reliability and validity of functional connectivity and prediction of individual behaviors. NeuroImage 197, 212–223 (2019).

94. Mediano, P. A. M., Rosas, F. E., Luppi, A. I., Carhart-Harris, R. L., Bor, D., Seth, A. K. & Barrett, A. B. Towards an extended taxonomy of information dynamics via Integrated Information Decomposition. arXiv (2021).

95. Luppi, A. I. et al. A synergistic core for human brain evolution and cognition. Nat Neurosci 25, 771–782 (2022).

96. Varley, T. F., Sporns, O., Schaffelhofer, S., Scherberger, H. & Dann, B. Information-processing dynamics in neural networks of macaque cerebral cortex reflect cognitive state and behavior. Proc Natl Acad Sci U S A 120, e2207677120 (2023).

97. Friston, K. J., Preller, K. H., Mathys, C., Cagnan, H., Heinzle, J., Razi, A. & Zeidman, P. Dynamic causal modelling revisited. NeuroImage 199, 730–744 (2019).

98. Hutchison, R. M. et al. Dynamic functional connectivity: Promise, issues, and interpretations. Neuroimage 80, 5–79 (2013).

99. Preti, M. G., Bolton, T. A. & Van De Ville, D. The dynamic functional connectome: State-of-the-art and perspectives. NeuroImage 160, 41–54 (2017).

100. Lurie, D. J. et al. Questions and controversies in the study of time-varying functional connectivity in resting fMRI. Network Neuroscience 4, 30–69 (2020).

101. Pourmotabbed, H., de Jongh Curry, A. L., Clarke, D. F., Tyler-Kabara, E. C. & Babajani-Feremi, A. Reproducibility of graph measures derived from resting-state MEG functional connectivity metrics in sensor and source spaces. Human Brain Mapping 43, 1342–1357 (2022).

102. Dimitriadis, S. I. Complexity of brain activity and connectivity in functional neuroimaging. J Neuro Res 96, 1741–1757 (2018).

101. Dimitriadis, S. I. Assessing The Repeatability of Multi-Frequency Multi-Layer Brain Network Topologies Across Alternative Researcher’s Choice Paths. bioRxiv 2021.10.10.463799 (2021) doi: 10.1101/2021.10.10.463799.

104. Liu, T. T., Nalci, A. & Falahpour, M. The Global Signal in fMRI: Nuisance or Information? Neuroimage 150, 213–229 (2017).

105. Tanabe, S. et al. Altered Global Brain Signal during Physiologic, Pharmacologic, and Pathologic States of Unconsciousness in Humans and Rats. Anesthesiology 1392–1406 (2020) doi:10.1097/ALN.0000000000003197.

106. Li, J., Bolt, T., Bzdok, D., Nomi, J. S., Yeo, B. T. T., Spreng, R. N. & Uddin, L. Q. Topography and behavioral relevance of the global signal in the human brain. Scientific Reports 9, 1–10 (2019).

107. Van Dijk, K. R. A., Hedden, T., Venkataraman, A., Evans, K. C., Lazar, S. W. & Buckner, R. L. Intrinsic functional connectivity as a tool for human connectomics: theory, properties, and optimization. J Neurophysiol 103, 297–321 (2010).

108. Laumann, T. O. et al. On the Stability of BOLD fMRI Correlations. Cerebral Cortex 27, 4719–4732 (2017).

109. Shah, L. M., Cramer, J. A., Ferguson, M. A., Birn, R. M. & Anderson, J. S. Reliability and reproducibility of individual differences in functional connectivity acquired during task and resting state. Brain Behav 6, e00456 (2016).

110. Kristo, G., Rutten, G., Raemaekers, M., de Gelder, B., Rombouts, S. A. R. B. & Ramsey, N. F. Task and task-free FMRI reproducibility comparison for motor network identification. Hum Brain Mapp 35, 340–352 (2012).

111. Somandepalli, K. et al. Short-term test-retest reliability of resting state fMRI metrics in children with and without attention-deficit/hyperactivity disorder. Dev Cogn Neurosci 15, 83–93 (2015).

112. Conwell, K., von Reutern, B., Richter, N., Kukolja, J., Fink, G. R. & Onur, O. A. Test-retest variability of resting-state networks in healthy aging and prodromal Alzheimer’s disease. Neuroimage Clin 19, 948–962 (2018).

111. 113. Song, J. et al. Age-related differences in test-retest reliability in resting-state brain functional connectivity. PLoS One 7, e49847 (2012).

114. Dafflon, J. et al. A guided multiverse study of neuroimaging analyses. Nat Commun 13, 3758 (2022).

115. Shehzad, Z. et al. The resting brain: unconstrained yet reliable. Cereb Cortex 19, 2209–2229 (2009).

116. Vatansever, D., Menon, X. D. K., Manktelow, A. E., Sahakian, B. J. & Stamatakis, E. A. Default Mode Dynamics for Global Functional Integration. The Journal of neuroscience : the official journal of the Society for Neuroscience 35, 15254–15262 (2015).

117. Manktelow, A. E., Menon, D. K., Sahakian, B. J. & Stamatakis, E. A. Working Memory after Traumatic Brain Injury: The Neural Basis of Improved Performance with Methylphenidate. Frontiers in Behavioral Neuroscience 11, (2017).

118. Stamatakis, E. A., Adapa, R. M., Absalom, A. R. & Menon, D. K. Changes in resting neural connectivity during propofol sedation. PloS one 5, e14224 (2010).

119. Adapa, R. M., Davis, M. H., Stamatakis, E. A., Absalom, A. R. & Menon, D. K. Neural Correlates of Successful Semantic Processing During Propofol Sedation. Human Brain Mapping (2013) doi:10.1002/hbm.22375.

120. Varley, T. F., Luppi, A. I., Pappas, I., Naci, L., Adapa, R., Owen, A. M., Menon, D. K. & Stamatakis, E. A. Consciousness & Brain Functional Complexity in Propofol Anaesthesia. Scientific Reports 10, 1–13 (2020).

121. Naci, L., Sinai, L. & Owen, A. M. Detecting and interpreting conscious experiences in behaviorally non-responsive patients. NeuroImage 145, 304–313 (2017).

122. Kandeepan, S., Rudas, J., Gomez, F., Stojanoski, B., Valluri, S., Owen, A. M., Naci, L., Nichols, E. S. & Soddu, A. Modeling an auditory stimulated brain under altered states of consciousness using the generalized ising model. NeuroImage 223, 117367 (2020).

123. Whitfield-Gabrieli, S. & Nieto-Castanon, A. Conn: A Functional Connectivity Toolbox for Correlated and Anticorrelated Brain Networks. Brain Connectivity 2, 125–141 (2012).

124. Behzadi Y, Restom K, Liau J, & Liu TT. A component based noise correction method (CompCor) for BOLD and perfusion based fMRI. NeuroImage 37, 90–101 (2007).

125. Messé, A. Parcellation influence on the connectivity-based structure–function relationship in the human brain. Human Brain Mapping 41, 1167–1180 (2020).

126. Popovych, O. V., Jung, K., Manos, T., Diaz-Pier, S., Hoffstaedter, F., Schreiber, J., Yeo, B. T. T. & Eickhoff, S. B. Inter-subject and inter-parcellation variability of resting-state whole-brain dynamical modeling. NeuroImage 118201 (2021) doi:10.1016/j.neuroimage.2021.118201.

127. Cammoun, L., Gigandet, X., Meskaldji, D., Thiran, J. P., Sporns, O., Do, K. Q., Maeder, P., Meuli, R. & Hagmann, P. Mapping the human connectome at multiple scales with diffusion spectrum MRI. Journal of Neuroscience Methods 203, 386–397 (2012).

128. Schaefer, A., Kong, R., Gordon, E. M., Laumann, T. O., Zuo, X.-N., Holmes, A. J., Eickhoff, S. B. & Yeo, B. T. T. Local-Global Parcellation of the Human Cerebral Cortex from Intrinsic Functional Connectivity MRI. Cerebral Cortex 28, 3095–3114 (2018).

129. Tian, Y., Margulies, D., Breakspear, M. & Zalesky, A. Topographic organization of the human subcortex unveiled with functional connectivity gradients. Nature Neuroscience 23, 1421–1432 (2020).

130. Tzourio-Mazoyer, N., Landeau, B., Papathanassiou, D., Crivello, F., Etard, O., Delcroix, N., Mazoyer, B. & Joliot, M. Automated Anatomical Labeling of Activations in SPM Using a Macroscopic Anatomical Parcellation of the MNI MRI Single-Subject Brain. NeuroImage 15, 273–289 (2002).

131. Fan, L. et al. The Human Brainnetome Atlas: A New Brain Atlas Based on Connectional Architecture. Cerebral Cortex 26, 3508–3526 (2016).

132. Glasser, M. F. et al. A multi-modal parcellation of human cerebral cortex. Nature 536, 171–178 (2016).

133. Preti, M. G. & Van De Ville, D. Decoupling of brain function from structure reveals regional behavioral specialization in humans. Nature Communications 10, (2019).

134. Yeh, F.-C., Panesar, S., Fernandes, D., Meola, A., Yoshino, M., Fernandez-Miranda, J. C., Vettel, J. M. & Verstynen, T. Population-averaged atlas of the macroscale human structural connectome and its network topology. NeuroImage 178, 57–68 (2018).

