## Supplementary Material for "Converging on consistent functional connectomics"

Luppi *et al.*

#### Guide to pipeline selection in the *Supplementary Interactive Tool*

This document provides a guide for the use of the pipeline selection tool. The tool is in the form of an Excel file which allows the user to filter pipelines based on specific user-defined criteria. Pipelines can be filtered based on multiple criteria combined to allow the user to specify preferred preconditions for a pipeline choice. The criteria for pipeline selection:

- **Criterion (I):** Avoiding spurious differences ("PDiv ranking"). Since the two networks that we consider are derived from different scans of the same healthy individuals under conditions in which no experimentally meaningful changes in functional network topology are expected, we aim to identify pipelines that minimise test-retest PDiv. We consider pipelines as candidates for optimal if they are in the top 20% in terms of the global PDiv rank calculated across all four test-retest intervals.
- **Criterion (II):** Detecting true experimental differences ("propofol"). Suitable pipelines should detect a significant effect for propofol, in the right direction, in both propofol datasets, i.e., a pipeline is excluded if it fails to detect the expected effect in either of the two propofol datasets.
- **Criterion (III):** Detecting inter-individual differences ("within-between"). A pipeline fails this criterion if the resulting networks are more similar between than within subjects more than 50% of the times, for any of the three test-retest datasets.
- **Criterion (IV):** Avoiding motion-induced differences ("motion"). A pipeline fails this criterion if its PDiv has a significant correlation with differences in head motion in any of the three test-retest datasets.
- **Criterion (V):** Non-empty networks. As a final sanity check, we also exclude any pipelines that remove all connections from a network, in any of the three test-retest datasets.

Columns B and C mark pipelines which passed criteria II-V and were either in the top 50% pipelines based on average global rank or in the top 20%, respectively. Pipelines that fulfil all of these criteria can be selected by clicking the option "Selected" in the filter.

Combinations of multiple user-defined criteria can be obtained by selecting options in multiple filters at once. For instance, if the user wanted to identify all pipelines which fulfil the above five criteria, used a single scale atlas type and no global signal regression, this is what the result would look like (showing one pipeline which fulfils these criteria):

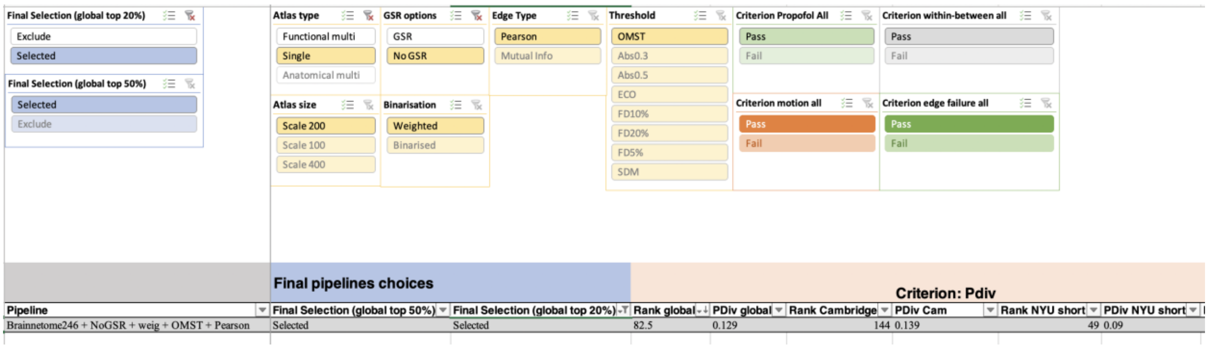


| Pipeline | Final Selection (global top 50%) | Final Selection (global top 20%) | Rank global | PDiv global | Rank Cambridge | PDiv Cam | Rank NYU short | PDiv NYU short |
|---|---|---|---|---|---|---|---|---|
| Brainnetome246 + NoGSR + weig + OMST + Pearson | Selected | Selected | 82.5 | 0.129 | 144 | 0.139 | 49 | 0.09 |

In contrast, if the user only cared about a pipeline passing Criteria II and V above, regardless of portrait divergence or pre-processing choices, the result may look as follows:

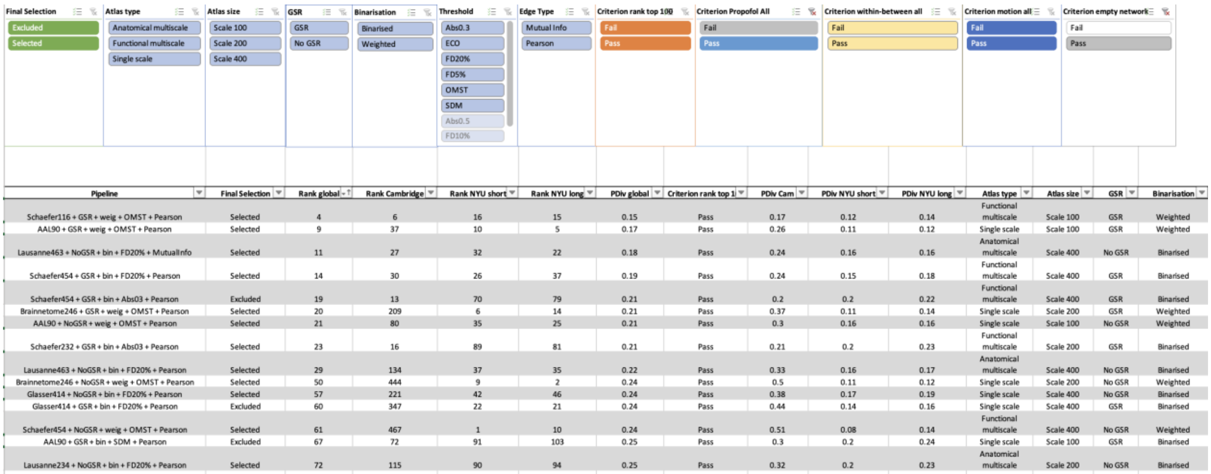


| Pipeline | Final Selection | Rank global | Rank Cambridge | Rank NYU short | Rank NYU long | PDiv global | Criterion rank top 1 | PDiv Cam | PDiv NYU short | PDiv NYU long | Atlas type | Atlas size | GSR | Binarisation |
|---|---|---|---|---|---|---|---|---|---|---|---|---|---|---|
| Schaefer116 + GSR + weig + OMST + Pearson | Selected | 4 | 6 | 16 | 15 | 0.15 | Pass | 0.17 | 0.12 | 0.14 | Functional multiscale | Scale 100 | GSR | Weighted |
| AAL90 + GSR + weig + OMST + Pearson | Selected | 9 | 37 | 10 | 5 | 0.17 | Pass | 0.26 | 0.11 | 0.12 | Single scale | Scale 100 | GSR | Weighted |
| Lausanne463 + NoGSR + bin + FD20% + MutualInfo | Selected | 11 | 27 | 32 | 22 | 0.18 | Pass | 0.24 | 0.16 | 0.16 | Anatomical multiscale | Scale 400 | No GSR | Binarised |
| Schaefer454 + GSR + bin + FD20% + Pearson | Selected | 14 | 30 | 26 | 37 | 0.19 | Pass | 0.24 | 0.15 | 0.18 | Functional multiscale | Scale 400 | GSR | Binarised |
| Schaefer454 + GSR + bin + Abs03 + Pearson | Excluded | 19 | 13 | 70 | 79 | 0.21 | Pass | 0.2 | 0.2 | 0.22 | Functional multiscale | Scale 400 | GSR | Binarised |
| Brainnetome246 + GSR + weig + OMST + Pearson | Selected | 20 | 209 | 6 | 14 | 0.21 | Pass | 0.37 | 0.11 | 0.14 | Single scale | Scale 200 | GSR | Weighted |
| AAL90 + NoGSR + weig + OMST + Pearson | Selected | 21 | 80 | 35 | 25 | 0.21 | Pass | 0.3 | 0.16 | 0.16 | Single scale | Scale 100 | No GSR | Weighted |
| Schaefer232 + GSR + bin + Abs03 + Pearson | Selected | 23 | 16 | 89 | 81 | 0.21 | Pass | 0.21 | 0.2 | 0.23 | Functional multiscale | Scale 200 | GSR | Binarised |
| Lausanne463 + NoGSR + bin + FD20% + Pearson | Selected | 29 | 134 | 37 | 35 | 0.22 | Pass | 0.33 | 0.16 | 0.17 | Anatomical multiscale | Scale 400 | No GSR | Binarised |
| Brainnetome246 + NoGSR + weig + OMST + Pearson | Selected | 50 | 444 | 9 | 2 | 0.24 | Pass | 0.5 | 0.11 | 0.12 | Single scale | Scale 200 | No GSR | Weighted |
| Glasser414 + NoGSR + bin + FD20% + Pearson | Selected | 57 | 221 | 42 | 46 | 0.24 | Pass | 0.38 | 0.17 | 0.19 | Single scale | Scale 400 | No GSR | Binarised |
| Glasser414 + GSR + bin + FD20% + Pearson | Excluded | 60 | 347 | 22 | 21 | 0.24 | Pass | 0.44 | 0.14 | 0.16 | Single scale | Scale 400 | GSR | Binarised |
| Schaefer454 + NoGSR + weig + OMST + Pearson | Selected | 61 | 467 | 1 | 10 | 0.24 | Pass | 0.51 | 0.08 | 0.14 | Functional multiscale | Scale 400 | No GSR | Weighted |
| AAL90 + GSR + bin + SDM + Pearson | Excluded | 67 | 72 | 91 | 103 | 0.25 | Pass | 0.3 | 0.2 | 0.24 | Single scale | Scale 100 | GSR | Binarised |
| Lausanne234 + NoGSR + bin + FD20% + Pearson | Selected | 72 | 115 | 90 | 94 | 0.25 | Pass | 0.32 | 0.2 | 0.23 | Anatomical multiscale | Scale 200 | No GSR | Binarised |

In this example, for the threshold slicer, options Abs0.5 and FD10% can now no longer be selected because no pipelines with these pre-processing choices fulfil the propofol and non-empty network criteria.

A reset can be achieved by clicking on the filter icon with the red cross in the upper right corner of a given slicer panel.

If the user wanted to include multiple options in a given slicer panel (for instance if all pipelines with atlas scale 200 and 400 were to be selected), the first option should be selected, followed by a click + command (or right click) on the second option. This would yield the following:

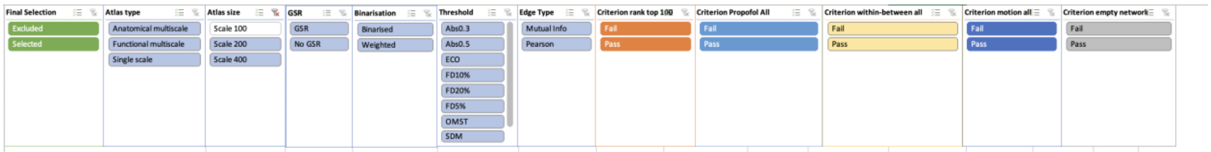


| Pipeline | Final Selection | Rank global | Rank Cambridge | Rank NYU short | Rank NYU long | PDiv global | Criterion rank top 1 | PDiv Cam | PDiv NYU short | PDiv NYU long | Atlas type | Atlas size | GSR | Binarisation |
|---|---|---|---|---|---|---|---|---|---|---|---|---|---|---|
| Schaefer454 + GSR + weig + OMST + Pearson | Excluded | 1 | 3 | 7 | 9 | 0.12 | Pass | 0.12 | 0.11 | 0.14 | Functional multiscale | Scale 400 | GSR | Weighted |
| Schaefer232 + GSR + weig + OMST + Pearson | Excluded | 2 | 7 | 2 | 3 | 0.13 | Pass | 0.18 | 0.09 | 0.12 | Functional multiscale | Scale 200 | GSR | Weighted |
| Schaefer232 + NoGSR + weig + OMST + Pearson | Excluded | 5 | 11 | 12 | 6 | 0.15 | Pass | 0.19 | 0.12 | 0.13 | Functional multiscale | Scale 200 | No GSR | Weighted |
| Lausanne234 + NoGSR + weig + OMST + Pearson | Excluded | 6 | 10 | 17 | 17 | 0.16 | Pass | 0.19 | 0.12 | 0.15 | Anatomical multiscale | Scale 200 | No GSR | Weighted |
| Schaefer454 + GSR + bin + FD20% + MutualInfo | Excluded | 10 | 15 | 39 | 43 | 0.18 | Pass | 0.21 | 0.16 | 0.18 | Functional multiscale | Scale 400 | GSR | Binarised |
| Lausanne463 + NoGSR + bin + FD20% + MutualInfo | Selected | 11 | 27 | 32 | 22 | 0.18 | Pass | 0.24 | 0.16 | 0.16 | Anatomical multiscale | Scale 400 | No GSR | Binarised |
| Schaefer232 + GSR + weig + Abs03 + Pearson | Excluded | 12 | 2 | 88 | 99 | 0.18 | Pass | 0.11 | 0.2 | 0.24 | Functional multiscale | Scale 200 | GSR | Weighted |
| Schaefer454 + GSR + bin + FD20% + Pearson | Selected | 14 | 30 | 26 | 37 | 0.19 | Pass | 0.24 | 0.15 | 0.18 | Functional multiscale | Scale 400 | GSR | Binarised |
| Schaefer454 + GSR + bin + Abs03 + Pearson | Excluded | 19 | 13 | 70 | 79 | 0.21 | Pass | 0.2 | 0.2 | 0.22 | Functional multiscale | Scale 400 | GSR | Binarised |
| Brainnetome246 + GSR + weig + OMST + Pearson | Selected | 20 | 209 | 6 | 14 | 0.21 | Pass | 0.37 | 0.11 | 0.14 | Single scale | Scale 200 | GSR | Weighted |
| Schaefer232 + GSR + bin + Abs03 + Pearson | Selected | 23 | 16 | 89 | 81 | 0.21 | Pass | 0.21 | 0.2 | 0.23 | Functional multiscale | Scale 200 | GSR | Binarised |
| Lausanne463 + NoGSR + bin + FD20% + Pearson | Selected | 29 | 134 | 37 | 35 | 0.22 | Pass | 0.33 | 0.16 | 0.17 | Anatomical multiscale | Scale 400 | No GSR | Binarised |
| Schaefer454 + GSR + bin + OMST + Pearson | Excluded | 30 | 22 | 79 | 117 | 0.22 | Pass | 0.23 | 0.2 | 0.24 | Functional multiscale | Scale 400 | GSR | Binarised |
| | | | | | | | | | | | Functional | | | |

Alternatively, filtering and sorting of the data based on any column available in the excel sheet can be done by clicking the downward facing arrow next to a column name in row 2.

### Supplementary Figures

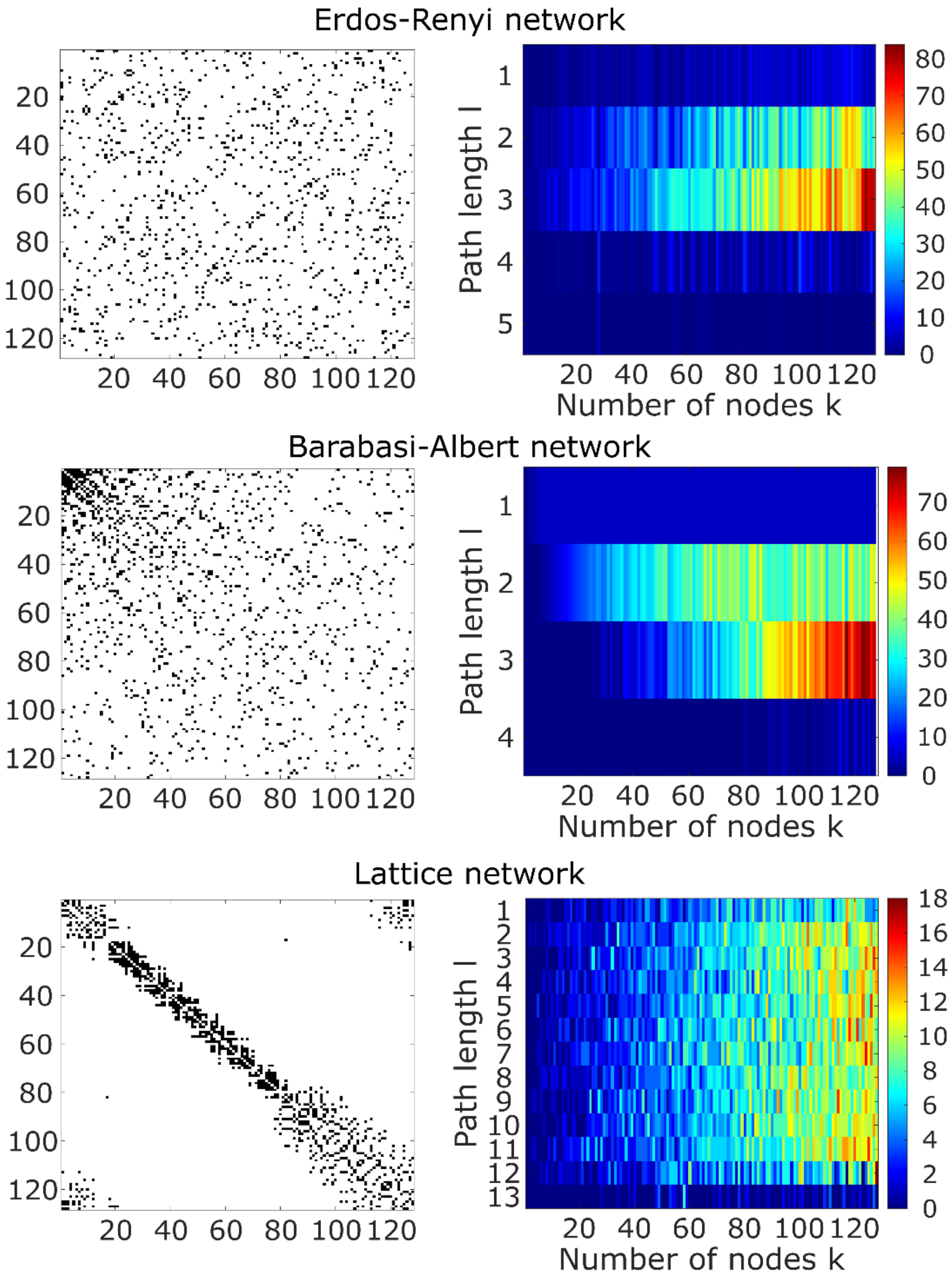


**Figure S1. Example networks (left) and their portraits (right).** From the top: Erdos-Renyi random network, Barabasi-Albert preferential attachment network, and lattice network. All networks are binary with an approximate density of 6%. A network portrait for a binary network is a matrix B whose rows each correspond to a histogram obtained by thresholding the matrix of shortest paths between the networks's constituent nodes, at each path length *l* between 0 and the network's diameter *L,* such that entry $B_{l,k}$ encodes the number of nodes that have *k* nodes at distance *l*. PDiv between ER and BA networks is 0.26; PDiv between ER and Lattice networks is 0.90; PDiv between BA and Lattice networks is 0.93.

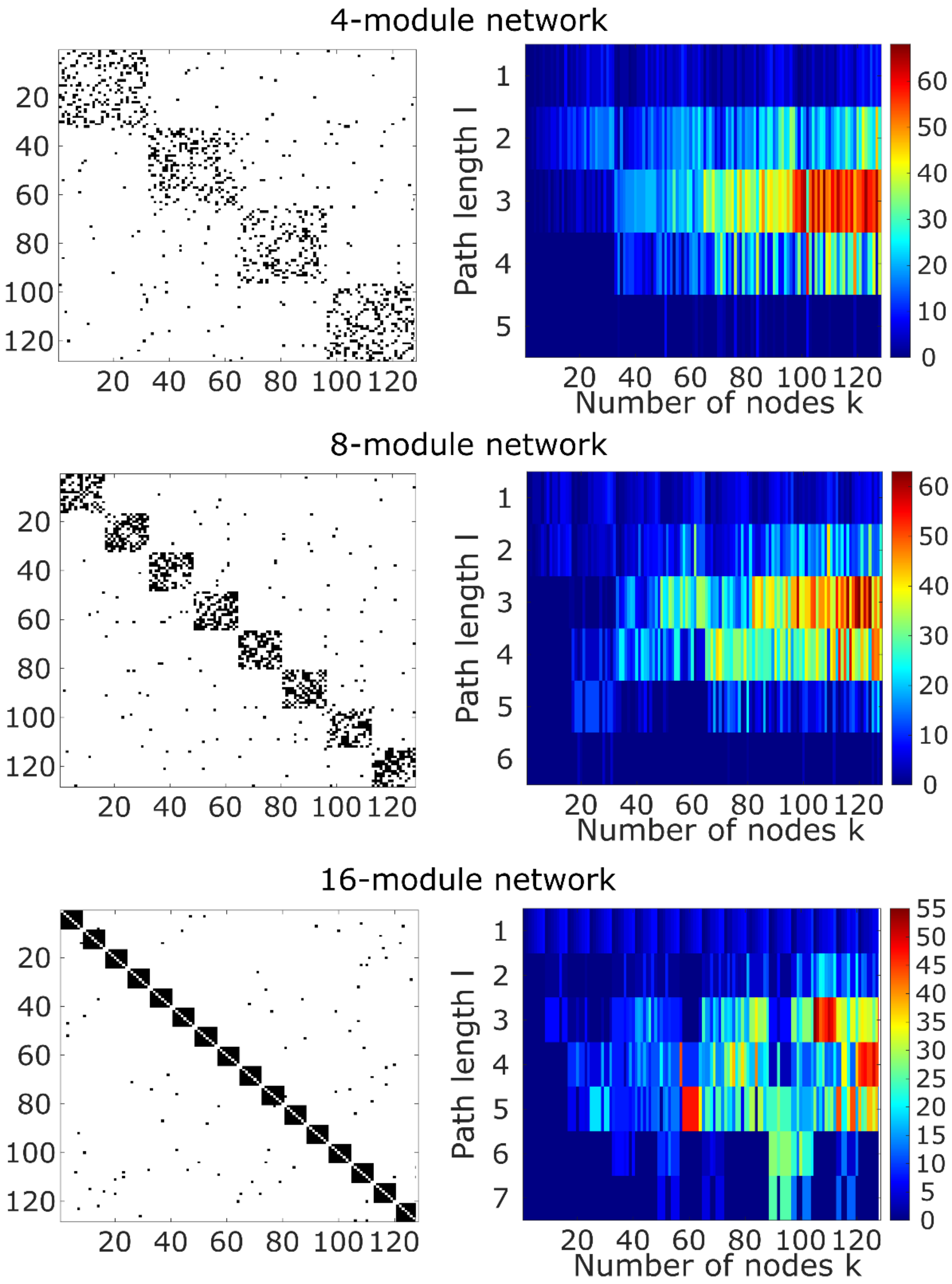


**Figure S2. Additional examples of networks (left) and their portraits (right).** From the top: modular networks with 4, 8, and 16 equal-sized modules, respectively. All networks are binary with an approximate density of 6%. A network portrait for a binary network is a matrix B whose rows each correspond to a histogram obtained by thresholding the matrix of shortest paths between the networks's constituent nodes, at each path length *l* between 0 and the network's diameter *L,* such that entry $B_{l,k}$ encodes the number of nodes that have *k* nodes at distance *l*. PDiv between the 4-module and 8-module networks is 0.36; PDiv between 8-module and 16-module networks is 0.52; PDiv between the 4-module and 16-module networks is 0.68. Note how the two most extreme cases (4 and 16 modules) have the largest PDiv, and how the modular organisation of each network is reflected in the first row of its network portrait.

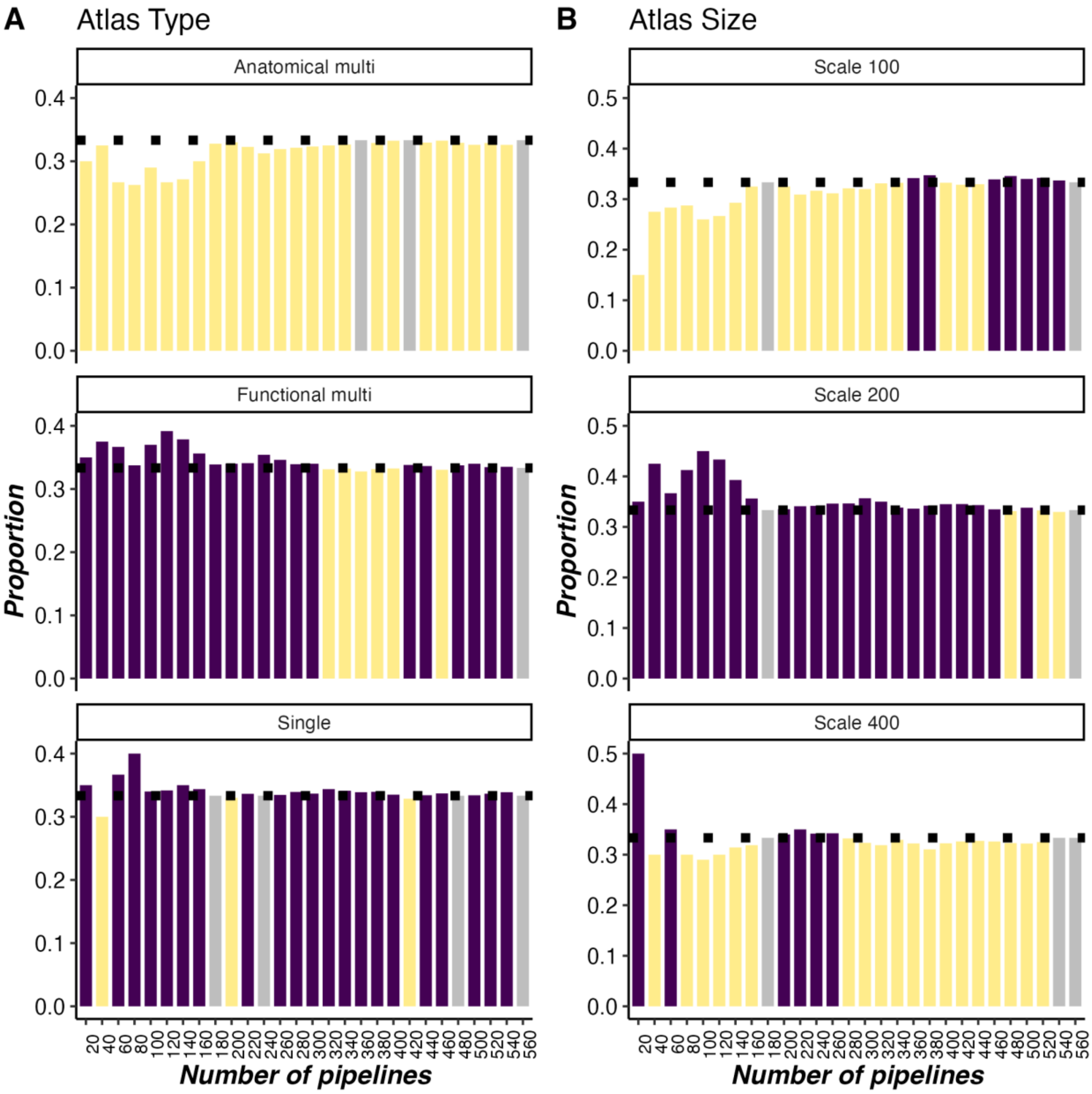


**Figure S3. Progression of pipeline choices as a function of node definition and average PDiv across all datasets.** (A) Divided by atlas type (anatomical multi-scale, functional multi-scale, or single-scale). (b) By atlas scale. With each subsequent bin, the next best 20 pipelines are added to calculate how many among this set of pipelines were constructed using each of the available options.

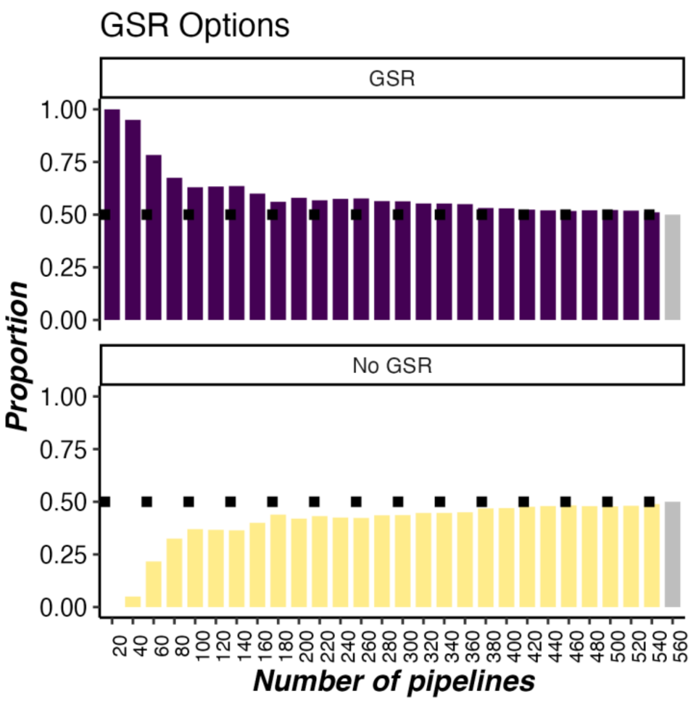


**Figure S4. Progression of pipeline choices as a function of GSR use and average PDiv across all datasets.** With each subsequent bin, the next best 20 pipelines are added to calculate how many among this set of pipelines were constructed using each of the available options.

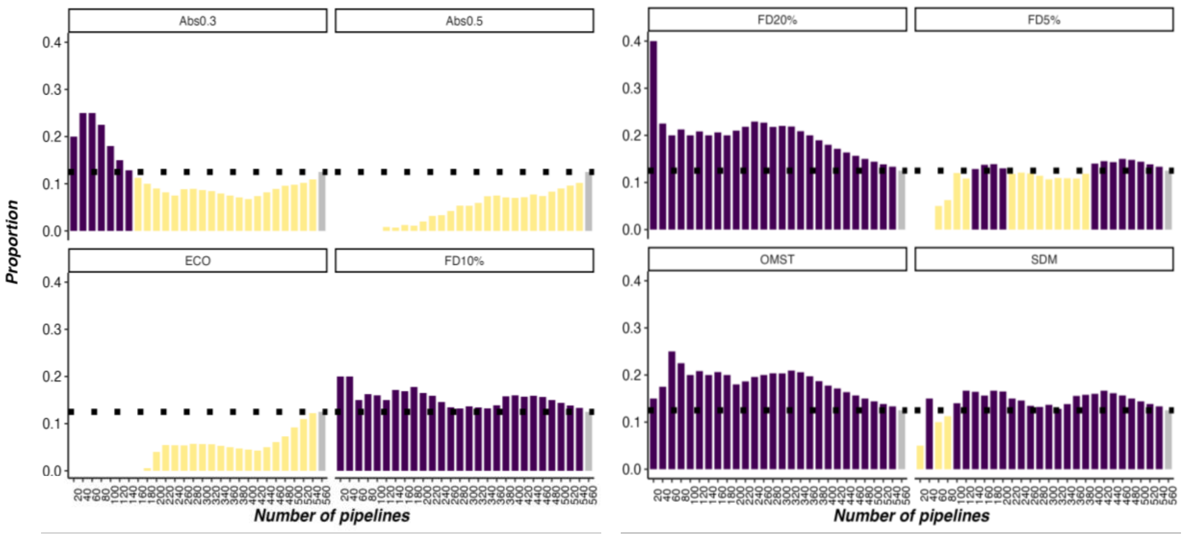


**Figure S5. Progression of pipeline choices as a function of filtering scheme and average PDiv across all datasets.** With each subsequent bin, the next best 20 pipelines are added to calculate how many among this set of pipelines were constructed using each of the available options.

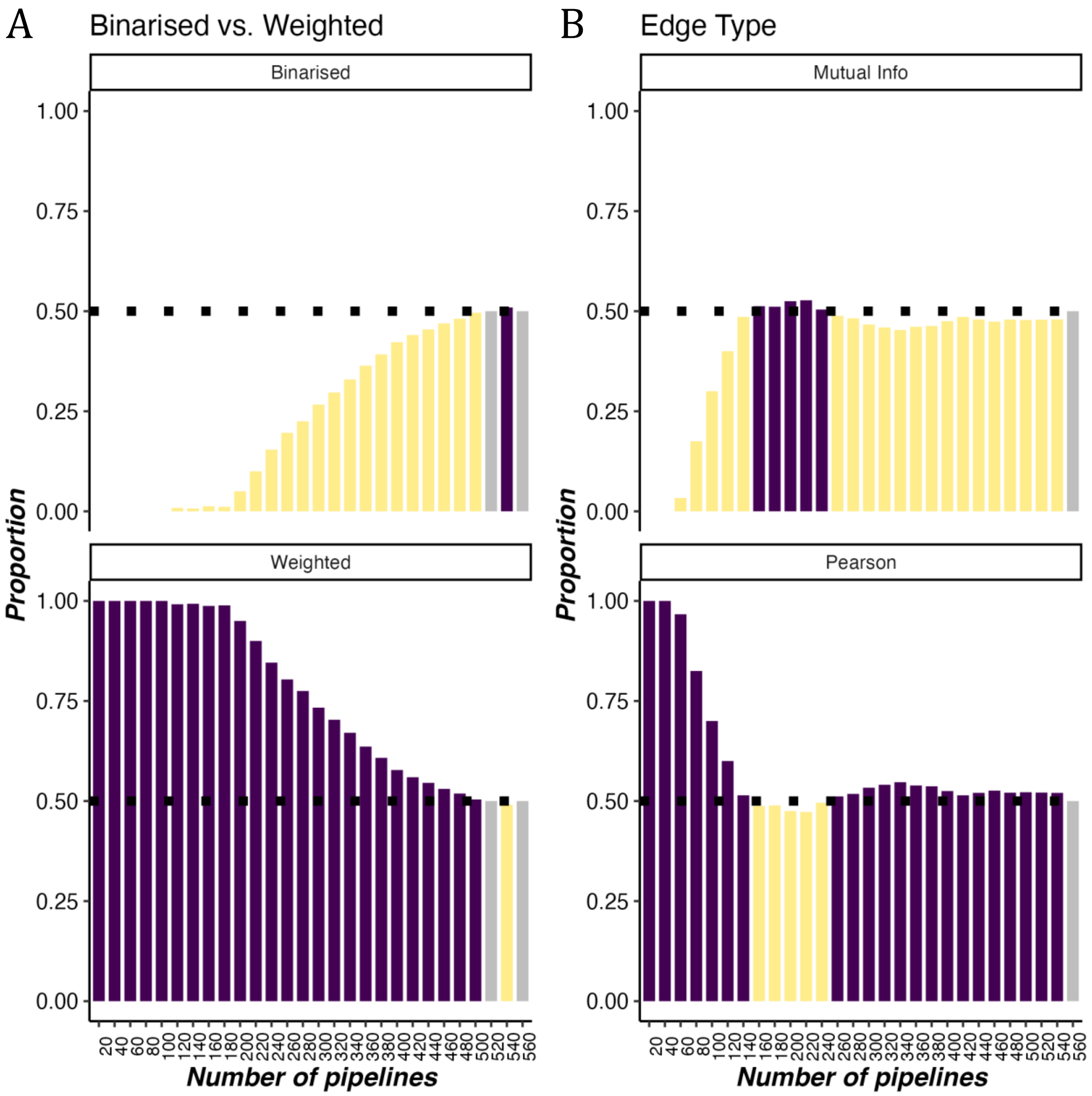


**Figure S6. Progression of pipeline choices as a function of edge construction and average PDiv across all datasets.** (A) Binary vs weighted edges. (B) Edges quantified in terms of mutual information or Pearson correlation. With each subsequent bin, the next best 20 pipelines are added to calculate how many among this set of pipelines were constructed using each of the available options.

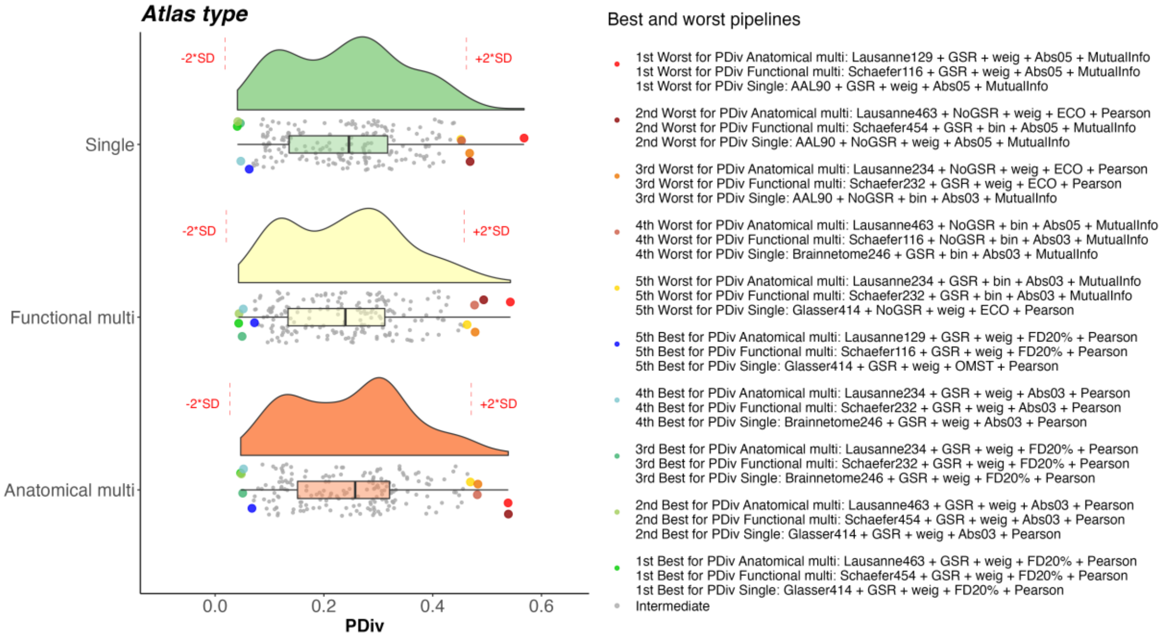


**Figure S7. Portrait divergence (PDiv) by atlas type – Cambridge test-retest dataset.** Box-plot center line, median; box limits, upper and lower quartiles; whiskers, 1.5x interquartile range.

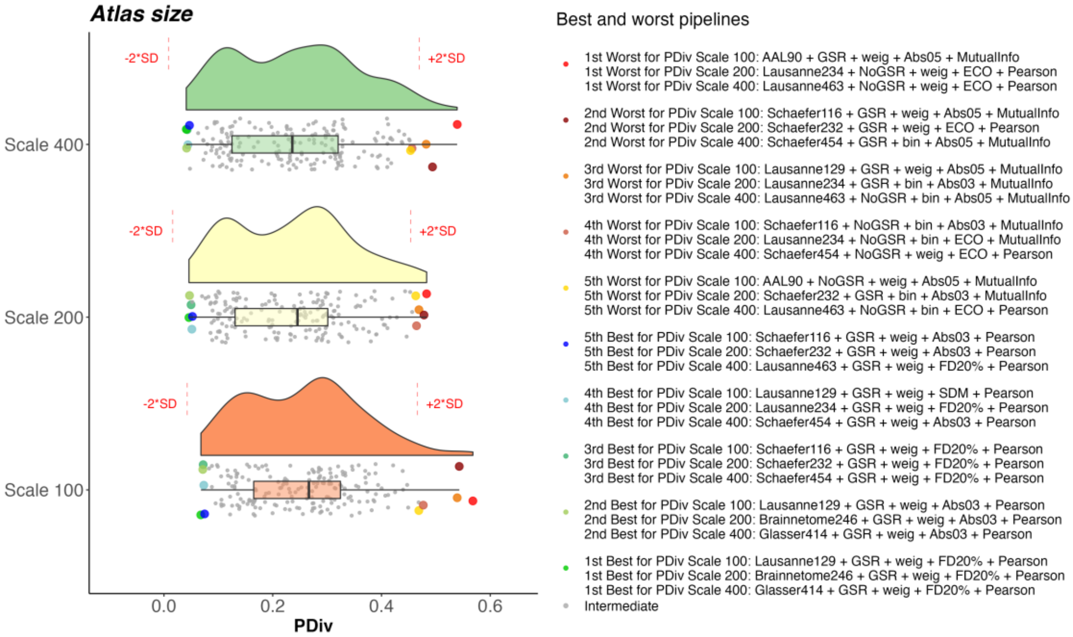


**Figure S8. Portrait divergence (PDiv) by atlas scale – Cambridge test-retest dataset.** Box-plot center line, median; box limits, upper and lower quartiles; whiskers, 1.5x interquartile range.

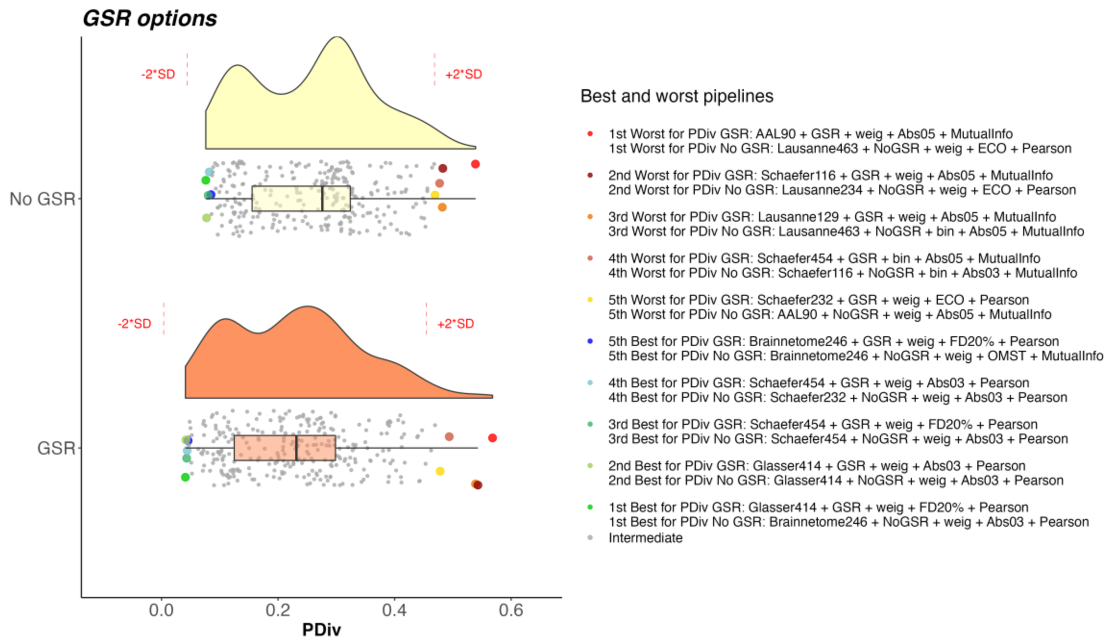


**Figure S9. Portrait divergence (PDiv) by GSR use – Cambridge test-retest dataset.** Box-plot center line, median; box limits, upper and lower quartiles; whiskers, 1.5x interquartile range.

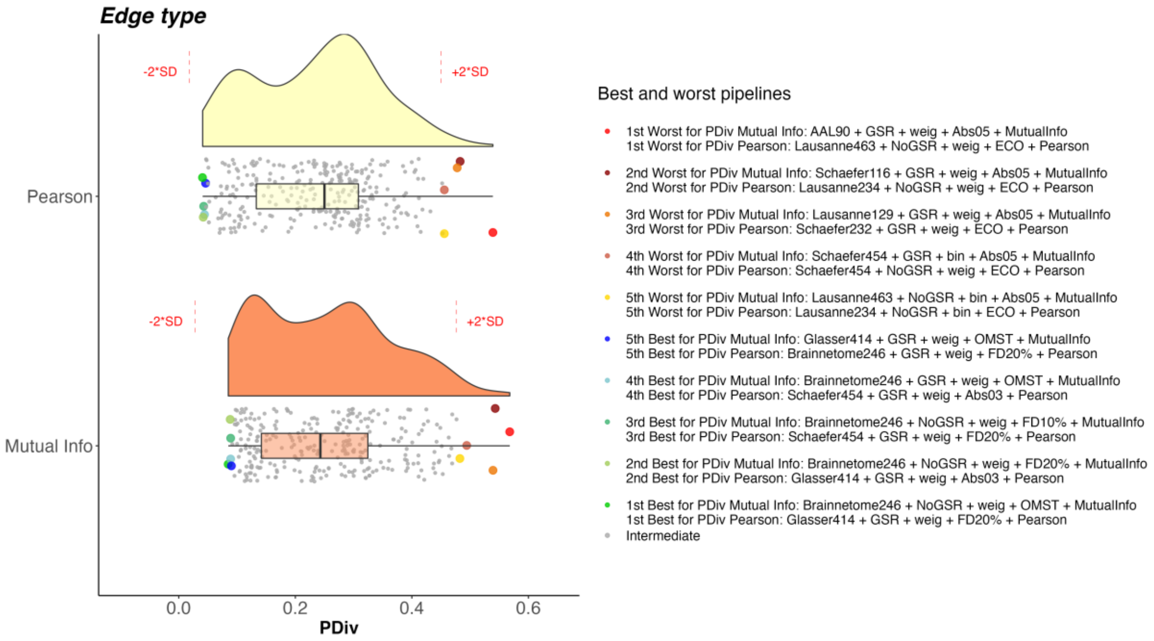


**Figure S10. Portrait divergence (PDiv) by edge quantification method type – Cambridge test-retest dataset.** Box-plot center line, median; box limits, upper and lower quartiles; whiskers, 1.5x interquartile range.

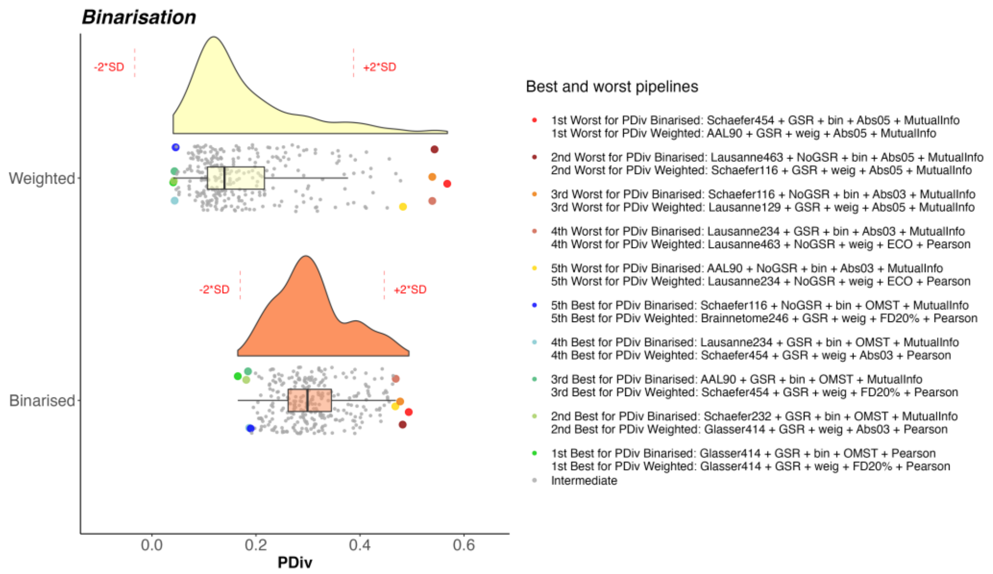


**Figure S11. Portrait divergence (PDiv) by binarisation choice – Cambridge test-retest dataset.** Box-plot center line, median; box limits, upper and lower quartiles; whiskers, 1.5x interquartile range.

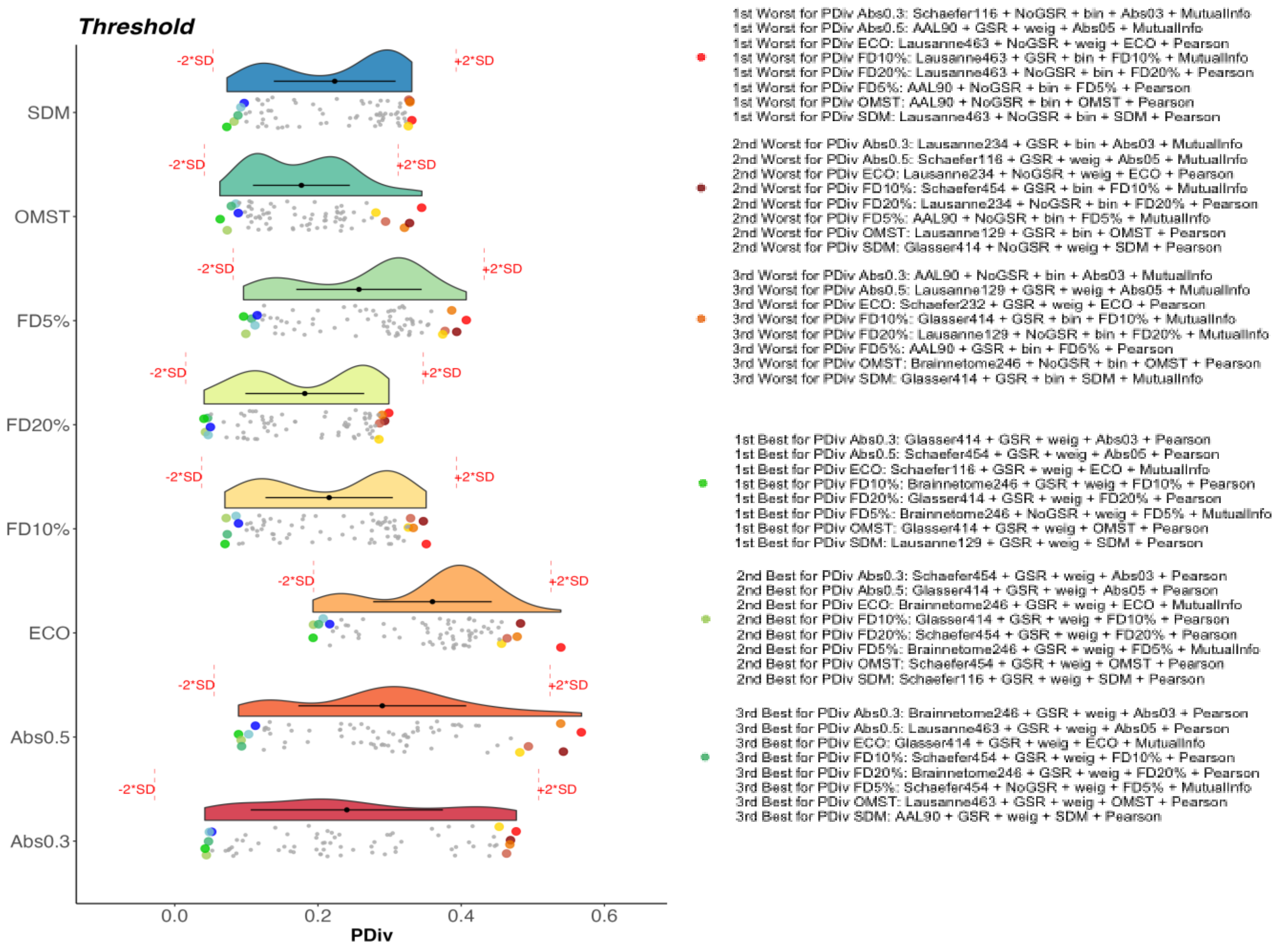


**Figure S12. Portrait divergence (PDiv) by edge filtering method – Cambridge test-retest dataset.** Box-plot center line, median; box limits, upper and lower quartiles; whiskers, 1.5x interquartile range.

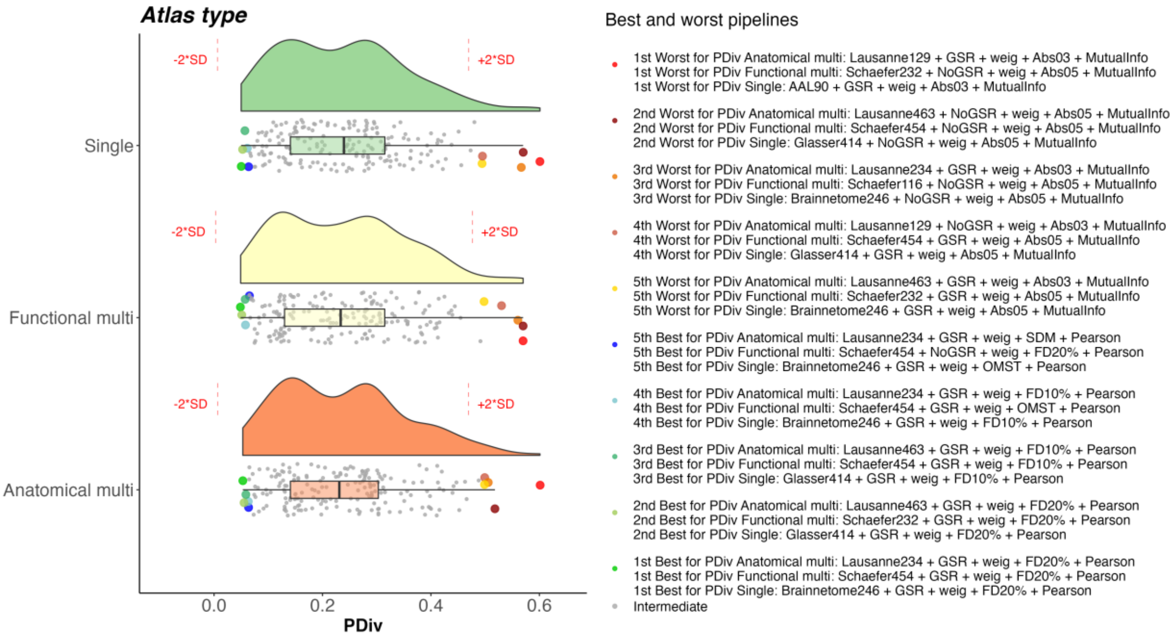


**Figure S13. Portrait divergence (PDiv) by atlas type – NYU short-term dataset.** Box-plot center line, median; box limits, upper and lower quartiles; whiskers, 1.5x interquartile range.

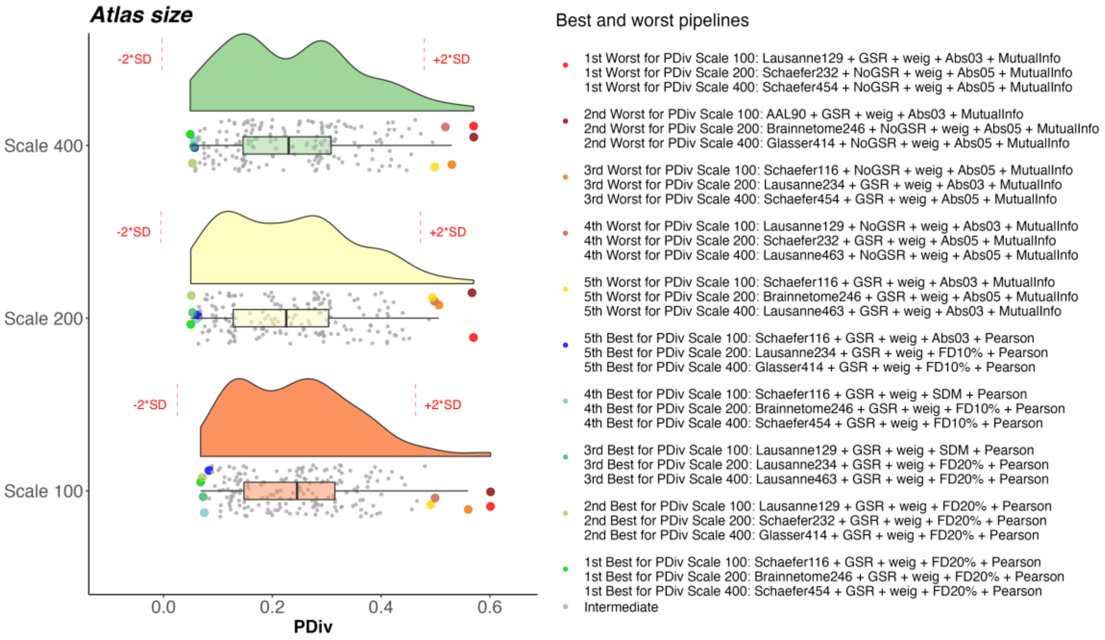


**Figure S14. Portrait divergence (PDiv) by atlas scale – NYU short-term dataset.** Box-plot center line, median; box limits, upper and lower quartiles; whiskers, 1.5x interquartile range.

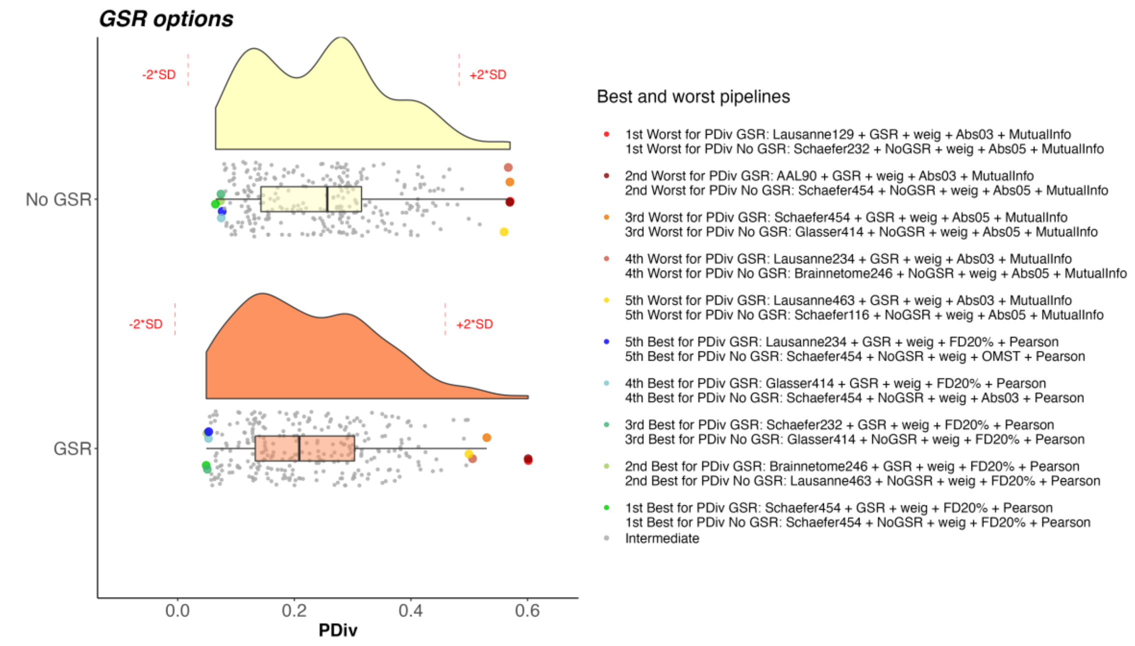


**Figure S15. Portrait divergence (PDiv) by GSR use – NYU short-term dataset.** Box-plot center line, median; box limits, upper and lower quartiles; whiskers, 1.5x interquartile range.

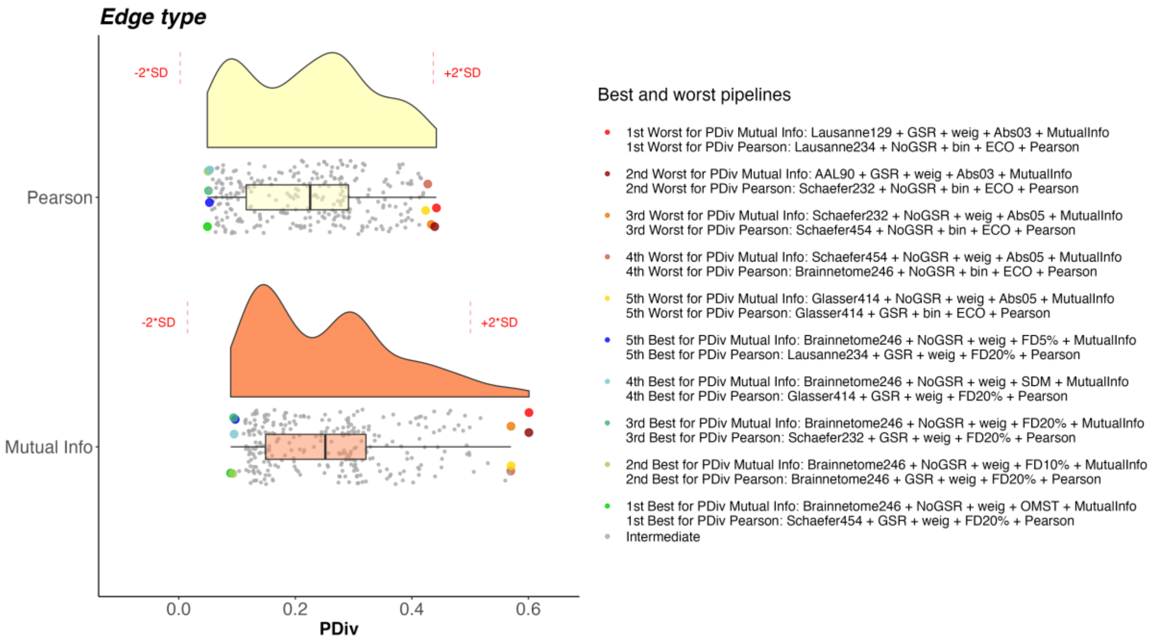


**Figure S16. Portrait divergence (PDiv) by edge quantification method type – NYU short-term dataset.** Box-plot center line, median; box limits, upper and lower quartiles; whiskers, 1.5x interquartile range.

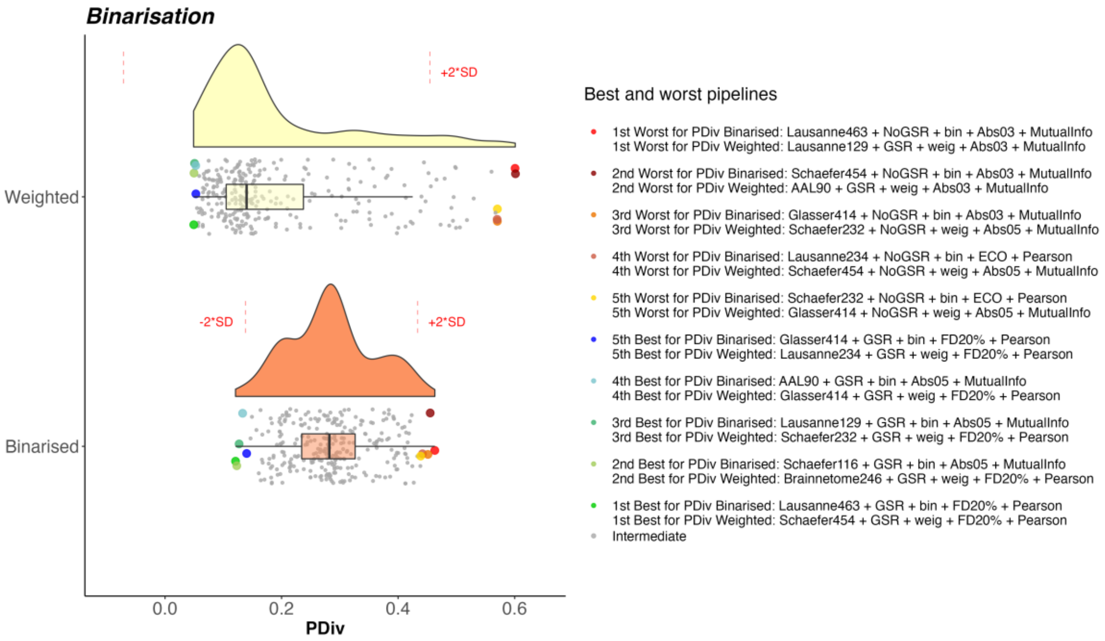


**Figure S17. Portrait divergence (PDiv) by binarisation choice – NYU short-term dataset.** Box-plot center line, median; box limits, upper and lower quartiles; whiskers, 1.5x interquartile range.

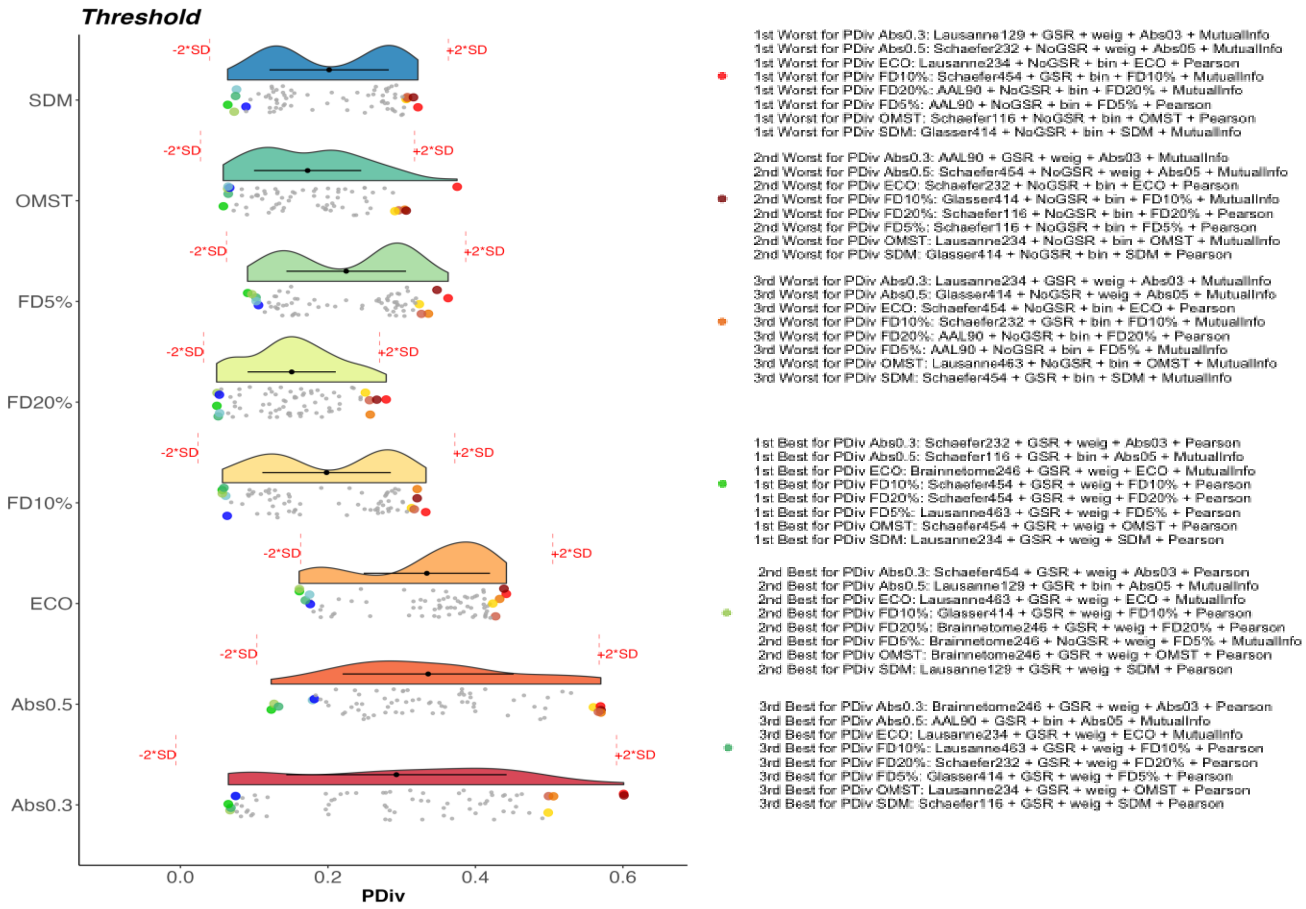


**Figure S18. Portrait divergence (PDiv) by edge filtering method – NYU short-term dataset.** Box-plot center line, median; box limits, upper and lower quartiles; whiskers, 1.5x interquartile range.

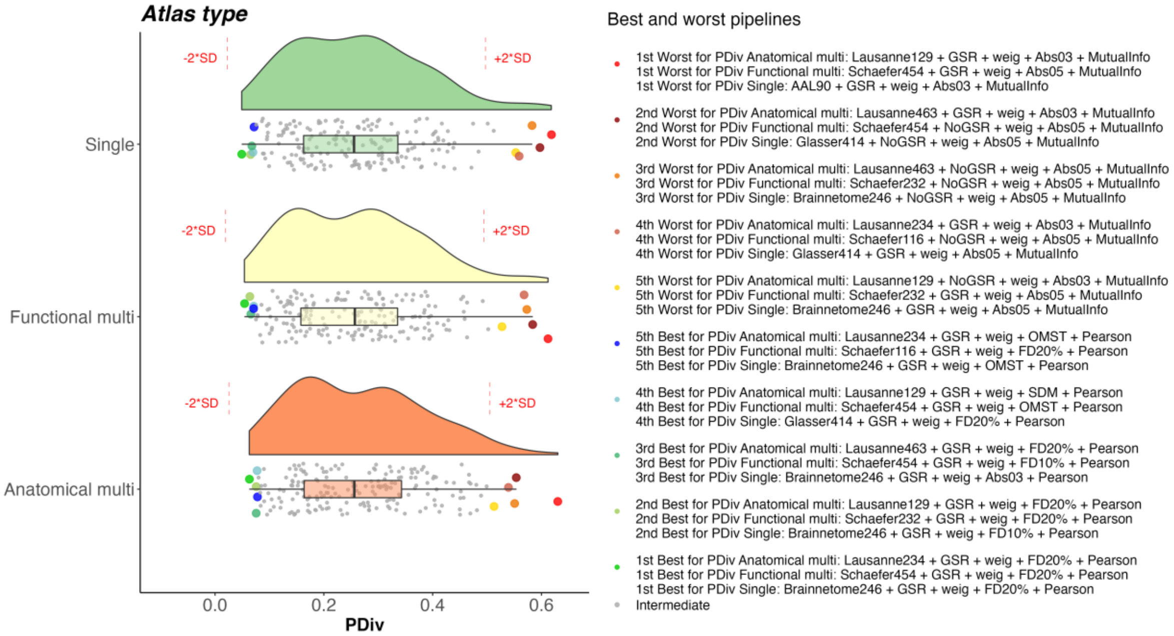


**Figure S19. Portrait divergence (PDiv) by atlas type – NYU long-term dataset.** Box-plot center line, median; box limits, upper and lower quartiles; whiskers, 1.5x interquartile range.

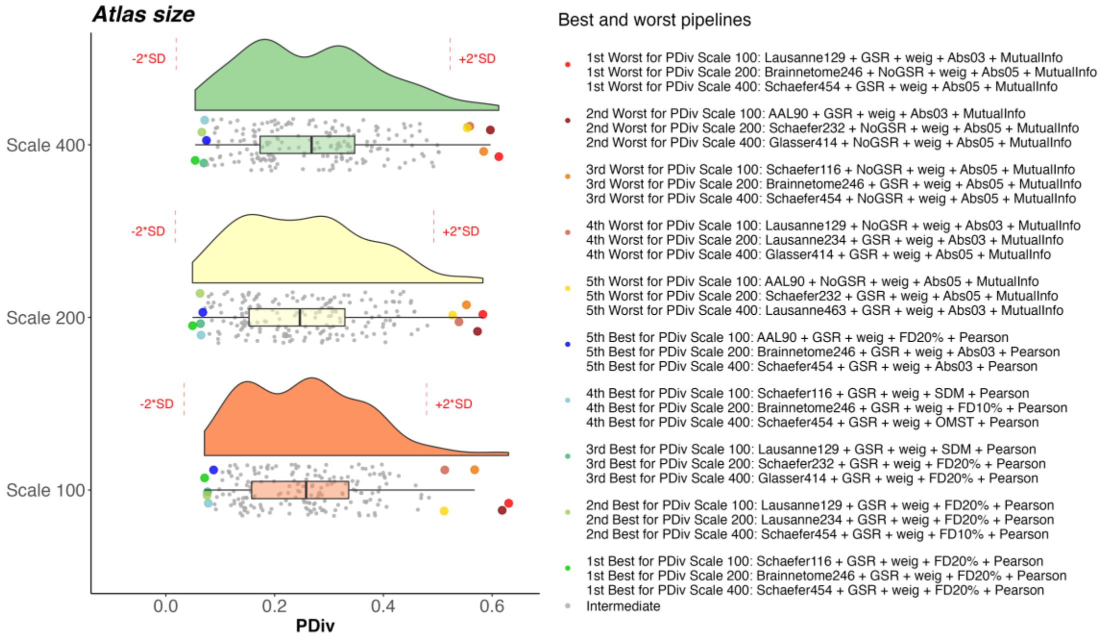


**Figure S20. Portrait divergence (PDiv) by atlas scale – NYU long-term dataset.** Box-plot center line, median; box limits, upper and lower quartiles; whiskers, 1.5x interquartile range.

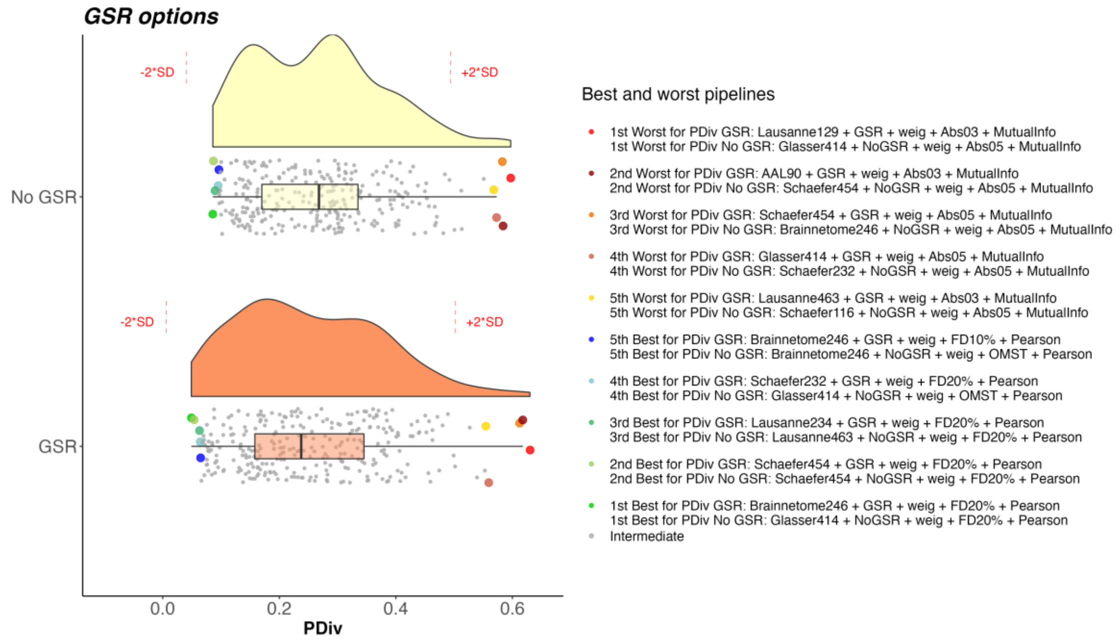


**Figure S21. Portrait divergence (PDiv) by GSR use – NYU long-term dataset.** Box-plot center line, median; box limits, upper and lower quartiles; whiskers, 1.5x interquartile range.

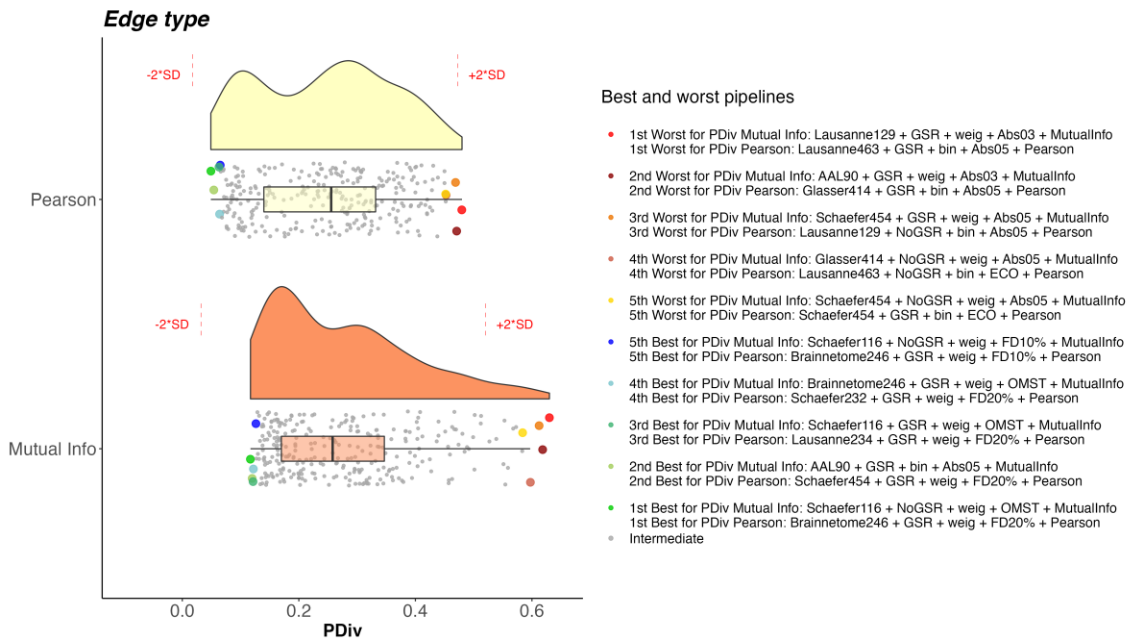


**Figure S22. Portrait divergence (PDiv) by edge quantification method type – NYU long-term dataset.** Box-plot center line, median; box limits, upper and lower quartiles; whiskers, 1.5x interquartile range.

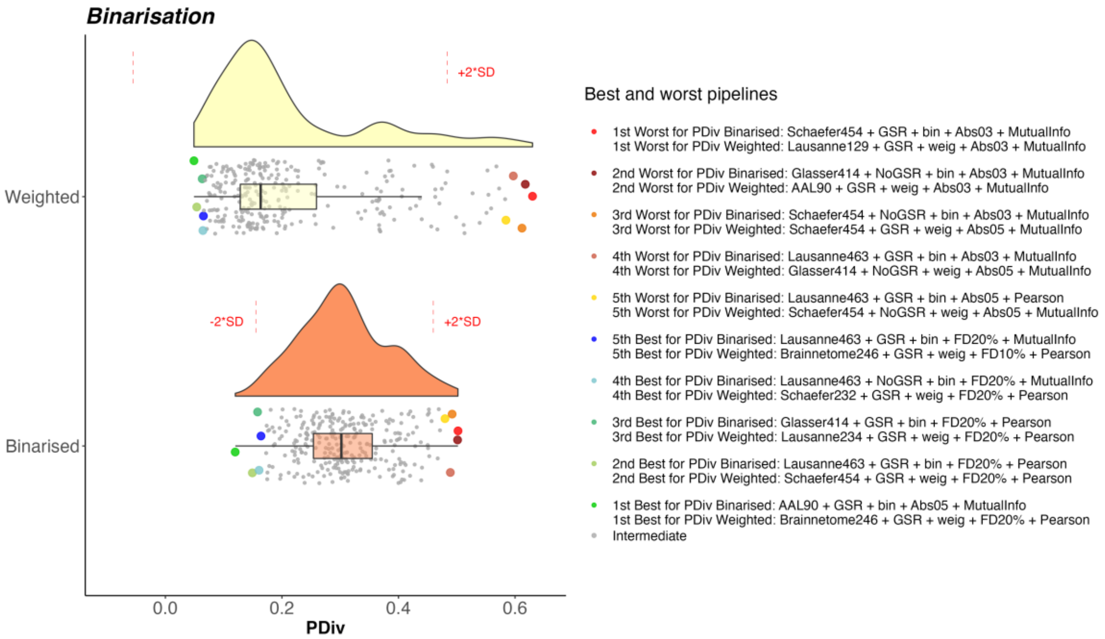


**Figure S23. Portrait divergence (PDiv) by binarisation choice – NYU long-term dataset.** Box-plot center line, median; box limits, upper and lower quartiles; whiskers, 1.5x interquartile range.

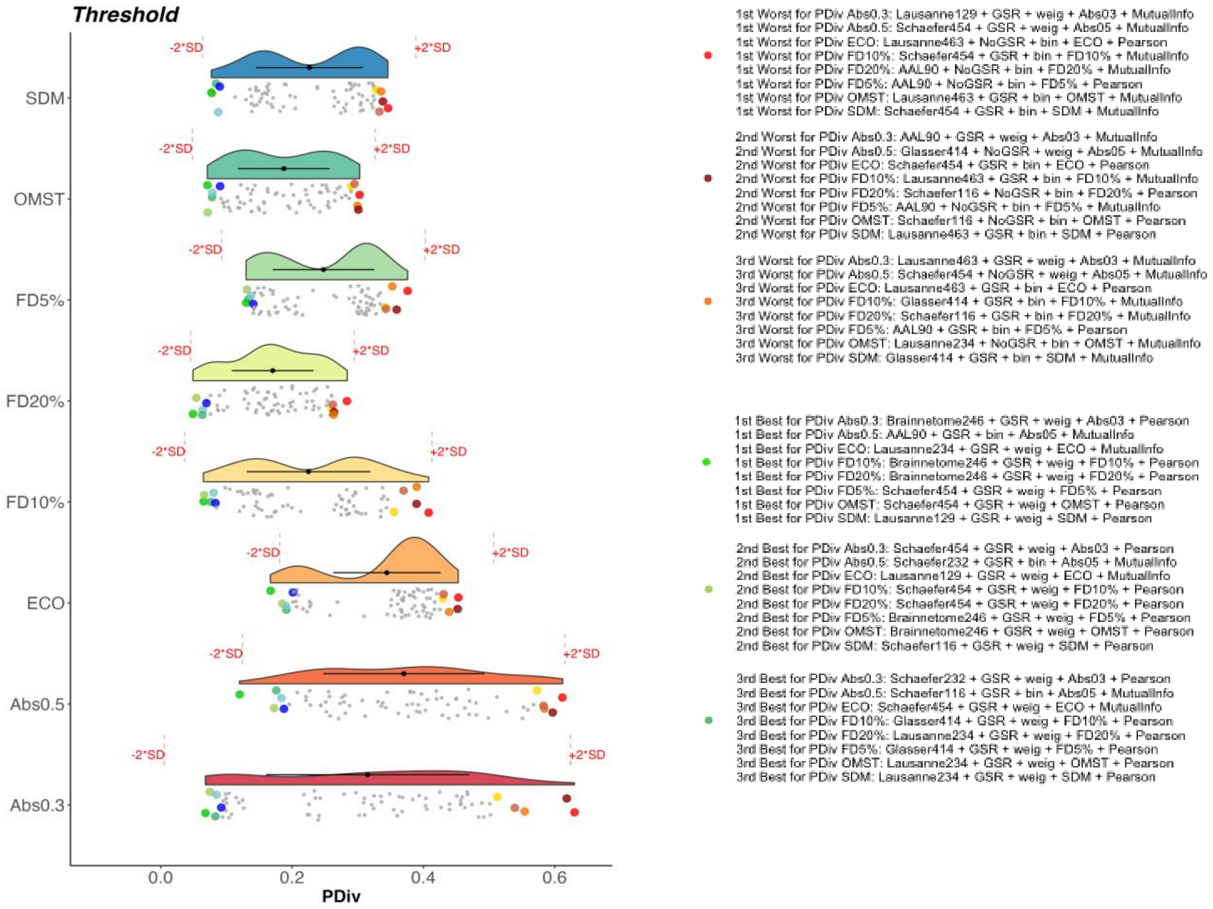


**Figure S24. Portrait divergence (PDiv) by edge filtering method – NYU long-term dataset.** Box-plot center line, median; box limits, upper and lower quartiles; whiskers, 1.5x interquartile range.

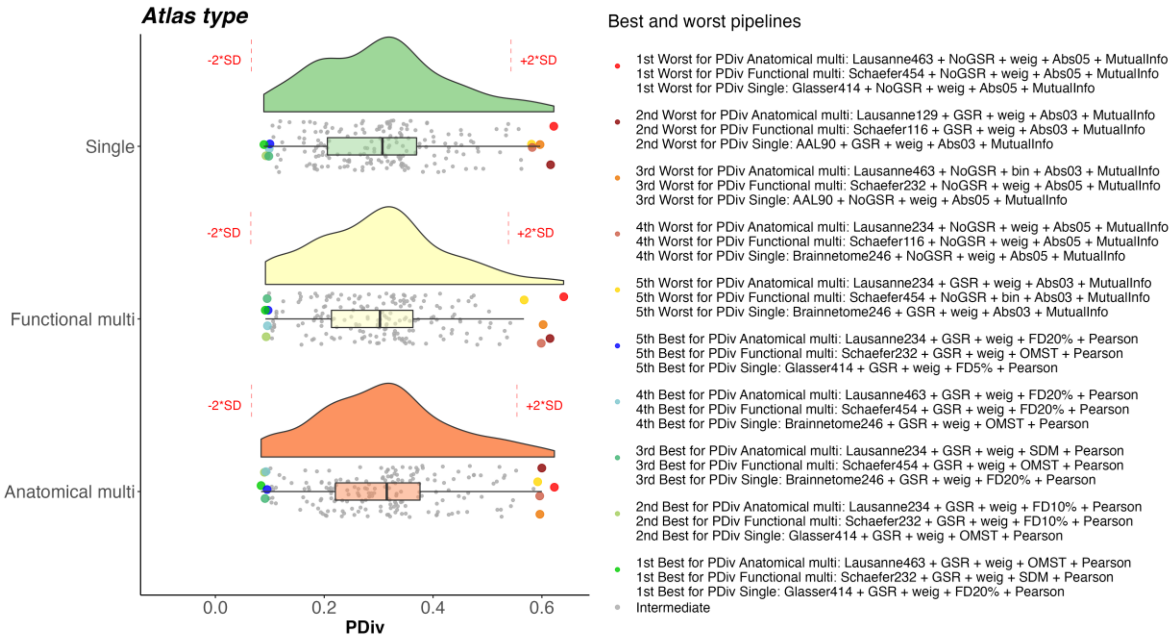


**Figure S25. Portrait divergence (PDiv) by atlas type – HCP test-retest dataset.** Box-plot center line, median; box limits, upper and lower quartiles; whiskers, 1.5x interquartile range.

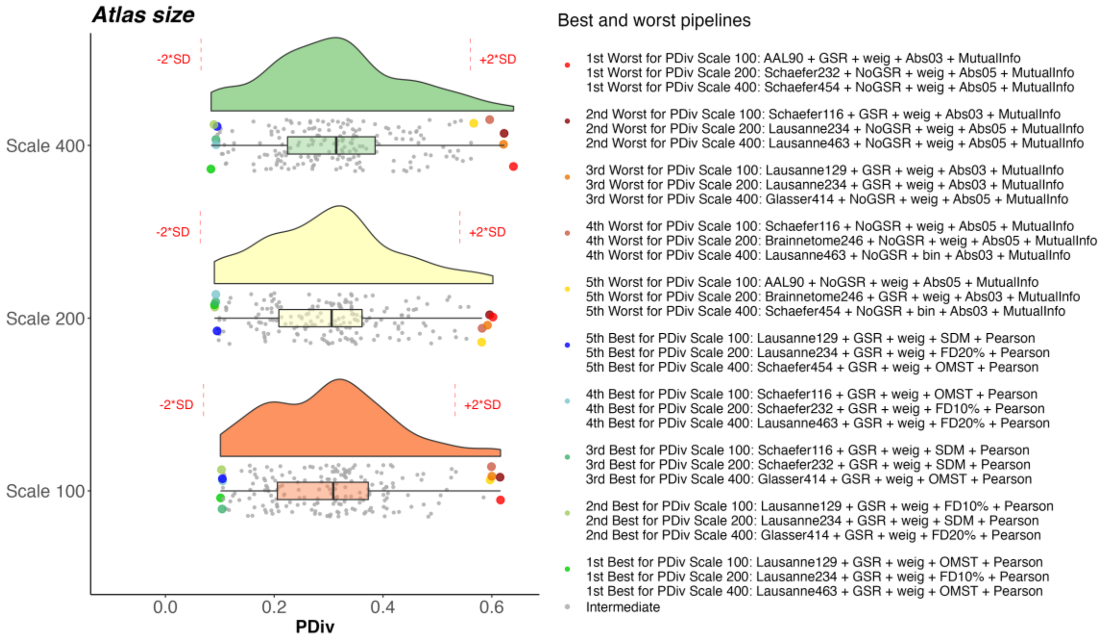


**Figure S26. Portrait divergence (PDiv) by atlas scale – HCP test-retest dataset.** Box-plot center line, median; box limits, upper and lower quartiles; whiskers, 1.5x interquartile range.

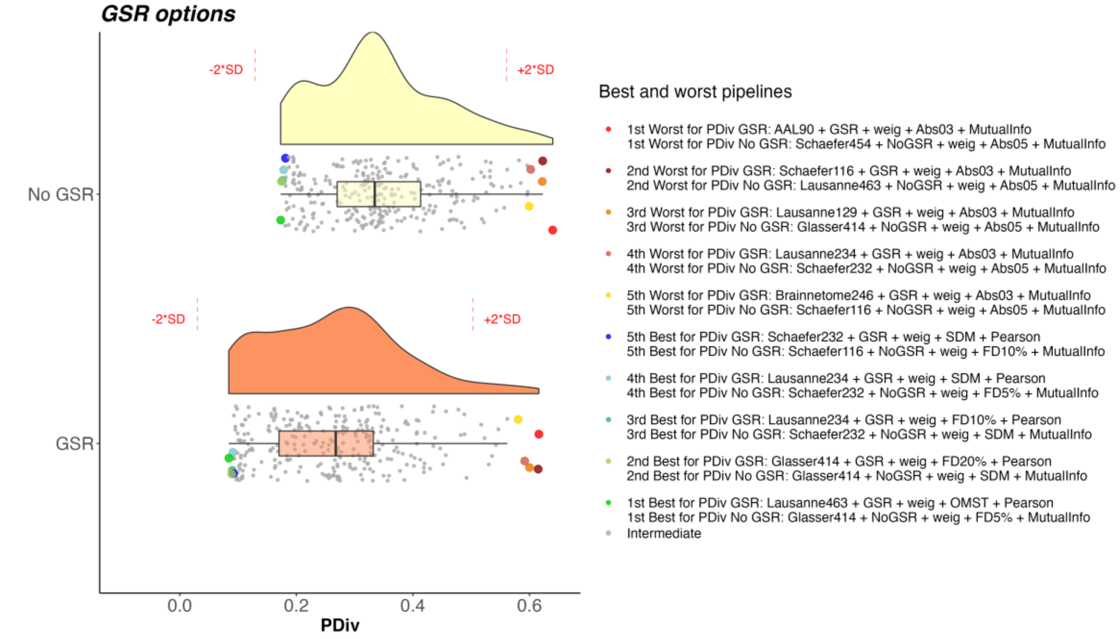


**Figure S27. Portrait divergence (PDiv) by GSR use – HCP test-retest dataset.** Box-plot center line, median; box limits, upper and lower quartiles; whiskers, 1.5x interquartile range.

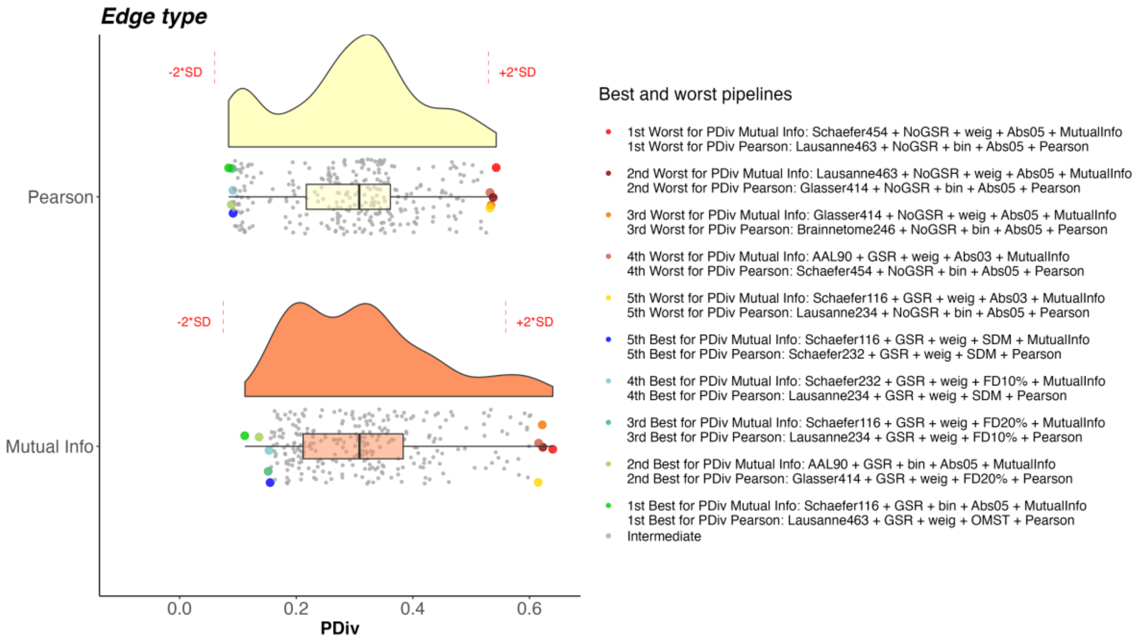


**Figure S28. Portrait divergence (PDiv) by edge quantification method type – HCP test-retest dataset.** Box-plot center line, median; box limits, upper and lower quartiles; whiskers, 1.5x interquartile range.

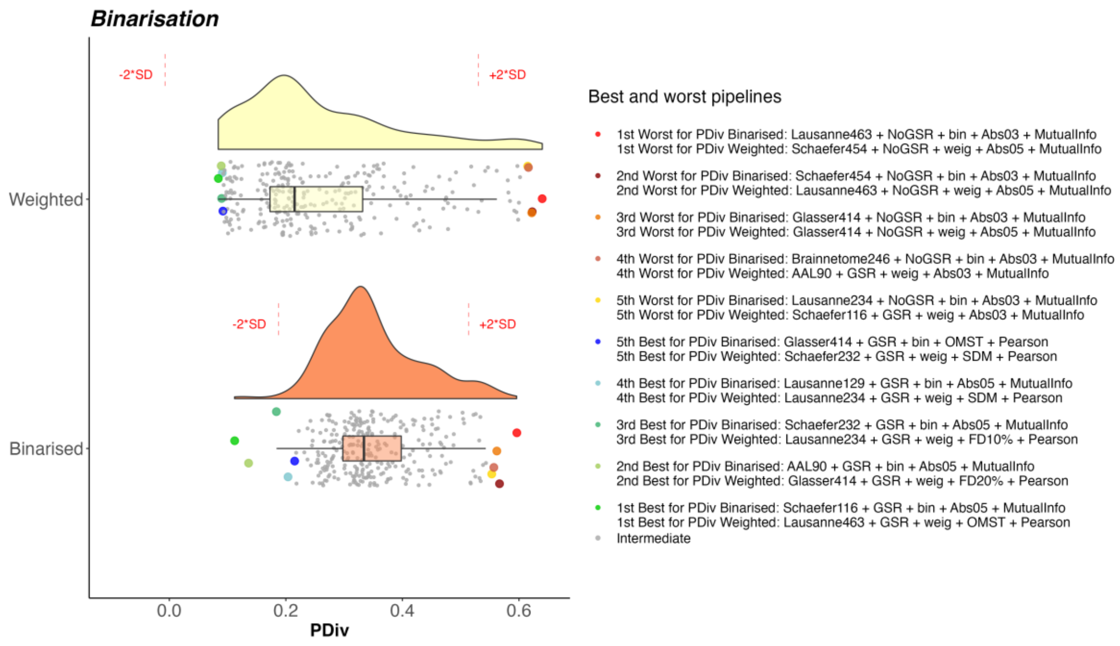


**Figure S29. Portrait divergence (PDiv) by binarisation choice – HCP test-retest dataset.** Box-plot center line, median; box limits, upper and lower quartiles; whiskers, 1.5x interquartile range.

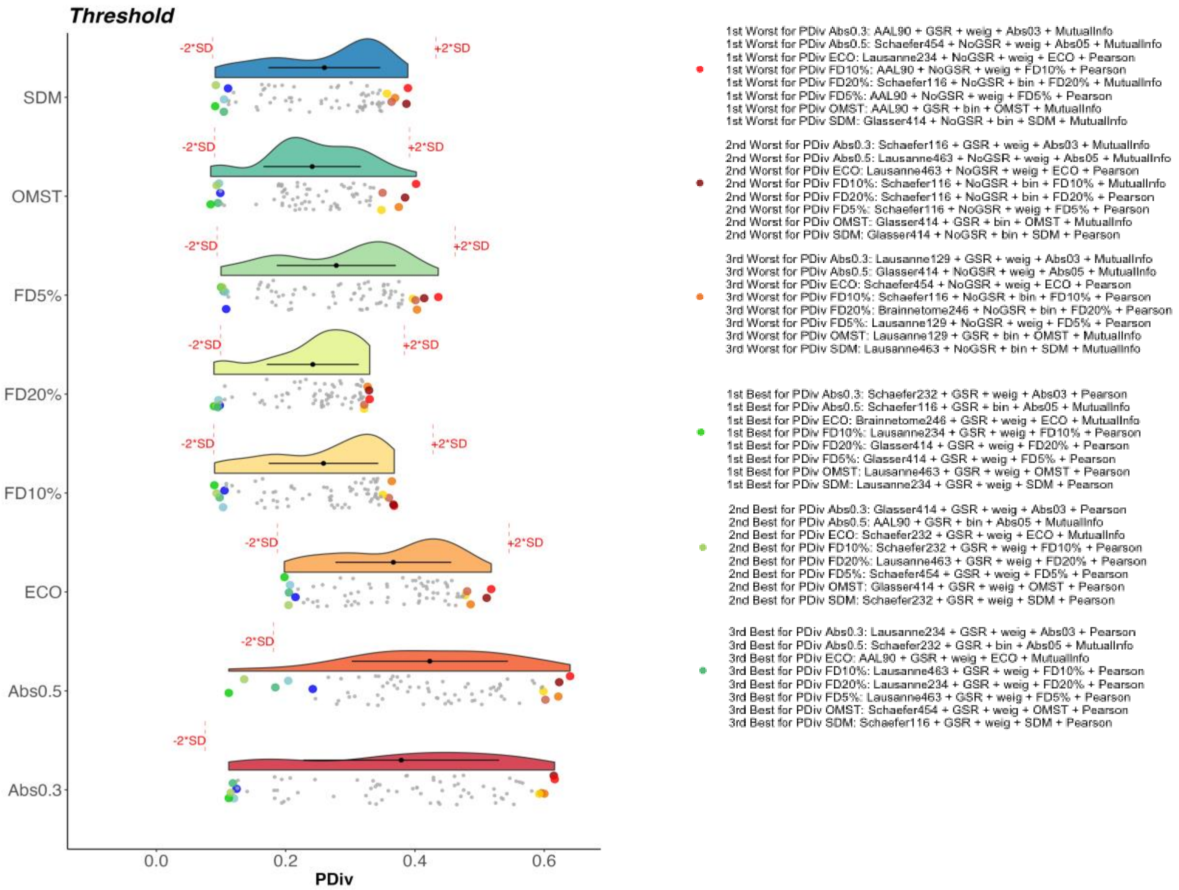


**Figure S30. Portrait divergence (PDiv) by edge filtering method – HCP test-retest dataset.** Box-plot center line, median; box limits, upper and lower quartiles; whiskers, 1.5x interquartile range.

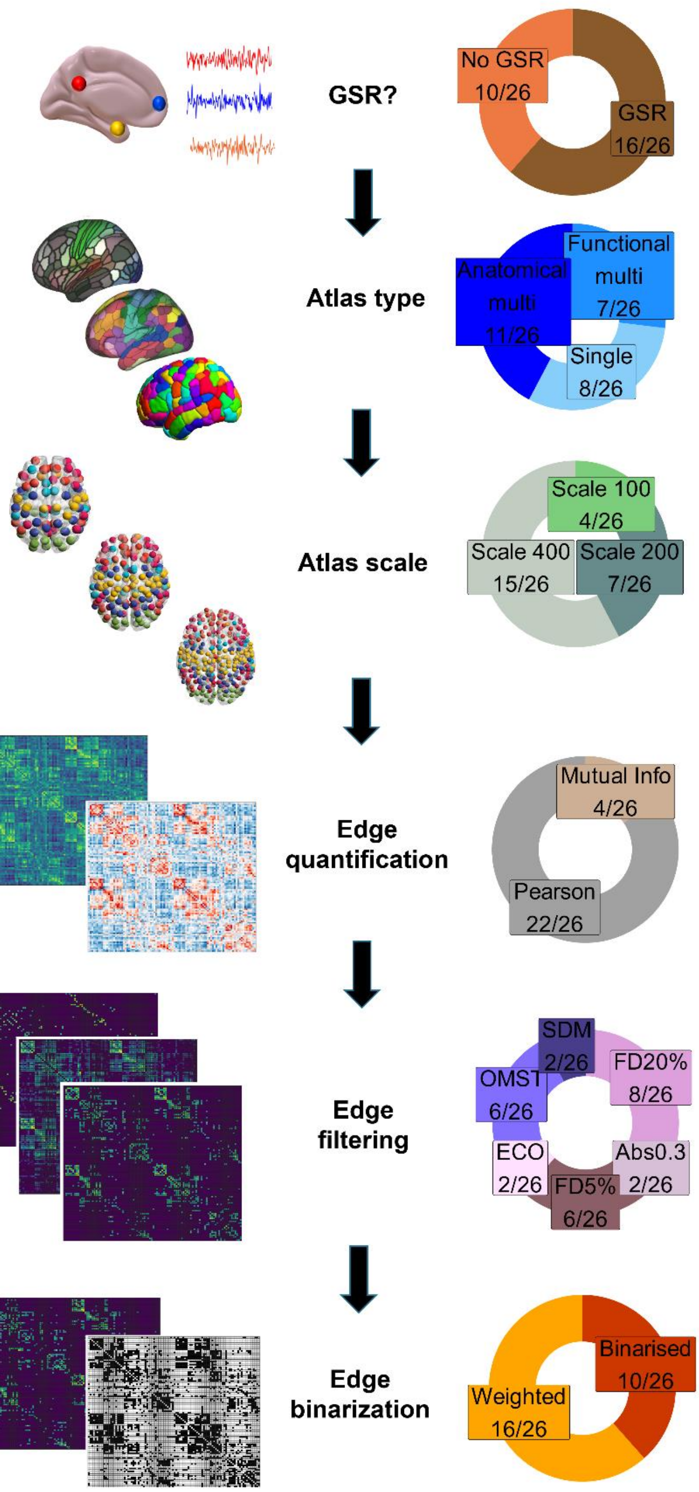


**Figure S31. Prevalence of specific network construction steps among the 26 optimal pipelines, when relaxing the PDiv criterion.** Pie charts demonstrate, for each network construction step, the proportion and absolute number of each option that is found among the optimal pipelines. **Abbreviations**. FD: fixed density. GSR: global signal regression. OMST: orthogonal minimal spanning tree. SDM: structural density.

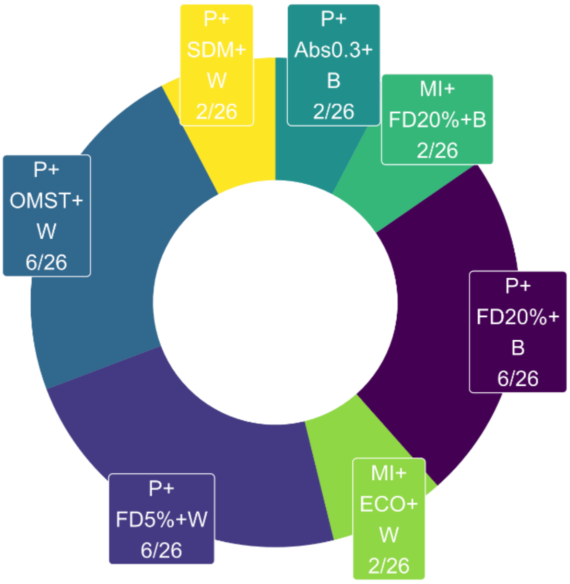


**Figure S32. Optimal edge processing combinations among the top 26 pipelines, when relaxing the PDiv criterion.** Pie chart displays the frequency of each combination of edge type definition, filtering, and binarisation among the 26 pipelines which fulfil all criteria for a suitable network construction pipeline. P, Pearson correlation; MI, mutual information; B, binary edges; W, weighted edges; FD5, 5% fixed density threshold; FD20, 20% fixed density threshold; ECO, efficiency-cost optimisation; OMST, orthogonal minimum spanning trees, Abs, absolute threshold; SDM, structural density matching.

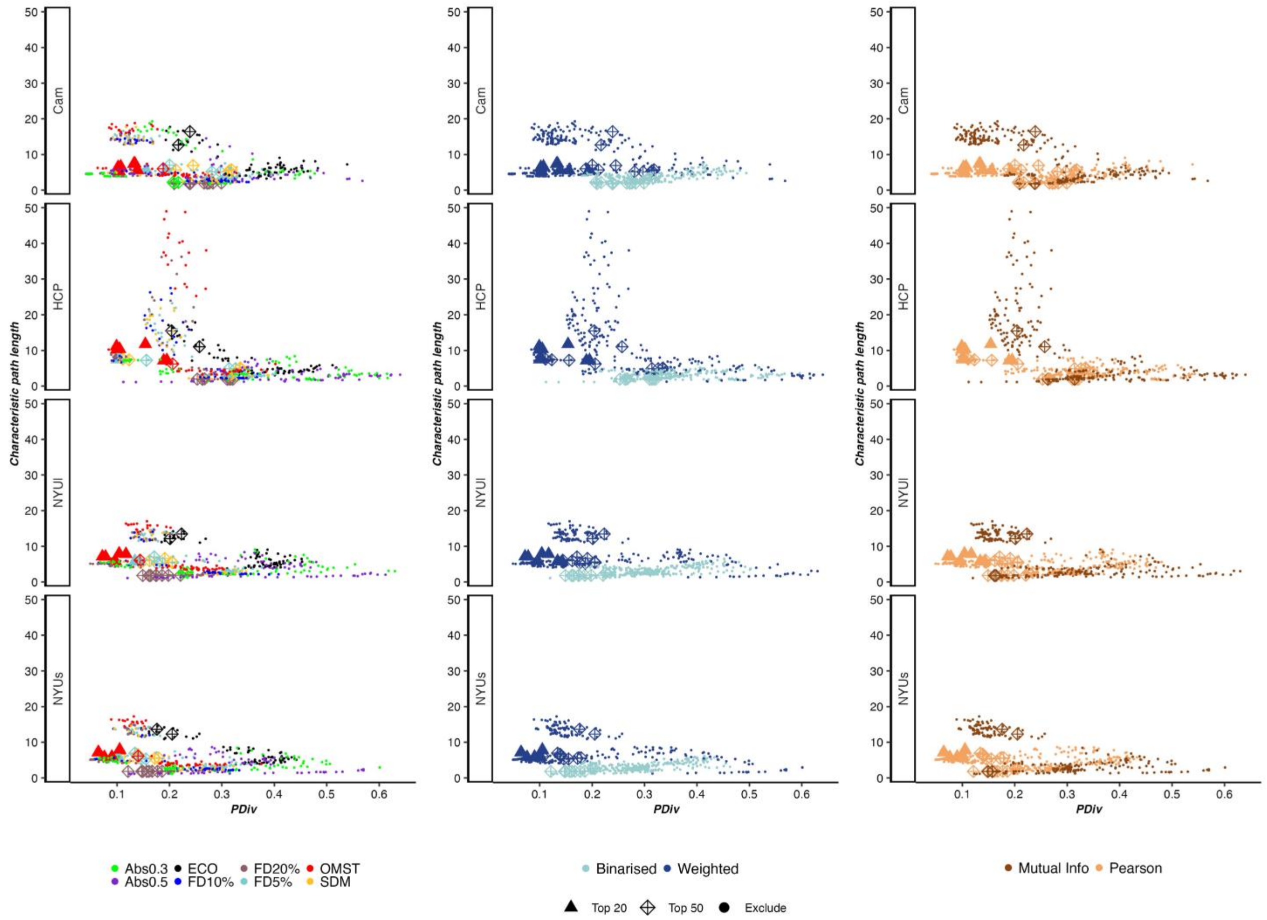


**Figure S33. Test-retest PDiv versus characteristic path length of the networks produced by each pipeline (averaged across all subjects), for each dataset, as a function of filtering scheme, edge binarisation, and edge type (Pearson correlation or mutual information).** Each data-point represents one pipeline; shape indicates optimality (optimal under stringent criteria, optimal under the relaxed PDiv criterion, or rejected).

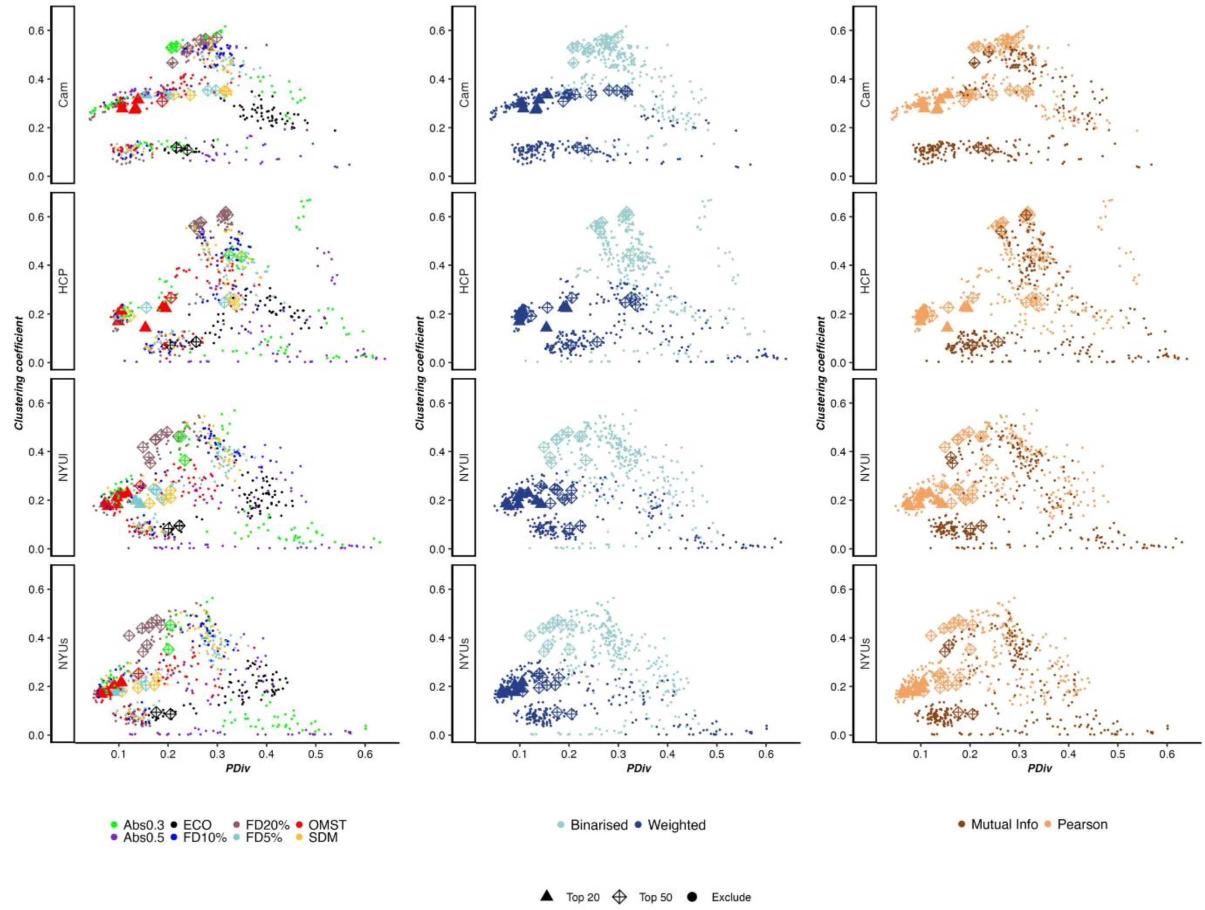


**Figure S34. Test-retest PDiv versus mean clustering coefficient of the networks produced by each pipeline (averaged across all subjects), for each dataset, as a function of filtering scheme, edge binarisation, and edge type (Pearson correlation or mutual information).** Each data-point represents one pipeline; shape indicates optimality (optimal under stringent criteria, optimal under the relaxed PDiv criterion, or rejected).

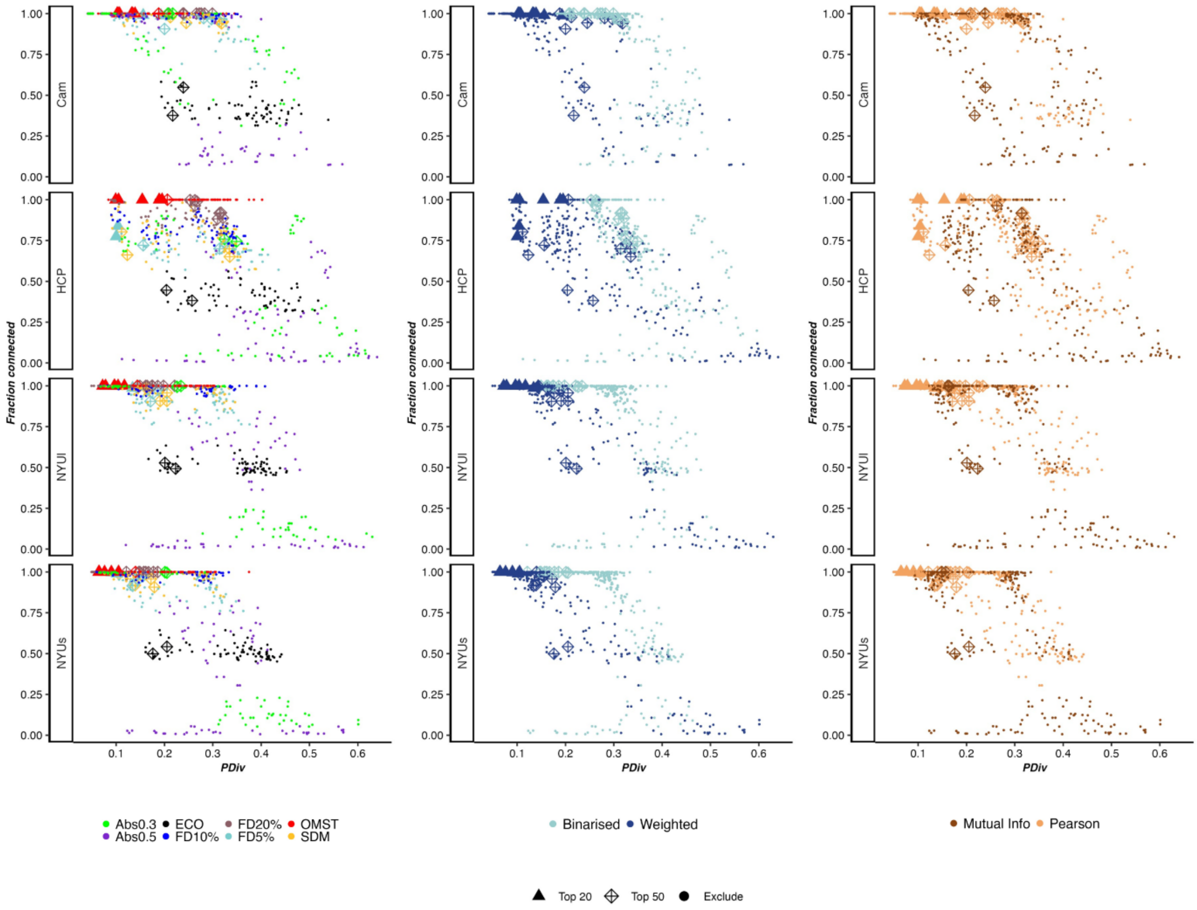


**Figure S35. Test-retest PDiv versus the size of the largest connected component (as a fraction of total number of nodes) of the networks produced by each pipeline (averaged across all subjects), for each dataset, as a function of filtering scheme, edge binarisation, and edge type (Pearson correlation or mutual information).** Each data-point represents one pipeline; shape indicates optimality (optimal under stringent criteria, optimal under the relaxed PDiv criterion, or rejected).
